# Sequence and structural rules define PY-NLS–dependent nuclear import by TRANSPORTIN 1 in plants

**DOI:** 10.1101/2025.07.20.665762

**Authors:** Ankush Ashok Saddhe, Přemysl Pejchar, Martin Potocky

## Abstract

Nuclear localization signals (NLSs) direct proteins to the nucleus, but whether the non-classical proline–tyrosine NLS (PY-NLS), characterized mainly in animals and fungi, functions in plants is unknown. Here we identify a basic PY-NLS pathway in tobacco and Arabidopsis and show that TRANSPORTIN 1 (TRN1) is its receptor. TRN1 binds PY-NLS cargoes *in vitro*, whereas loss of TRN1 causes their cytoplasmic accumulation. Mutational, structural and quantitative localization analyses of motifs from the jasmonate signaling repressor JAZ1 and a PLA_2_-like protein reveal an extended linker between binding epitopes and a short α-helix preceding the conserved PY dipeptide as features that distinguish plant PY-NLSs from established eukaryotic models. A validated prediction pipeline identifies 179 Arabidopsis proteins containing candidate PY-NLSs. Together, our findings establish TRN1-mediated PY-NLS recognition as a previously unrecognized plant nuclear import pathway and uncover a plant-specific signal architecture with broad implications for the nuclear control of gene expression, RNA metabolism and development.

## INTRODUCTION

Nucleocytoplasmic transport is a highly regulated process that moves proteins and RNAs between the cytoplasm and nucleus through the nuclear pore complex (NPC). Nuclear import receptors recognize nuclear localization signals (NLSs) on cargo proteins and orchestrate the import through the Ran GTPase cycle (McPherson et al., 2015; Tamura and Hara-Nishimura, 2014; Soniat and Chook, 2015). Among these signals, classical NLSs (cNLSs) have been the most extensively studied. They usually consist of one or two clusters of basic residues and are categorized as monopartite or bipartite motifs, with lysine-rich stretches as a hallmark feature (Kosugi et al., 2009; Karasev et al., 2022; Lu et al., 2021). In plants, several cNLSs have been experimentally characterized (reviewed in Muñoz-Díaz and Sáez-Vásquez (2022)), including signals that mediate the nuclear envelope association of ER proteins (Groves et al., 2019) and microbial effector targeting during host colonization (Tehrani and Mitra, 2023). Monopartite and bipartite cNLSs mainly engage importin-α and importin-β receptor machinery to cross the nuclear membrane barrier (McPherson et al., 2015).

In addition to the classical pathway, several classes of non-classical NLSs (ncNLSs) that diverge from canonical cNLS features have been reported, comprising a diverse and still incompletely defined group, including spatial epitope NLSs, cryptic motifs, and modular elements such as the proline–tyrosine NLS (PY-NLS) (reviewed in Lu et al. (2021)). PY-NLS motifs have been studied in detail in mammals and fungi, where they are recognized by the nuclear import receptor karyopherin β2 (Kapβ2, also known as transportin 1, TNPO1; importin β2, IMB2) (Lee et al., 2006; McPherson et al., 2015; Bourgeois et al., 2020; Wing et al., 2022). Based on structural and biochemical studies in yeast/animal cells, PY-NLS motifs were proposed to possess an overall disordered conformation with three essential epitopes: an N-terminal hydrophobic or basic region (epitope 1) loosely spaced from a C-terminal region that includes a conserved R/K/H residue (epitope 2) spaced 2-5 amino acids from the PY dipeptide (epitope 3). These features were first described in human hnRNP A1 (Lee et al., 2006), and subsequent studies extended the definition to other proteins in yeast and human cells (Lange et al., 2008). Interestingly, in the soybean pathogen *Phytophthora sojae*, PY-NLS motifs function poorly compared with cNLSs, suggesting possible lineage-specific differences (Fang et al., 2017). Despite their importance in animal and yeast systems, PY-NLS motifs have not been functionally characterized in plants. Ziemienowicz et al. (2003) performed the initial characterization of Arabidopsis TRANSPORTIN 1 (TRN1) as a plant karyopherin β2 ortholog, and Yan et al. (2016) identified 23 putative cargoes in a yeast two-hybrid screen for interactors of Arabidopsis TRN1, including proteins with and without predicted PY-NLS motifs, but neither work resolved functional motif rules or directly test PY-NLS relevance. To date, no comprehensive study has established whether PY-NLS motifs function in plant cells or whether they follow the same sequence and structural principles as their fungal and animal counterparts.

Here, we present the first functional and systematic characterization of PY-NLS motifs in plants, using *Arabidopsis thaliana*, *Nicotiana benthamiana*, and *Nicotiana tabacum* as model systems. By combining microscopy, molecular biology, biochemical assays, genetics, and structural bioinformatics, we demonstrate that PY-NLS motifs are essential for the nuclear import of the jasmonate signaling regulator JAZ1 and a previously uncharacterized phospholipase A_2_-like (PLA_2_-like) protein. Functional plant PY-NLS motifs show sequence and structural features distinct from those in yeast and animals, including an often-present N-terminal α-helix and a longer linker between epitopes 1 and 2. Applying updated sequence and structural consensus rules, we designed an *in silico* pipeline, which predicted PY-NLS-containing proteins in Arabidopsis and confirmed the validity of these rules for several additional targets. Together, our data define a plant-specific PY-NLS architecture that depends on the TRN1-mediated nuclear import pathway.

## RESULTS

### Plant cells contain a functional non-classical nuclear import system dependent on PY-NLS motifs

During our comprehensive *in silico* analysis of plant phospholipase A_2_ genes (PLA_2_), we uncovered a previously uncharacterized family that we termed PLA_2_-like (Saddhe and Potocký, 2023). PLA_2_-like localizes strongly to the nucleus in both sporophyte and male gametophyte cells (Fig. 1A-C, Supplementary Figure 1A), but lacks any predictable canonical nuclear localization signal. During the initial sequence characterization of PLA_2_-like, we identified a motif that partially resembles the basic PY-NLS consensus known from animal and yeast cells (Fig. 1A). We could not find any report describing the presence of a functional PY-NLS system in plants, but we noticed that the members of the JAZ family, well-known nuclear-localized proteins implicated in jasmonate signaling and lacking any cNLS sequence, had been hypothesized to contain PY-NLS motifs at their C-termini (Pauwels and Goossens, 2011). We therefore investigated the importance of putative plant PY-NLS motifs in the nuclear import of PLA_2_-like and JAZ1.

**Figure 1.**
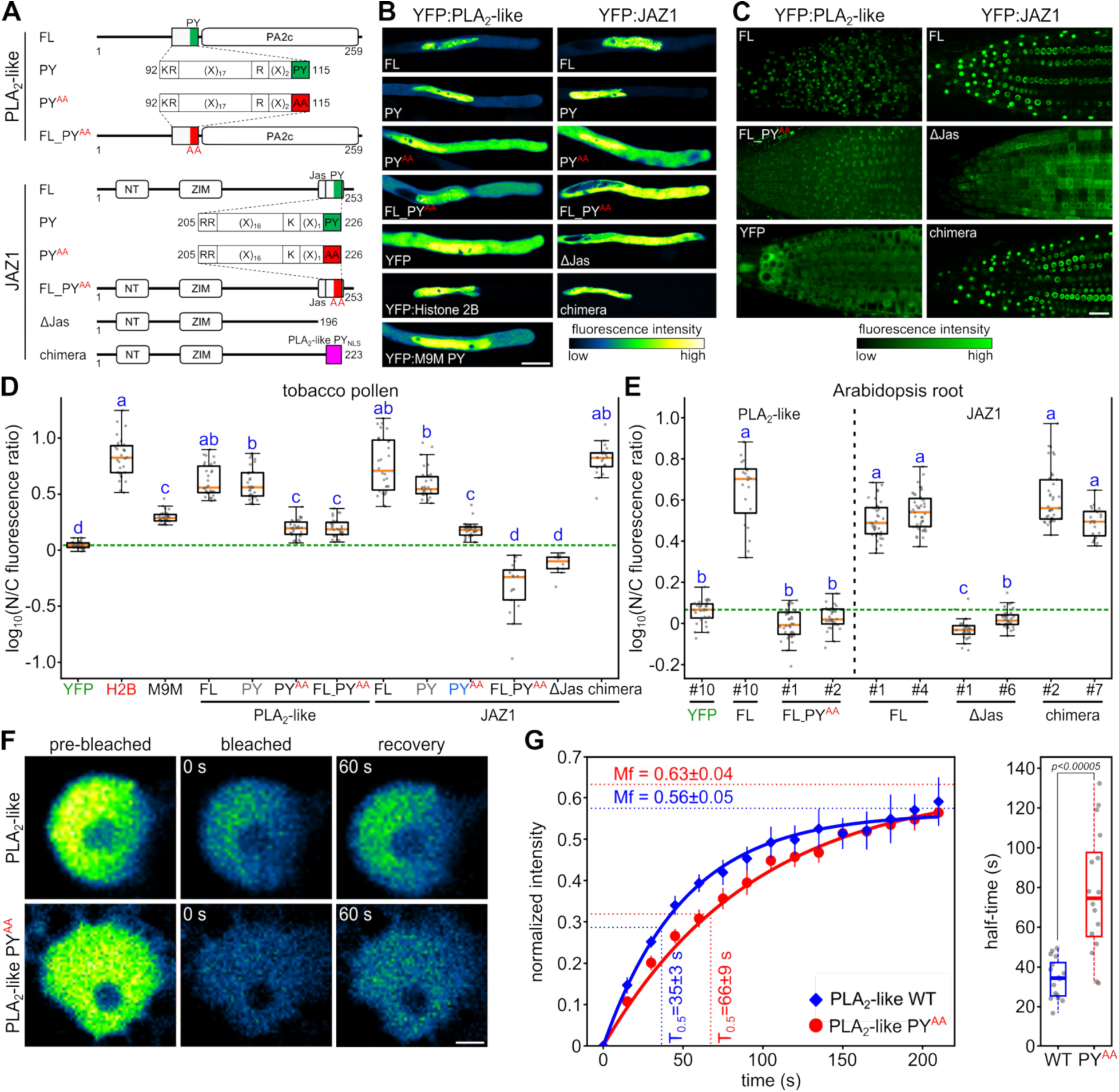
Identification and validation of non-classical basic PY-NLS motifs and nucleocytoplasmic dynamics of PY-NLS in plants. **A)** Schematic representation of PLA_2_-like and JAZ1 full-length (FL) and its PY-to-AA mutant (FL_PY^AA^), motif-only (PY) and its PY-to-AA mutant PY^AA^, ΔJas JAZ1 variant, and chimeric JAZ1 construct (using the PLA_2_-like PY-NLS fused to JAZ1 lacking the Jas domain). **B**) Representative confocal images of tobacco pollen tubes expressing LAT52p-driven YFP fusions. Nuclear and cytoplasmic localization is visualized with a green-fire-blue LUT to highlight relative intensity (scale bar = 20 μm). **C)** Confocal images of stable Arabidopsis Col-0 lines expressing YFP-only and YFP-tagged variants of PLA_2_-like and JAZ1 proteins (scale bar = 50 μm). **D**) Quantification of nucleus-to-cytoplasm (N/C) fluorescence ratio in pollen tubes. Log_10_-transformed data from ∼30 cells across three independent experiments are shown. Distinct letters denote statistically significant differences (Kruskal-Wallis test with Holm multiple-comparison correction, p < 0.05). **E**) Quantification of N/C fluorescence ratio in stable Arabidopsis lines. Data from ∼30 cells across two independent lines for each construct except PLA_2_-like FL (one line) were log₁₀-transformed. Distinct letters denote statistically significant differences (Kruskal-Wallis test with Holm multiple-comparison correction, p < 0.05). **F)** Fluorescence recovery after photobleaching (FRAP) analysis of full-length PLA_2_-like and PLA_2_-like PY^AA^. Representative FRAP time-course images showing pre-bleach, post-bleach, and recovery stages for PLA_2_-like and PLA_2_-like PY^AA^. Time after bleaching is indicated in each panel. Scale bar = 10 μm. **G)** Nuclear fluorescence recovery was monitored following photobleaching, full-scale normalized and fitted to derive the mobile fraction (Mf) and half-time of recovery (T_0.5_). Boxplots summarize T_0.5_ values obtained from independent FRAP experiments. Statistical significance was determined using a two-tailed Mann–Whitney U test (P < 0.00005).

To test whether a bona-fide plant PY-NLS pathway operates *in vivo*, we compared full-length proteins, isolated motifs, and PY-to-AA (PY^AA^) point-mutated versions in *Nicotiana benthamiana* leaves and in transiently transformed *Nicotiana tabacum* pollen tubes. The pollen-tube system enables rapid screening of multiple constructs and, because of its large vegetative nucleus and vacuole-free apical cytoplasm, allows convenient quantification of nuclear/cytoplasmic partitioning of fluorescently tagged proteins. The predicted PY-NLS motifs from both PLA_2_-like and JAZ1 localized strongly to the nucleus (Fig. 1B, D). Crucially, nuclear localization of the full-length proteins and isolated PY-NLS motifs was significantly reduced for PLA_2_-like and completely lost for JAZ1 when the PY dipeptide was mutated to alanine-alanine (Fig. 1B, D). Free YFP, histone H2B, and the human chimeric karyopherin 2 inhibitor M9M motif were included as controls for quantification of the nuclear-to-cytoplasmic (N/C) fluorescence ratio (Fig. 1B, D).

To confirm that the PY-NLS system functions beyond transient expression systems, we generated stable Arabidopsis lines expressing N-terminal YFP fusions under the UBQ10 promoter and quantified their nucleocytoplasmic partitioning in roots. Full-length PLA_2_-like and JAZ1 accumulated in the nucleus, whereas substitution of the PY residues with alanines in PLA_2_-like (PLA_2_-like PY^AA^) reduced nuclear accumulation and deletion of the Jas domain from JAZ1 (ΔJas) eliminated nuclear enrichment (Fig. 1C, E). The loss of nuclear enrichment caused by the ΔJas deletion was also observed in tobacco pollen tubes (Fig. 1B, D), consistent with Withers et al. (2012). To determine whether the PY-NLS motifs of JAZ1 and PLA_2_-like are interchangeable, we generated a chimeric JAZ1 construct in which the deleted Jas domain was replaced with the PLA_2_-like PY-NLS motif. This substitution fully restored JAZ1 nuclear targeting in transient expression systems (Fig. 1B, D) and in stable Arabidopsis lines (Fig. 1C, E). Co-expression with the nuclear marker mCherry–H2B in *N. benthamiana* leaves followed by quantitative analysis independently corroborated the findings from tobacco pollen tubes and Arabidopsis roots (Supplementary Fig. 1).

To determine whether PLA_2_-like undergoes active and ongoing nucleocytoplasmic exchange rather than static nuclear retention, we performed fluorescence recovery after photobleaching (FRAP) on YFP-tagged PLA_2_-like wild-type (WT) and PY^AA^ mutant full-length proteins transiently expressed in *N. benthamiana* leaves (Fig. 1F). Following nuclear photobleaching, YFP:PLA_2_-like WT showed robust fluorescence recovery with a mobile fraction (Mf) of 0.56 ± 0.04 and a half-time of recovery (T_0.5_) of 35 ± 3 s, indicating rapid and efficient nuclear re-import of PLA_2_-like from the cytoplasmic pool (Fig. 1G), corresponding with the rate of nuclear import described for SIMKK or PHY1 proteins (Rausenberger et al., 2011; Ovečka et al., 2014). In contrast, the PY^AA^ mutant exhibited approximately two-fold slower recovery kinetics, with T_0.5_ prolonged to 66 ± 9 s (p < 0.00005), but eventually reached similar mobile fraction values (Mf = 0.63 ± 0.05; Fig. 1G).

Taken together, these results establish that plants possess a dynamic, functional, non-classical PY-NLS-dependent nuclear localization system, and that intact PY residues are essential for efficient nuclear import.

### Epitope composition and linker length shape plant PY-NLS activity

We noticed that both JAZ1 and PLA_2_-like differed from the sequence consensus established for animal and yeast PY-NLS motifs (Lee et al., 2006). Specifically, both proteins contained longer linkers between epitopes 1 and 2 and between epitopes 2 and 3, and they also differed in epitope 1 (Fig. 2A). To dissect the modular organization of plant PY-NLS motifs, we generated a series of engineered constructs varying in epitope composition and linker length (Fig. 2A) and tested their capacity for efficient nuclear import in the pollen-tube system (Fig. 2B, C). Extending epitope 1 in PLA_2_-like (PY-long) did not substantially improve nuclear import. Further, neither epitope 1 alone nor epitopes 2+3 alone localized to the nucleus, confirming the importance of all three epitopes for plant PY-NLS functionality (Fig. 2B, C). To test optimal linker length, we examined PLA_2_-like PY-NLS variants with linkers of 1–22 residues. The native 17-residue linker between epitopes 1 and 2 was optimal for nuclear localization, whereas the introduction of both shorter (1-13 residue) and longer (22 residue) variants resulted in significantly weaker nuclear import (Fig. 2C). Similar behavior was observed for the JAZ1 PY-NLS, which contains an additional epitope 1-like sequence in the linker (Fig. 1A), forming a motif variant conforming exactly to animal/yeast-like PY-NLS consensus. However, this variant (PY-short) performed poorly, again supporting a longer linker (∼16 aa) as optimal for plant PY-NLS activity (Fig. 2B, C). Finally, we tested the roles of epitopes 1 and 2 in JAZ1. While mutating epitope 1 reduced import to the level of the PY^AA^ (epitope 3 mutation), mutating the epitope 2 residue left import essentially intact (Fig. 2B, C)

**Figure 2.**
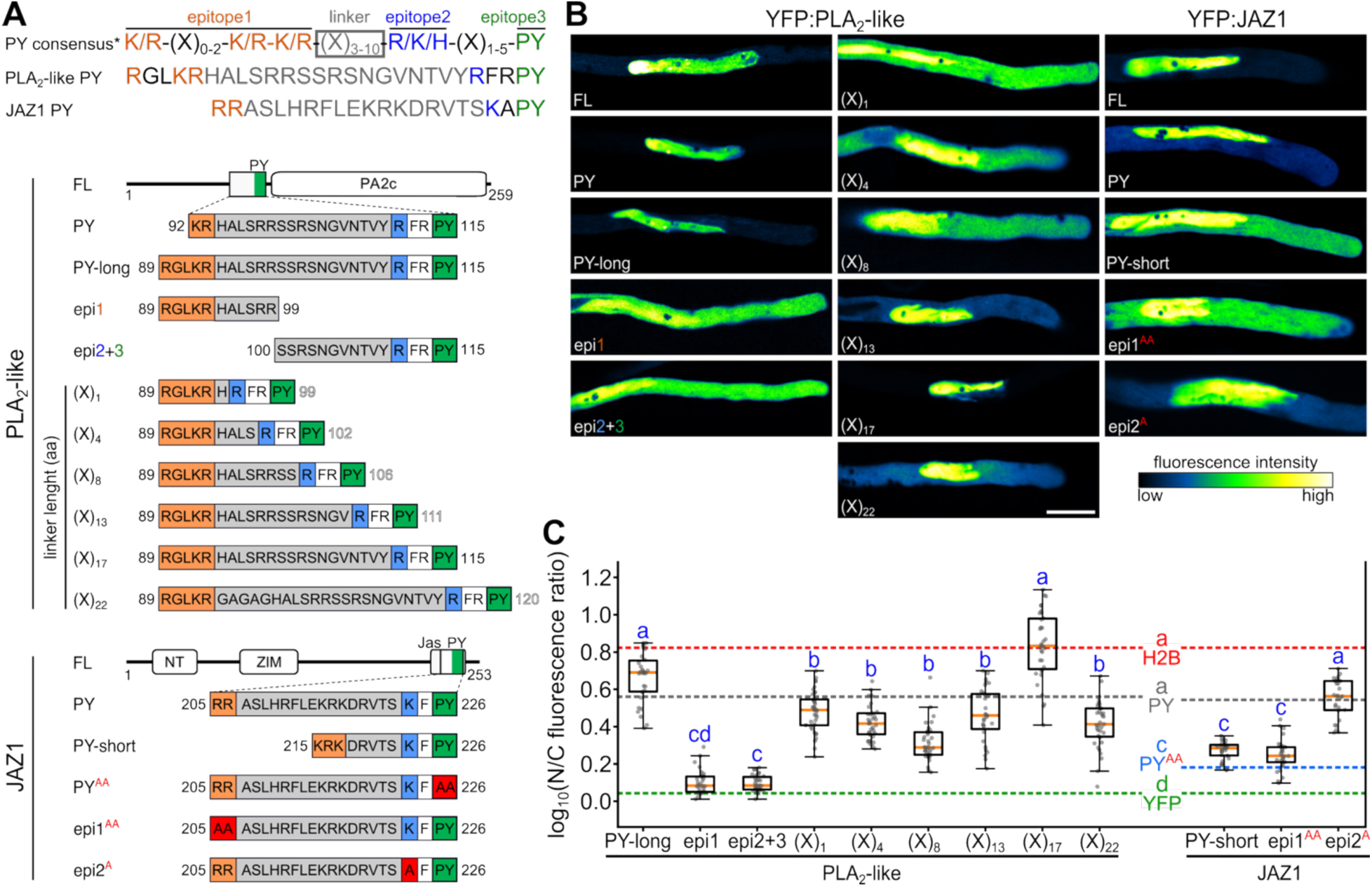
Linker length and epitope modularity define optimal plant PY-NLS activity. **A)** Top: Human consensus for PY-NLS motifs highlighting basic epitope 1 with variable-length linker (Xₙ), epitope 2 and conserved PY residues (epitope 3). Bottom: Schematic representation of PLA_2_-like and JAZ1 PY motif variants, including truncated and mutated constructs expressed in pollen tubes. **B)** Confocal images of tobacco pollen tubes expressing the indicated YFP fusion constructs of PLA_2_-like and JAZ1 variants (scale bar = 20 μm). **C)** Quantification of N/C fluorescence ratios in pollen tubes for the indicated constructs. Log_10_-transformed values from ∼30 cells are plotted. Horizontal red, green, grey and blue lines denote reference values for H2B, free YFP, PY motifs and JAZ1 PY-to-AA mutated motif, respectively (data from Fig. 1). Different letters indicate significant differences (Kruskal-Wallis test with Holm multiple-comparison correction, p < 0.05).

Collectively, these results establish optimal epitope spacing as an important determinant of plant PY-NLS functionality and confirm the modularity and functional portability of PY-NLS motifs in plants.

### PLA_2_-like undergoes a TRN1-dependent non-classical nuclear import pathway

To further investigate the mechanism underlying PLA_2_-like nuclear localization, we assessed whether PLA_2_-like is recognized by members of the Arabidopsis importin-β (IMB) family using AlphaFold multimer predictions. Among all tested IMB members, TRANSPORTIN 1 (TRN1; IMB2) yielded the highest confidence interaction scores with the PLA_2_-like, JAZ1 and M9M PY-NLS motifs (Fig. 3A). To gain insight into TRN1–PY-NLS interactions, we employed AlphaFold structural modeling, which have shown that PLA_2_-like and JAZ1 PY-NLS motifs form conserved TRN1-binding interfaces similar to the synthetic M9M motif and the experimentally solved structure of the human hnRNPM PY-NLS complex (Fig. 3B). Analysis of PAE-based contact confidences and plots supported high-confidence interactions for wild-type motifs but not PY^AA^ mutants (Fig. 3B and 3C, left panel). An AlphaFold multimer combined-scoring metric (0.8*ipTM + 0.2*pTM, Omidi et al., 2024) further indicated a significant propensity for interaction in wild-type motifs, which was reduced in PY^AA^ mutants (Fig. 3C, right panel).

**Figure 3.**
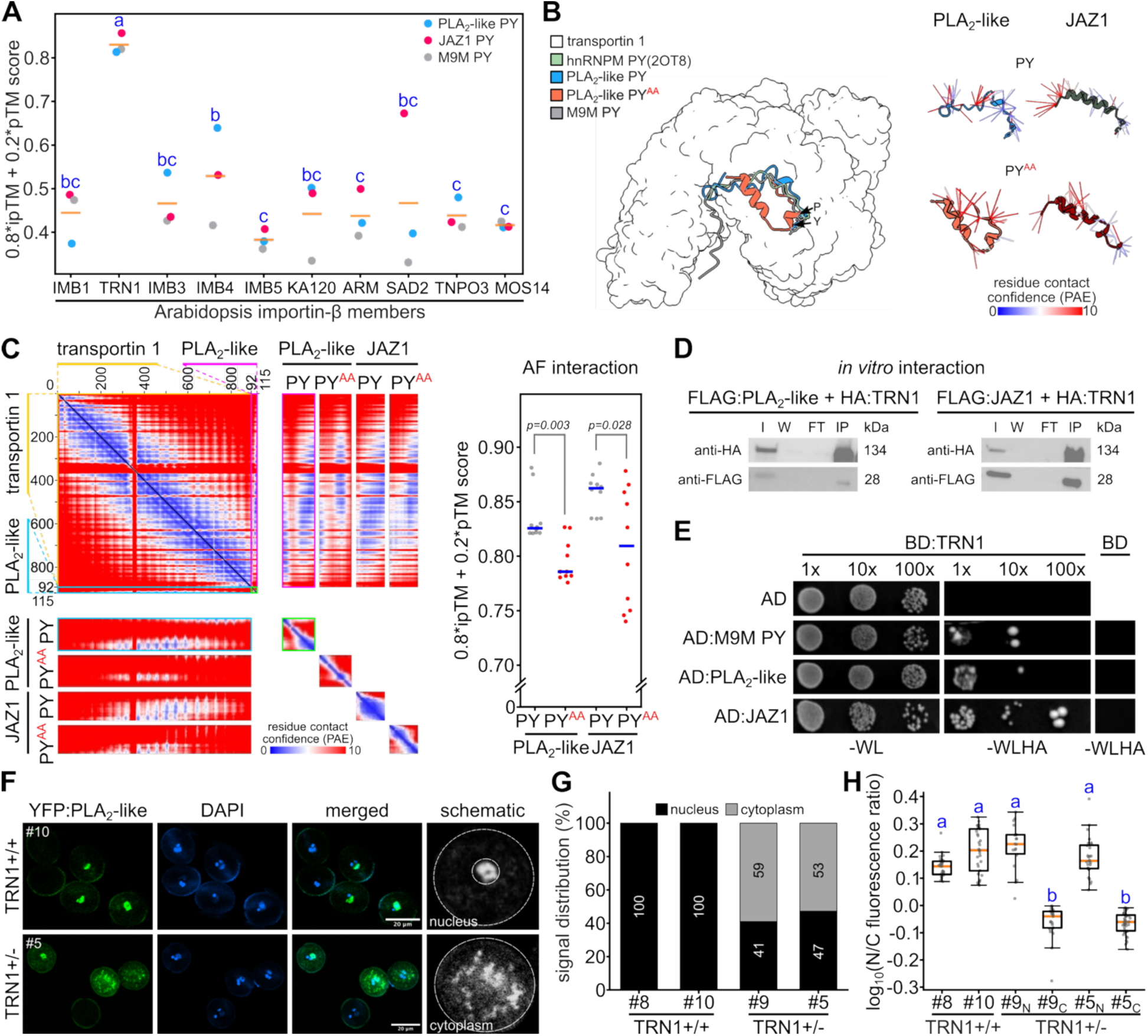
Arabidopsis TRANSPORTIN 1 (TRN1) recognizes plant PY-NLS cargoes and mediates nuclear trafficking. **A)** *In silico* prediction of interactions between Arabidopsis importin-β family members and PY-NLS cargoes. AlphaFold-multimer models were generated using the PY-NLS motifs of PLA_2_-like, JAZ1, and the high-affinity TRN1 inhibitor M9M. Interaction confidence was assessed using the composite ranking score (0.8*ipTM + 0.2*pTM). Among Arabidopsis importin-β family members, TRN1 displayed the highest predicted interaction scores with all three PY-NLS-containing ligands. **B)** Left: Structural prediction of TRN1 complexes with human M9M and plant PLA_2_-like and JAZ1 PY-NLS motifs using AlphaFold. The human hnRNPM PY-NLS complex (PDB: 2OT8) is shown as a reference. Right: PAE-colored contact maps highlight high-confidence residue interactions for WT (PY) and mutant motifs (PY^AA^). **C)** Full PAE plots from AlphaFold predictions of TRN1 complexes with WT and mutant PY-NLS motifs (PY^AA^). Insets: high-resolution views of TRN1-PY motif interfaces. Right: Quantitative scoring of the top 10 AlphaFold models (based on combined pTM and ipTM scores). Significant differences in predicted interaction strength are indicated (Kruskal-Wallis test). **D)** Pull-down assay performed with *in vitro* translated HA:TRN1 as a bait and FLAG:PLA_2_-like and FLAG:JAZ1 as preys. I, input; W, wash; FT, flow-through; IP, immunoprecipitated fraction. **E)** Yeast two-hybrid assay showing interactions between Arabidopsis TRN1 and PLA_2_-like and JAZ1 full-lengths and synthetic M9M PY motif. Growth on selective medium (dropout tryptophan, leucine, histidine and adenine, -WLHA) indicates interaction. **F)** Subcellular localization of YFP-tagged PLA_2_-like in mature Arabidopsis pollen grains derived from wild-type (TRN1+/+) and heterozygous (TRN1+/−) plants. Representative confocal images show reduced nuclear accumulation and increased cytoplasmic distribution of PLA_2_-like in pollen presumably carrying the mutant *trn1* allele. **G)** Percentages of pollen grains assigned to nuclear-enriched or cytoplasmic localization classes in two independent lines of each parental genotype. **H)** Log_10_-transformed nuclear-to-cytoplasmic fluorescence ratios in TRN1+/+ pollen and in the nuclear (N) and cytoplasmic (C) classes from TRN1+/− plants. Different letters indicate significant differences (Kruskal-Wallis test with Holm multiple-comparison correction, p < 0.05).

To experimentally validate the predicted interaction between TRN1 and JAZ1 and PLA_2_-like cargos, we employed a pull-down assay with *in vitro*-translated proteins. Both FLAG:PLA_2_-like and FLAG:JAZ1 bound HA:TRN1 and were recovered in the eluted fractions (Fig. 3D). confirming a direct physical association between TRN1 and the PLA_2_-like and JAZ1 cargo proteins. The interaction was further confirmed by a yeast two-hybrid assay (Fig. 3E).

Finally, to test whether TRN1 is required for cargo nuclear localization *in vivo*, we examined YFP:PLA_2_-like distribution in Arabidopsis lines carrying different TRN1 genotypes. Homozygous *trn1* mutants exhibited severe developmental defects and failed to produce viable progeny (Cui et al., 2016, Supplementary Fig. 2A-C). Qualitative imaging of mutant roots revealed reduced nuclear accumulation of YFP:PLA_2_-like together with prominent cytoplasmic and punctate signals (Supplementary Fig. 2D). Because the abnormal root morphology and heterogeneous extranuclear signal precluded reliable nucleocytoplasmic quantification, we exploited segregation of the TRN1 locus in haploid pollen from heterozygous plants.

In pollen from two TRN1+/+ lines (#8 and #10), YFP: PLA_2_-like localized exclusively to DAPI-positive vegetative nuclei (Fig. 3F, G). By contrast, pollen from two TRN1+/− lines (#9 and #5) segregated into two localization classes: 41% and 47% of grains, respectively, retained nuclear localization, whereas 59% and 53%, respectively, displayed a granular cytoplasmic signal (Fig. 3F, G). This near-1:1 distribution is consistent with segregation of the TRN1 and *trn1* alleles and supports impaired PLA_2_-like nuclear localization in *trn1*-bearing pollen. Accordingly, cytoplasmic-class grains had significantly lower nuclear-to-cytoplasmic fluorescence ratios than TRN1+/+ pollen and nuclear-class grains from the same heterozygous lines (Fig. 3H). Together, the root and pollen data support a requirement for TRN1 in efficient nuclear localization of PLA_2_-like *in vivo*.

Taken together, structural modeling, biochemical interaction assays, and *in vivo* localization analyses identify TRN1 as a receptor for the PY-NLS cargoes PLA_2_-like and JAZ1 and define a TRN1-dependent non-classical nuclear import pathway in plants.

### Proteome-wide identification and functional validation of PY-NLS-containing cargoes in Arabidopsis

Building on our experimental and computational analyses of the PLA_2_-like and JAZ1 PY-NLS motifs, we developed a pipeline to predict functional PY-NLSs across the Arabidopsis proteome (Supplementary Data S1). The initial pattern scan used an updated consensus comprising three epitopes: epitope 1, containing two basic residues; a 3–20-aa linker; epitope 2, comprising a K, R, or H residue; a 1–5-aa linker; and epitope 3, comprising the PY dipeptide. Candidate motifs were then evaluated and filtered using sequence- and structure-based features, including secondary structure, linker geometry, intrinsic disorder, AlphaFold confidence, relative solvent accessibility, and motif surface exposure. Protein-level filters further excluded candidates containing predicted classical NLSs, transmembrane domains, or signal or transit peptides, while optional TRN1 co-folding and interface analyses provided additional interaction-based evidence (Fig. 4A).

**Figure 4.**
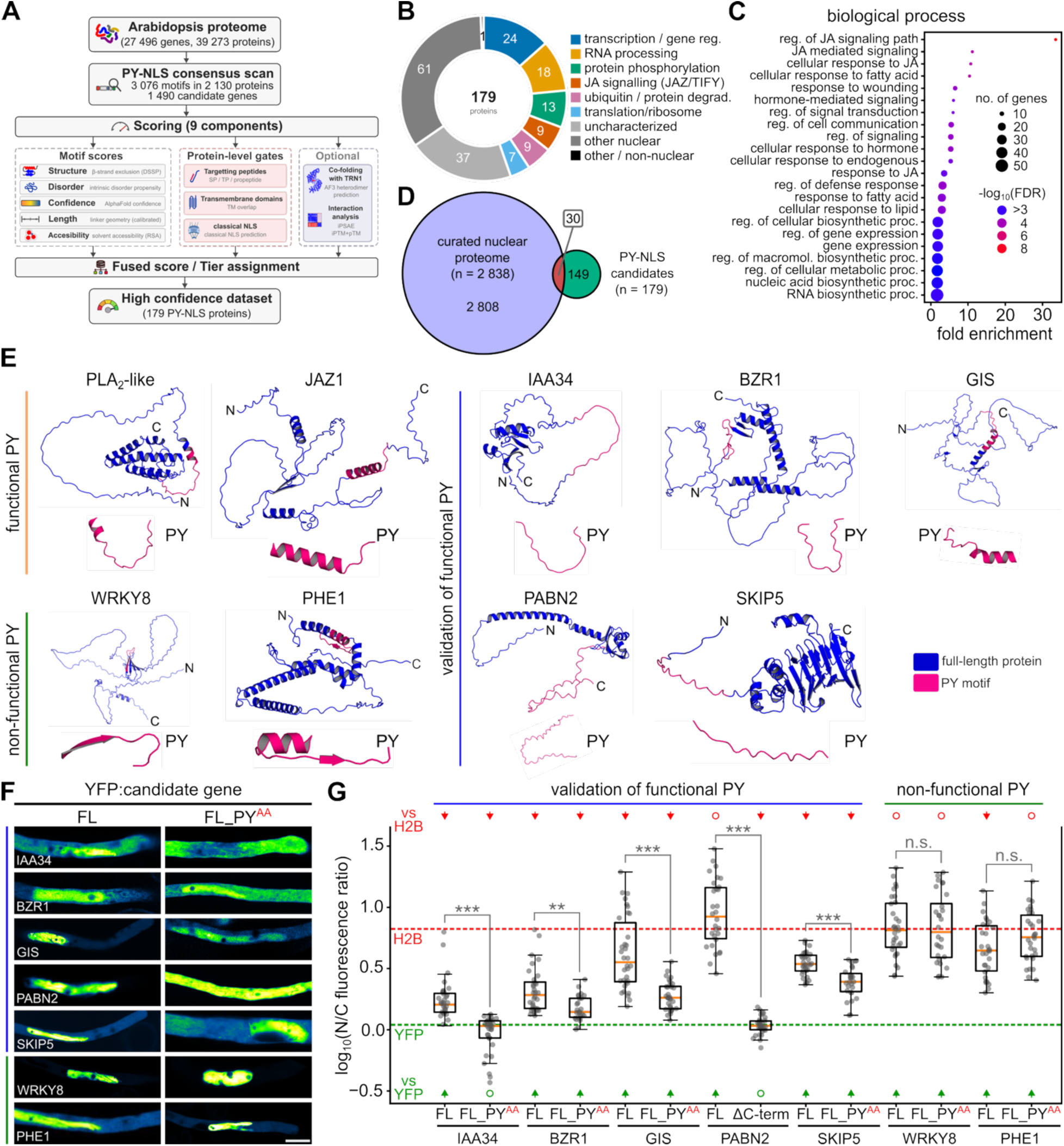
Proteome-wide identification and experimental validation of functional PY-NLS cargo proteins in Arabidopsis. **A)** Schematic pipeline for identification of high-confidence PY-NLS-containing cargoes in the Arabidopsis proteome using structural filters (secondary structure, linker-length constraints, and exclusion of β-sheet-associated motifs), confidence filters (AlphaFold pLDDT scores, relative solvent accessibility (RSA), and motif exposure), and localization filters that removed proteins with classical nuclear localization signals (cNLSs), transmembrane domains, signal peptides, or organellar transit peptides. **B)** Functional classification of the 179 predicted PY-NLS-containing proteins. Each protein was assigned its primary molecular function from UniProt keywords, and groups with >= 5 members were kept; the remainder are shown as uncharacterized or as a nuclear / non-nuclear localization residual. **C)** Gene Ontology (GO) enrichment analysis of identified PY-NLS cargoes. Dot size reflects gene counts per term; color indicates statistical significance (FDR-corrected p-value). **D)** Venn diagram comparing PY-NLS cargoes with nuclear proteome datasets (Ayash et al., 2021). **E)** AlphaFold structural models of selected Arabidopsis proteins containing predicted PY-NLS motifs. Full-length proteins are shown in blue; PY motifs in magenta. N- and C-termini and PY positions are indicated. **F)** Representative confocal images of YFP-tagged candidate proteins expressed in tobacco pollen tubes under the LAT52 promoter. Full-length proteins and corresponding PY-NLS mutant variants (PY→AA or C-terminal deletion, where indicated) were examined for nuclear accumulation. **G)** Quantification of nuclear localization expressed as the log_10_ nuclear-to-cytoplasmic (N/C) fluorescence ratio. The green and red reference lines represent the average N/C ratios of YFP:Histone H2B and free YFP controls, respectively. Brackets indicate the comparisons between full-length proteins and their corresponding PY-NLS mutants or deletion variants; *P* values were adjusted by the Holm method across these seven tests (*** *P*<0.001, ** *P*<0.01, n.s. not significant). Arrows indicate significantly higher or lower N/C ratios relative to YFP or H2B control; open circles indicate that no significant difference was detected.

Application of the pipeline to the complete Arabidopsis proteome identified 3,076 consensus matches in 2,130 protein sequences encoded by 1,490 genes. Of these, 179 proteins passed the filtering criteria and formed the high-confidence PY-NLS dataset. Functional classification based on UniProt annotations revealed diverse protein categories, including transcription and gene regulation (24 proteins), RNA processing (18), protein phosphorylation (13), jasmonate signaling (9), protein degradation (9) and translation (7) (Fig. 4B). Gene ontology enrichment analysis identified overrepresented biological processes related to jasmonic acid signaling, responses to fatty acids and wounding, defense regulation, RNA biosynthesis, and transcriptional regulation (Fig. 4C). The enrichment of RNA-related and gene-regulatory functions is consistent with the known activities of animal TRN1 cargoes (Lee et al., 2006). Comparison with a curated Arabidopsis nuclear proteome (Ayash et al., 2021) showed that only 30 of the 179 candidates (17%) were represented in this reference dataset (Fig. 4D). Thus, most predicted cargoes are absent from existing nuclear proteome maps and may include condition- or cell-type-specific nuclear proteins as well as previously unrecognized import substrates.

To assess the predictive value of the PY-NLS identification pipeline, we first used AlphaFold 3 to model complexes of TRN1 with the PY-NLS regions of 20 proteins bearing predicted functional motifs and 20 proteins bearing predicted non-functional motifs, including the characterized JAZ1 and PLA_2_-like PY-NLSs, and the corresponding PY^AA^ mutant of each construct. Predicted functional motifs formed higher-confidence interfaces with TRN1 than predicted non-functional motifs, and PY^AA^ substitution significantly reduced interface confidence for the functional motifs but not for the non-functional ones (Supplementary Fig. 3A). The analysis of detailed per-residue interface scoring (ipSAE) across the motif region showed that both the difference between functional and non-functional motifs and the effect of PY^AA^ substitution were indeed concentrated around the PY dipeptide (Supplementary Figure 3B). This corroborates the pipeline’s ability to predict motifs capable of engaging the TRN1 cargo-binding site in a PY-dependent manner.

To further test the PY-NLS predictions experimentally, we selected five candidates with diverse sequences and predicted structures: INDOLEACETIC ACID-INDUCED PROTEIN 34 (IAA34), BRASSINAZOLE-RESISTANT 1 (BZR1), GLABROUS INFLORESCENCE STEMS (GIS), POLYADENYLATE-BINDING PROTEIN 2 (PABN2), and SKP1-INTERACTING PARTNER 5 (SKIP5). All five proteins contain predicted PY-NLSs but lack a predicted classical NLS (Supplementary Fig. 3, Supplementary Data 2). The PY-NLS regions of IAA34, BZR1, PABN2, and SKIP5 were predominantly disordered, whereas the GIS motif contained an α-helical segment upstream of the PY dipeptide. PABN2 contained several clustered PY-NLS motifs near its C-terminus (Fig. 4E). We also selected the transcription factors WRKY8 and PHE1 as predicted negative controls. Both contain PY-NLS consensus matches, but these occur within structured regions containing β-strands and were therefore predicted to be non-functional (Fig. 4E, Supplementary Fig. 3).

We expressed each full-length candidate as a YFP fusion together with its corresponding PY^AA^ mutant and quantified nuclear-to-cytoplasmic partitioning. All five predicted functional candidates showed nuclear enrichment in their full-length forms, and the introduction of PY^AA^ substitutions reduced their nuclear accumulation (Fig. 4F, G). Because PABN2 contains multiple clustered motifs, we instead deleted its C-terminal region; this abolished nuclear enrichment, confirming that the functional targeting determinant resides within this region. By contrast, PY^AA^ substitutions did not significantly alter the localization or nuclear-to-cytoplasmic partitioning of WRKY8 or PHE1, consistent with the predicted non-functional configuration of their consensus motifs.

Finally, to examine whether PY-NLS-mediated import targets similar cargo functions across plants, we predicted PY-NLS candidate proteins from reference proteomes of five additional angiosperms spanning the major lineages: rosid *Vitis vinifer*a, asterid *Solanum lycopersicum*, monocots *Hordeum vulgare* and *Brachypodium distachyon*, and Amborella trichopoda, the basal angiosperm from the ANA-grade (Supplementary Data S2), and compared them with Arabidopsis. Gene ontology enrichment analysis of the resulting high-confidence candidate sets revealed a conserved functional core, with 15 biological process terms shared by at least four species and enriched for jasmonate signaling and related wound, fatty acid, and defense responses (Supplementary Figure S4A, B). Grouping the enriched terms into functional themes further confirmed jasmonate/oxylipin signaling and defense, and RNA metabolism, processing, and transport as the categories most consistently enriched across all six species, along with general stress responses and gene expression regulation (Supplementary Figure S4C). In addition, each species exhibited independently enriched terms, indicating lineage-specific recruitment of PY-NLS cargoes.

Collectively, our results demonstrate that sequence context and structural accessibility are key predictors of functional PY-NLS for TRN1-mediated nuclear import in plants and can be leveraged to predict functional import substrates proteome-wide.

## DISCUSSION

The present study provides the comprehensive functional characterization of proline-tyrosine nuclear localization signals (PY-NLSs) and their recognition by transportin 1 (TRN1) in plants, simultaneously establishing a plant-specific paradigm for this type of non-classical nuclear import. Although animal and yeast PY-NLSs are well described and mediate nuclear import of numerous cargoes via ortholog of Arabidopsis TRN1, analogous pathways in plants have remained poorly defined, with most plant NLS studies focused on classical monopartite and bipartite signals recognized by canonical importin-α and importin-β proteins (Lange et al., 2007; Chen et al., 2020; Mori et al., 2025). Our findings extend this framework by demonstrating that plants possess bona-fide functional PY-NLS motifs and a TRN1-dependent import pathway that follows the same general logic as other eukaryotic systems but has distinct plant-specific features.

Plant PY-NLS motifs represented by proteins tested in this study differ from the canonical animal and yeast consensus in several key aspects. As in animal and yeast systems, plant PY-NLS motifs are modular: efficient nuclear import requires a positively charged N-terminal region (epitope 1), a central linker, a conserved K/R/H residue (epitope 2), and a C-terminal PY dipeptide (epitope 3). However, the optimal linker between epitopes 1 and 2 is longer in plants (16–18 amino acids), and the minimum linker between epitopes 2 and 3 is one residue, whereas a short animal-like form performs poorly in nuclear import assays. These differences suggest lineage-specific consensus requirements for TRN1 recognition in the plant kingdom. Because the interaction of epitope 1 with TRN1 remains incompletely defined (Fig. 3B; Twyffels et al., 2014), the structural basis of this evolutionary difference is still unclear. Lee et al. (2006) used isothermal calorimetry to analyze M9NLS mutant interactions with Kapβ2 (the animal ortholog of TRN1) and demonstrated that glycine in epitope 1 and the R and PY residues in epitopes 2 and 3 were crucial for Kapβ2 interaction. Structural modeling in the present study further indicates that functional plant PY-NLS motifs can also contain a short α-helix upstream of the disordered PY residues. Conversely, motifs without disordered features, containing residues in β-sheet contexts or in buried conformations, fail to efficiently direct nuclear localization.

The functional confirmation and sequence/structure analysis of PY-NLS motifs in PLA_2_-like and JAZ1 enabled the identification of a broader catalog of candidate nuclear import cargoes in Arabidopsis. Many predicted PY-NLS proteins identified here, as in the pioneering work of Lee et al. (2006), are implicated in RNA biology. At the same time, the plant-specific determinants of a functional PY-NLS underscore the evolutionary flexibility of this signal type in plants. We hypothesize that the plant-specific PY-NLS architecture emerged after major plant-lineage divergence through adaptation of pre-existing karyopherin recognition surfaces to new cargo classes. A clear example of this is the presence of PY-NLS in the recently identified PLA_2_-like family, which appears to be plant-specific (Saddhe and Potocký, 2023). Comparative testing across algae, bryophytes, and vascular plants will be needed to determine when the longer linker and upstream α-helical features became fixed features of plant PY-NLS motifs.

Notably, the presence of PY-NLS motifs in all JAZ family proteins (Supplementary Data S2) suggests a biological rationale for a dedicated TRN1 pathway in hormone signaling. JAZ repressors are rapidly deployed nuclear regulators whose abundance and interactions change during jasmonate responses, and the presence of PY-NLS/TRN1-pathway could allow their nuclear entry to be regulated independently of the classical importin-α pathway and its many competing cargoes. Such separation may be especially useful when stress, development, or pathogen responses alter the demand for classical NLS import. Interestingly, the physical interaction between MYC2 and the Jas domain of both JAZ1 and JAZ9 was proposed to mediate their nuclear import (Withers et al., 2012). However, the ability of the PLA_2_-like PY-NLS to restore nuclear localization of JAZ1ΔJas expands this hypothesis and indicates that JAZ nuclear targeting can be mediated by multiple, partly independent import signals.

These findings also point to potential crosstalk between PY-NLS and cNLS pathways. In Arabidopsis, 163 proteins were predicted to carry both signal types. Competition between import pathways, changes in importin abundance, or stress-induced modification of cargoes could therefore shift the balance between TRN1-dependent and importin-α-dependent nuclear entry.

Notably, our TRN1-related data corroborate those of Cui et al. (2016), who reported that a *trn1* loss-of-function line exhibits severe developmental defects, further underscoring the importance of *trn1* for plant growth and development. Since this work demonstrated that TRN1 in plants can function in nuclear import via PY-NLS recognition, it is tempting to speculate that some of the pleiotropic *trn1* phenotypes are caused by perturbed nuclear localization of plant PY-NLS cargoes. It is important to emphasize that PY-NLS motifs show modular behavior, where individual epitopes can behave as quasi-independent units contributing variably to overall binding affinity towards the TRN1 and that individual motifs display a range of TRN1 affinities (Süel et al., 2008). This framework is also consistent with our observation that plant motifs differ significantly in import efficiency even among validated cargoes.

In summary, this work lays the groundwork for a more comprehensive examination of nuclear import diversity in plants. The predictive rules established here can now be deployed for genome-wide annotation of PY-NLS cargoes, aiding the discovery of previously unrecognized nuclear regulatory pathways. Future investigations will be needed to dissect the full cargo spectrum of TRN1 *in planta*, clarify the mechanisms underlying PY-NLS structural requirements, and explore the prevalence and function of similar non-classical import pathways in other plant lineages. By extending the knowledge of PY_NLS_ motif architecture, consensus, and function to the plant kingdom, our study bridges a major evolutionary gap in nuclear transport research and opens new vistas for mechanistic and applied work in plant nuclear biology.

## MATERIALS AND METHODS

### Plant material and growth conditions

*Arabidopsis thaliana* seeds were surface sterilized, stratified for 2 days at 4 °C, and germinated on vertical ½ Murashige-Skoog medium supplemented with 1% sucrose, vitamin mixture, and 1.6% plant agar (Duchefa Biochemie). Young seedlings were transferred to pellets (Jiffy Products International) and moved to *ex vitro* conditions (22 °C, 16/8 h light/dark regime). The T-DNA insertion line SALK_003127 of TRN1 (designated *trn1*) was obtained from the European Arabidopsis Stock Centre (NASC) and genotyped using the listed primer combinations (Supplementary Table 1). After genotyping, heterozygous *trn1* plants were transformed with pUBQ10:YFP-PLA_2_-like. Transformants were selected with hygromycin and genotyped. Flowers from heterozygous *trn1* lines were used for pollen imaging, and pollen nuclei were visualized by DAPI staining.

To prepare stable transgenic lines of *A. thaliana* Col-0, plants were transformed with pUBQ10:YFP constructs using *Agrobacterium tumefaciens* strain GV3101 by the floral dip method. YFP-positive seedlings were visually selected and transferred to soil. The phenotype was documented 8 weeks after germination.

### Molecular cloning

Total RNA was extracted from vegetative and reproductive tissues of *Arabidopsis thaliana* (Col-0) using the RNeasy Kit (Qiagen). Genomic DNA was removed using the TURBO DNA-free Kit (Applied Biosystems). First-strand cDNA synthesis was performed using the Transcriptor High Fidelity cDNA Synthesis Kit (Roche) with anchored oligo(dT)₁₈ primers; the quality was validated via amplification of the ACTIN7 gene.

Full-length coding sequences (CDS) for PLA_2_-like (AT4G29070), JAZ1 (AT1G19180), WRKY8 (AT5G46350), PHE1 (AT1G65330), and TRN1 (AT2G16950) were amplified using Phusion DNA polymerase (NEB) with standard PCR parameters. PCR products were cloned into pWEN240 (pLAT52::YFP:CDS), pHD32 (pLAT52::CDS:YFP), or pHD71 (together with pUBQ10 promoter, pUBQ10::YFP:CDS) vectors (Klahre et al., 2006) using NgoMIV and ApaI restriction sites unless otherwise noted. Wild-type and mutant PY-NLS motifs were cloned by primer annealing into the corresponding vector as full-lenght proteins.

Site-directed mutagenesis was used to generate mutants AtPLA_2_-like PY114/115AA and AtJAZ1 PY225/226AA. Each mutant was produced by first generating a megaprimer (e.g., using primers AS193 and AS194 for AtPLA_2_-like PY114/115AA with the corresponding template), followed by amplification of the full-length mutant and cloning into the corresponding vector. N-terminal truncations were engineered using the same approach.

Histone 2B (AT3G45980) was cloned into pWEN240 and to pHD71 (together with pUBQ10 promoter and mCherry) vectors using XmaI and ApaI. The synthetic human M9M PY-NLS motif was cloned by primer annealing into pWEN240 via NgoMIV and ApaI sites.

### Transient expression in tobacco pollen tubes and *Nicotiana benthamiana* leaves

Tobacco pollen transformation was performed via particle bombardment using a helium-driven gene gun (PDS-1000/He; Bio-Rad) as previously described (Pejchar et al., 2020), with 1 µg DNA-coated gold particles per shot.

For leaf expression, pUBQ10-driven constructs were transformed into *A. tumefaciens* strain GV3101 and infiltrated into *N. benthamiana* leaves, which were incubated for 36–72 h at room temperature before imaging.

### Confocal microscopy and image analysis

Live-cell imaging was conducted with a Yokogawa CSU-X1 spinning disk confocal unit mounted on a Nikon Ti-E microscope using a 60× water immersion objective (NA 1.2) and an Andor Zyla sCMOS camera. A 514 nm laser was used for YFP excitation, and emission was collected through a Semrock brightline 542/27 nm filter. For mCherry imaging, 561 nm laser was used for excitation, and emission was collected through a Semrock brightline 641/75 nm filter. All imaging parameters were held constant across experimental conditions to ensure comparability.

ImageJ/Fiji (Schindelin et al., 2012) was used for quantitative image processing; nuclear-to-cytoplasmic (N/C) ratios of fluorescence intensity were calculated for ∼30 cells per construct. Statistical analyses were conducted in R using the Kruskal–Wallis test, with Holm correction for multiple comparisons. Unless otherwise stated, *P*-values < 0.05 were considered significant.

### Yeast two-hybrid (Y2H) assays

The full-length CDS of Arabidopsis TRN1 (AT2G16950) was cloned into the pGBKT7 bait vector using EcoRI and SalI. Corresponding prey constructs (full-length PLA_2_-like, full-length JAZ1, and human M9M PY-NLS motif) were cloned into pGADT7 via EcoRI and BamHI. Final constructs were co-transformed into *Saccharomyces cerevisiae* AH109 using the lithium acetate method. Transformants were selected on SD/-Trp/-Leu/ (-WL) medium, and protein–protein interactions were assessed by spotting serial dilutions (OD_600_ = 1 (1x), 1:10 and 1:100 dilutions) onto dropout media SD/-Trp/-Leu/-His/-Ade (-WLHA) and incubating at 30 °C for 2–5 days.

### *In vitro* translation and pull-down assay

Full-length HA:TRN1, FLAG:PLA_2_-like, and FLAG:JAZ1 constructs were cloned with N-terminal tags into the modified pTNT vector (Promega) using NgoMIV and ApaI restriction sites. For *in vitro* translation, TNT SP6 High-Yield Wheat Germ Protein Expression System (Promega) was used according to the manufacturer’s instructions, and translated proteins were confirmed by western blot. HA-tagged pull-down assays were then performed using Pierce HA-Tag Magnetic IP/Co-IP Kit (Thermo Scientific) to isolate the HA:TRN1 bait, followed by detection of the FLAG-tagged prey PLA_2_-like and JAZ1 proteins.

### *In silico* analysis of PY-NLS motifs in plant proteomes

Candidate PY-NLS motifs were identified using pynls_v1.py (version 1.0; Supplementary Data S1) in the *Arabidopsis thaliana* UniProt reference proteome. The pipeline searches for overlapping matches to the consensus [KR][KR].{3,20}[KRH].{1,5}PY, derived from established PY-NLS architecture (Lee et al., 2006; Yan et al., 2016) and our experimental analysis of PLA_2_-like and JAZ1.

Each motif was scored for linker geometry, intrinsic disorder, secondary structure, solvent accessibility, and AlphaFold pLDDT using the calibrated weights (see Supplementary Data S1 for more detailed description). Disorder was evaluated using AIUPred, metapredict, or IUPred3; secondary structure and solvent accessibility were calculated from AlphaFold models using DSSP and FreeSASA, respectively. Extended β-strand structure was considered incompatible with PY-NLS activity, whereas disordered and α-helical configurations were permitted. Proteins containing signal peptides, transit peptides, or propeptides were excluded. Motifs overlapping a transmembrane helix, located within 50 residues of one, or present in a polytopic loop shorter than 50 residues were also excluded; other transmembrane-associated candidates were downweighted by 50%. Detection of a strong classical NLS by cNLS Mapper resulted in a 15% score reduction. AlphaFold co-folding evidence for interaction with TRN1 was incorporated as an optional component.

Candidates were classified as high-confidence T1 (score ≥0.65), moderate-confidence T2 (≥0.45), weak or borderline T3 (≥0.25), or rejected T4 (<0.25 or subject to a structural, annotation, or topology veto). The highest-scoring motif was selected for each protein, and protein entries were collapsed by gene, yielding 179 high-confidence Arabidopsis candidates.

Interactions of Arabidopsis importin-β family members with the M9M, PLA_2_-like, and JAZ1 PY-NLS motifs were modeled separately using AlphaFold. Candidate proteins or their isolated motifs were similarly modeled with TRN1 and ranked using the combined metric 0.8*ipTM + 0.2*pTM (Omidi et al., 2024). Structures were visualized using ChimeraX (Meng et al., 2023). Venn diagrams were generated using the PSB Venn webtool, and Gene Ontology enrichment was analysed using ShinyGO v0.85.

### Accession numbers

The accession numbers of *A. thaliana* genes described in this manuscript are: PLA_2_-like (AT4G29070), JAZ1 (AT1G19180), WRKY8 (AT5G46350), PHE1 (AT1G65330), H2B (AT3G45980), TRN1/IMB2 (AT2G16950), IAA34 (AT1G15050), BZR1 (AT1G75080), SKIP5 (AT3G54480), GIS (AT3G58070), and PABN2 (AT5G65260).

## ACKNOWLEDGEMENTS

This work was supported by the Czech Science Foundation project 26-23500S to P.P. and by the Ministry of Education, Youth and Sports (MEYS) project TowArds Next GENeration Crops, reg. no. CZ.02.01.01/00/22_008/0004581 of the ERDF Programme Johannes Amos Comenius. The Imaging Facility of the Institute of Experimental Botany CAS is supported by the MEYS CR (LM2023050 Czech-BioImaging), the Czech Academy of Sciences, and Institute of Experimental Botany CAS. Computational resources were provided by the e-INFRA CZ project (ID:90254), supported by the Ministry of Education, Youth and Sports of the Czech Republic. During the preparation of this manuscript, the authors used ChatGPT to assist in developing the PY-NLS prediction pipeline and to refine the English language. The authors reviewed and edited all generated content.

