## Supplementary Information for "Sequence and structural rules define PY-NLS–dependent nuclear import by TRANSPORTIN 1 in plants"

### Contents

Supplementary Figures 1-4

Supplementary Tables 1-2

Supplementary Methods

Supplementary Data 1

Supplementary Data 2

Supplementary References

*Supplementary Data 1 and 2 are supplied as separate files.*

### SUPPLEMENTARY FIGURES

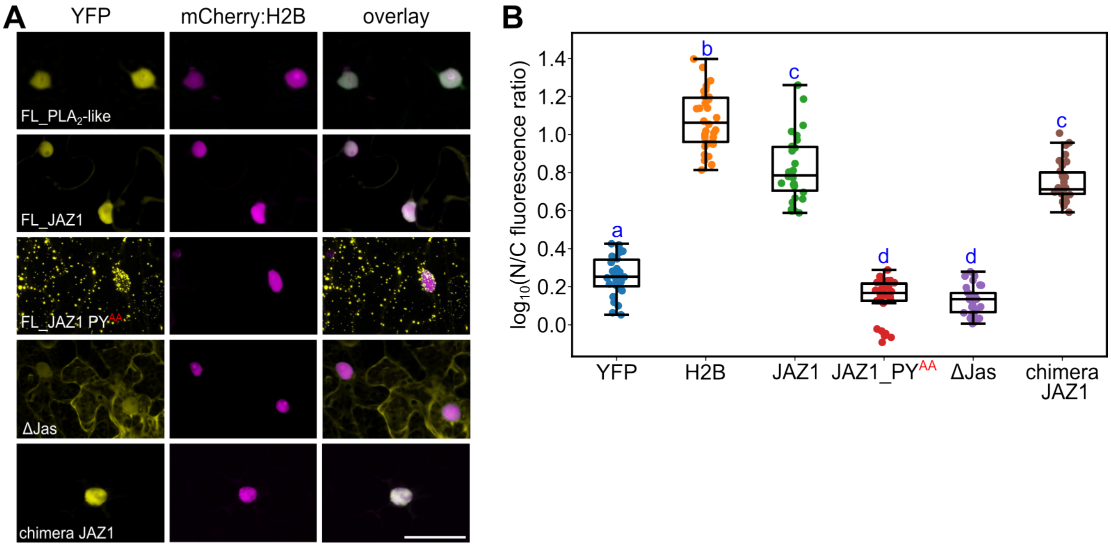


**Supplementary Figure 1**. **A)** A representative image of transient agro-infiltration of *N. benthamiana* leaves showing co-localization of YFP-PLA$_2$-l and JAZ1 variants with nuclear marker mCherry-Histone 2B. **B)** Quantification of nuclear localization expressed as the log10 nuclear-to-cytoplasmic (N/C) fluorescence ratio.

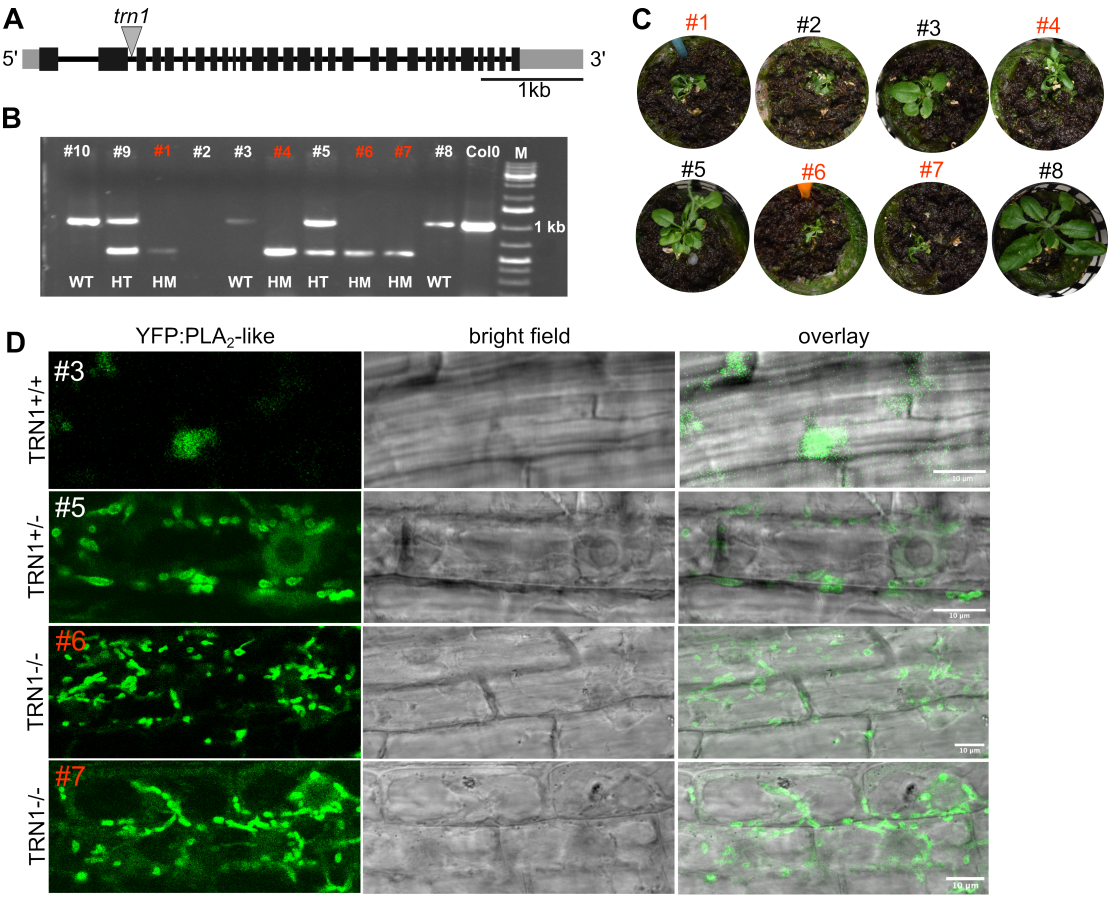


**Supplementary Figure 2. Genotyping and selection of TRN1 mutant lines used for in vivo localization analyses. A)** Schematic representation of the TRN1 locus showing the position of the T-DNA insertion. **B)** Representative PCR genotyping results for independent transgenic lines expressing YFP-$PLA_2$-like in the heterozygous (TRN1+/−) mutant background. Genotypes were assigned based on the presence of wild-type and T-DNA-specific amplification products. Lines carrying wild-type (TRN1+/+), heterozygous (TRN1+/−), and homozygous (TRN1−/−) alleles are indicated. HM, homozygous; HT, heterozygous; M, DNA ladder. **C)** Representative images of independent transgenic Arabidopsis lines expressing YFP-$PLA_2$-like in the TRN1(+/−) mutant background. Multiple independent transformants were recovered and screened for fluorescence expression and genotype. **D)** Qualitative confocal analysis of YFP–$PLA_2$-like localization in roots from representative independent lines carrying TRN1+/+, TRN1+/−, and TRN1−/− genotypes. YFP fluorescence, bright-field images, and merged channels are shown. Whereas YFP–$PLA_2$-like was enriched in nuclei of TRN1+/+ roots, *trn1* backgrounds showed reduced nuclear accumulation and increased cytoplasmic and punctate signals. Severe root abnormalities and heterogeneous extranuclear fluorescence precluded reliable quantitative segmentation. Scale bars are as indicated.

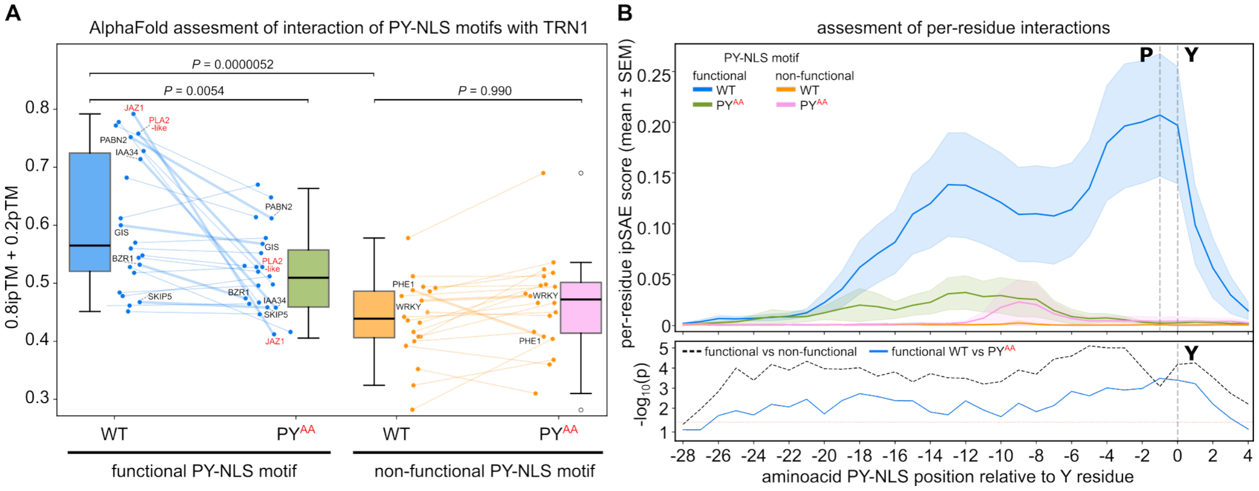


**Supplementary Figure 3. AlphaFold3-based assessment of TRN1 interactions with predicted functional and non-functional PY-NLS motifs and their PY/AA mutated variants. A)** Paired boxplots of the AlphaFold3 combined confidence score (0.8×ipTM + 0.2×pTM; best-of-5 models) for wild-type (WT) and PY→AA point mutants (PY/AA) motif constructs in complex with TRN1. Left: 20 proteins predicted to bear functional PY-NLS motifs (blue/green) together with PY-NLS motifs from JAZ1 and PLA2-like . Right: 20 proteins predicted to bear non-functional PY-NLS motifs (orange/pink). Paired lines connect WT and PY/AA values for each protein. Proteins selected for experimental validation are labeled; PLA2-like and JAZ1 are highlighted in red. P-values: Mann-Whitney U (functional WT vs. non-functional WT, one-sided) and Wilcoxon signed-rank (WT vs. PY/AA within each group, one-sided). **B)** Per-residue ipSAE profile across the PY-NLS motif region, aligned at the tyrosine residue (position 0) of the PY dipeptide. Top panel: per-residue ipSAE values computed from best-of-5 models (mean ± SEM). Dashed vertical lines mark the P (−1) and Y (0) positions. Bottom panel: $-\log_{10}(p)$ for functional vs. non-functional comparison (black dashed, Mann-Whitney U, one-sided) and functional WT vs. PY/AA comparison (blue, Wilcoxon signed-rank, one-sided). Red dotted line indicates p = 0.05.

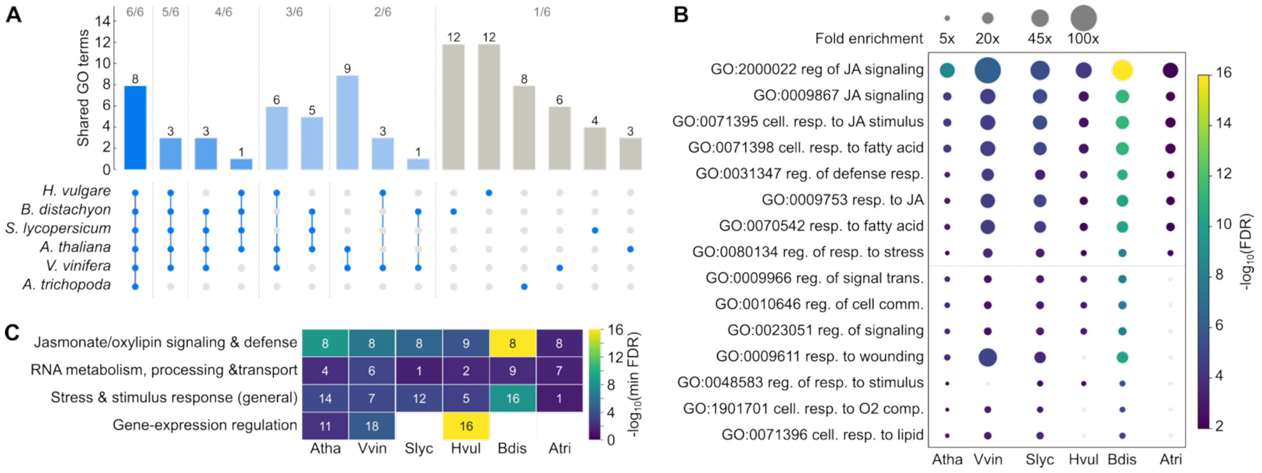


**Supplementary Figure 4. Conservation of Gene Ontology (GO) enrichment among high-confidence PY-NLS candidate proteins across angiosperms. A)** Upset plot showing the sharing of enriched GO terms (Biological process) among six angiosperm species. Bars give the number of terms in each exclusive intersection, grouped by the number of species sharing them (6/6 to 1/6); filled dots in the matrix below indicate the participating species for each intersection. **B)** Enrichment profiles of the 15 GO terms shared by at least four species. The dashed line separates GO terms shared by all tested species. **C)** Functional-theme summary across species. Color intensity represents $-\log_{10}$ of the minimum FDR observed for any term in the theme; numbers give the count of enriched GO terms per theme and species (see Supplementary Data 2 for the details on GO-term groupings into the four functional themes). Species abbreviations: Atha, *Arabidopsis thaliana*; Vvin, *Vitis vinifera*; Slyc, *Solanum lycopersicum*; Hvul, *Hordeum vulgare*; Bdis, *Brachypodium distachyon*; Atri, *Amborella trichopoda*.

### Supplementary Table 1. List of primers used in this study.

| Label | Name | 5'-3' Sequence | Description |
|---|---|---|---|
| PP868 | AtPLA2-like_ F_Ngo MIV | ataGCCGGCATGAATTTTGGGTTGCCAAG | pWEN vector FL_N-term YFP fusion, pTNT fusion |
| PP869 | AtPLA2-like _S-R_ApaI (Stop codon) | ataGGGCCCTCAGTTTTTCTGAAAACTCCAG | FL_N-term YFP fusion, pTNT Fusion |
| AS9 | AtPLA2-Like_NLS_F | ataGCCGGCAGAGGGTTAAAAAGGCATG | Truncated_N-term YFP fusion |
| AS10 | AtPLA2-Like-NoNLS_F | ataGCCGGCGTGTCTAAGGTTCCGTGGCAT | Truncated_N-term YFP fusion |
| AS19 | AtPLA2-Like_NLS_R | ataGGGCCCctaATAAGGTCTAAACCGATAAAC | PY-NLS N-term YFP fusion |
| AS21 | AtPLA2-Like_mutNLS_R | ataGGGCCCctaagcagcTCTagcCCGATAAACAGTATTCACACC | mutPY-NLS N-term YFP fusion |
| AS36 | AtPLA2-like_N-term_KR motif_F | CCG GCAGAGGGTTAAAAAGGCATGCTCTTTCTCGGAGGG GGC C | Short N-term of PY-NLS motif YFP fusion |
| AS37 | AtPLA2-like_N-term_KR motif_R | CCCTCCGAGAAAGAGCATGCCTTTTTAACCCTCTG | Short N-term of PY-NLS motif YFP fusion |
| AS49 | AtPLA2-like_C-term_PY motif_NgoMIV_ApaIR | CCGGCTCATCAAGATCAAATGGTGTGAATACTGTTTATCGGTTTAGACCTTATGTAGGGGCC | Short C-term of PY-NLS motif YFP fusion |
| AS50 | AtPLA2-like_C-term_PY motif_NgoMIV_ApaI | CCATAAGGTCTAAACCGATAAACAGTATTCACACCATTTGATCTTGATGAG | Short C-term of PY-NLS motif YFP fusion |
| AS53 | AtHistone2B_FL_XmaI_F | ataCCCGGGATGGCGCCGAGAGCAGAGAAGAAGC | FL_N-term YFP/mCherry fusion |
| AS54 | AtHistone2B_FL_ApaI_R | ataGGGCCCTCAAGAGCTTGTGAATTTGGTAACAGC | FL_N-term YFP/mCherry fusion |
| AS106 | PLA2-like-PYNLS1_NgoMIV_ApaI_F | ataGCCGGCAGAGGGTTAAAAAGGCATCGGTTTAGACCTTATTAGGGGCCCata | Linker 1, N-term YFP fusion |
| AS107 | PLA2-like-PYNLS1_NgoMIV_ApaI_R | tatGGGCCCCTAATAAGGTCTAAACCGATGCCTTTTTAACCCTCTGCCGGCtat | Linker 1, N-term YFP fusion |
| AS112 | PLA2-like-At_NLS4_NgoMIV_ApaI_F | ataGCCGGCAGAGGGTTAAAAAGGCATGCTCTTTCTCGGTTTAGACCTTATTAGGGGCCCata | Linker 4, N-term YFP fusion |
| AS113 | At_NLS4_NgoMIV_ApaI_R | tatGGGCCCCTAATAAGGTCTAAACCGAGAAAGAGCATGCCTTTTTAACCCTCTGCCGGCtat | Linker 4, N-term YFP fusion |
| AS114 | At_NLS8_NgoMIV_ApaI_F | ataGCCGGCAGAGGGTTAAAAAGGCATGCTCTTTCTCGGAGGTCATCACGGTTTAGACCTTATTAGGGGCCCata | Linker 8, N-term YFP fusion |
| AS115 | At_NLS8_NgoMIV_ApaI_R | tatGGGCCCCTAATAAGGTCTAAACCGTGATGACCTCCGAGAAAGAGCATGCCTTTTTAACCCTCTGCCGGCtat | Linker 8, N-term YFP fusion |
| AS116 | At_NLS13_NgoMIV_ApaI_F | ataGCCGGCAGAGGGTTAAAAAGGCATGCTCTTTCTCGGAGGTCATCAAGATCAAATGGTGTGCGGTTTAGACCTTATTAGG GGCCCata | Linker 13, N-term YFP fusion |
| AS117 | At_NLS13_NgoMIV_ApaI_R | tatGGGCCCCTAATAAGGTCTAAACCGCACACCATTTGATCTTGATGACCTCCGAGAAAGAGCATGCCTTTTTAACCCTCTG CCGGCtat | Linker 13, N-term YFP fusion |
| AS167 | At_NLS22_NgoMIV_ApaI_R | ataGCCGGCAGAGGGTTAAAAAGGCATGGGGCTGGGGCTGGGGCTCTTTCTCGGAGGTCATCAAGATCAAATGGTGTGAATA CTGTTTATCGGTTTAGACCTTATTAGGGGCCCata | Linker 22, N-term YFP fusion |
| AS168 | At_NLS22_NgoMIV_ApaI_R | tatGGGCCCCTAATAAGGTCTAAACCGATAAACAGTATTCACACCATTTGATCTTGATGACCTCCGAGAAAGAGCCCCAGCC CCAGCCCCATGCCTTTTTAACCCTCTGCCGGCtat | Linker 22, N-term YFP fusion |
| AS77 | M9MNLS_NgoMIV_ApaI_F | ataGCCGGCatgttcggcaactacaacaaccagtcgtcgaacttcggcccgatgaagggcggcaacttcggcggccgcttcg agccgtacTAGGGGCCCata | pWEN vector N-term YFP fusion |
| AS78 | M9MNLS_NgoMIV_ApaI_R | tatGGGCCCCTAgtacggctcgaagcggccgccgaagttgccgcccttcatcgggccgaagttcgacgactggttgttgtag ttgccgaacatGCCGGCtat | N-term fusion |

| | | | |
|---|---|---|---|
| AS178 | FL_JAZ1_NgoMIV_F | ataGCCGGCATGTCGAGTTCTATGGAATGT | FL_N-term YFP fusion, pTNT fusion |
| AS164 | FL_JAZ1_ApaI_R | ataGGGCCCTCATATTTCAGCTGCTAAACC | FL_N-term YFP fusion, pTNT fusion |
| AS173 | FL_WRKY8_NgoMIV_F | ataGCCGGCATGTCTCATGAAATCAAAGAT | FL_N-term YFP fusion |
| AS174 | FL_WRKY8_ApaI_R | ataGGGCCCTCAAGGCTCTTGTTTGAAGAAAAC | FL_N-term YFP fusion |
| AS183 | JAZ1_PY_NgoMIV_F | ataGCCGGCATTGCTAGAAGAGCTTCACTTCACCGGTTCTTGGAGAAGAGAAAGGACAGAGTTACGTCAAAGGCACCATACtagGGGCCCata | PY-NLS N-term YFP fusion |
| AS184 | JAZ1_PY_ApaI_R | tatGGGCCCCTAGTATGGTGCCTTTGACGTAACTCTGTCTTTCTCTTCTCCAAGAACCGGTGAAGTGAAGCTCTTCTAGCAATGCCGGCTAT | PY-NLS N-term YFP fusion |
| AS185 | JAZ1_mPYaa_NgoMIV_F | ataGCCGGCATTGCTAGAAGAGCTTCACTTCACCGGTTCTTGGAGAAGAGAAAGGACAGAGTTACGTCAAAGGCAgctgcttagGGGCCCata | PY-NLS N-term YFP fusion |
| AS186 | JAZ1_mPYaa_ApaI_R | tatGGGCCCctaagcagcTGCCTTTGACGTAACTCTGTCCTTTCTCTTCTCCAAGAACCGGTGAAGTGAAGCTCTTCTAGCAATGCCGGCTAT | PY-NLS N-term YFP fusion |
| AS191 | WRKY8_PY_NgoMIV_F | ataGCCGGCatgCGTTGGAGAAAGTATGGCCAAAAAGCAGTCAAAAACAGTCCTTATTAGGGGCCCata | PY-NLS N-term YFP fusion |
| AS192 | WRKY8_PY_ApaI_R | tatGGGCCCCTAATAAGGACTGTTTTTGACTGCTTTTTGGCCATACTTTCTCCAACGcatGCCGGCtat | PY-NLS N-term YFP fusion |
| AS355 | WRKY8 Pynls_megaprimer PYaa_F | AACAGTGCAGCACCGAGGAGTTACTATAGATGC | pWEN N-ter YFP fusion |
| AS193 | megaprimer_FL_AtPLA2_PYAA_ApaI_R | CCACGGAACCTTAGACACCGCCGCTCTAAACCG | FL_mutPY N-term YFP fusion |
| AS194 | megaprimer_FL_ATJAZ1_PYAA_NgoMIV_F | ACGTCAAAGGCAGCAGCACAATTATGCGAT | FL_mutPY N-term YFP fusion |
| AS82 | AtTRN1_EcoRI_FL_F | ataGAATTCATGGCGGCGACGGCGGTGGTCTGGC | Y2H_BD fusion |
| AS83 | AtTRN1_SalI_FL_R | ataGTCGACTTACACTTGATATCTCGCAAGCCTTTC | Y2H_BD fusion |
| AS152 | M9MNLS_Y2H EcoRI_ BamHI | ataGAATTCataGCCGGCatgttcggcaactacaacaaccagtcgtcgaacttcggcccgatgaagggcggcaacttcggcggccgcttcgagccgtacTAGGGATCCata | Y2H AD fusion |
| AS153 | M9MNLS_Y2H_EcoRI_BamHI | tatGGATCCCTAgtacggctcgaagcggccgccgaagttgccgcccttcatcgggccgaagttcgacgactggttgttgtagttgccgaacatGCCGGCtatGAATTCtat | Y2H AD fusion |
| AS92 | AtPLA2-L_Y2H_EcoRI_F | ataGAATTCATGAATTTTGGGTTGCCAAGT | Y2H_AD fusion |
| AS93 | AtPLA2-L_Y2H_BamHI_R | ataGGATCCGTTTTTCTGAAAACTCCAGCT | Y2H_AD fusion |
| AS200 | JAZ1_Y2H_EcoRI_F | ataGAATTCATGTCGAGTTCTATGGAATGT | Y2H_AD fusion |
| AS201 | JAZ1_Y2H_BamHI_R | ataGGATCCTATTTCAGCTGCTAAACCGAGCCA | Y2H_AD fusion |
| AS351 | JAZ1 Pynls_RRaa_F Epi1 | ataGCCGGCatgGCAGCAGCTTCACTTCACCGGTTCTTGGAGAAGAGAAAGGACAGAGTTACGTCAAAGGCACCATACtgaGGGCCCata | pWEN N-ter YFP fusion |
| AS352 | JAZ1 Pynls_RRaa_R Epi1 | tatGGGCCCtcaGTATGGTGCCTTTGACGTAACTCTGTCCTTTCTCTTCTCCAAGAACCGGTGAAGTGAAGCTGCTGCcatGCCGGCtat | pWEN N-ter YFP fusion |
| AS353 | JAZ1 Pynls_Ka_F Epi2 | ataGCCGGCatgAGAAGAGCTTCACTTCACCGGTTCTTGGAGAAGAGAAAGGACAGAGTTACGTCAGCAGCACCATACtgaGGGCCCata | pWEN N-ter YFP fusion |
| AS354 | AZ1 Pynls_Ka_R Epi2 | tatGGGCCCtcaGTATGGTGCTGCTGACGTAACTCTGTCCTTTCTCTTCTCCAAGAACCGGTGAAGTGAAGCTCTTCTcatGCCGGCtat | pWEN N-ter YFP fusion |
| AS264 | ΔJas_JAZ1_ApaI_R | ataGGGCCCCTAAGCAATAGGAAGTTCTGTCAATGG | pWEN N-ter YFP fusion |
| AS265 | ChimeraJAZ1_PL_ApaI_R | ataGGGCCCCTAATAAGGTCTAAACCGATAAACAGTATTCACACCATTTGATCTTGATGACCTCCGAGAAAGAGCATGCCTTTTTAACCCTCTAGCAATAGGAAGTTCTGTCAATGG | ChimeraJAZ1_PL_ApaI_R |
| AS266 | AtPLA2-like-2xHA_KpnI_F | ataggtaccatgtacccatacgatgttccagattacgcttacccatacgatgttccagattacgctATGAATTTTGGGTTGCCAAGTATATCTTGG | pWEN N-ter fragment clone into pHD71 vector for agrobacterium |
| AS267 | AtPLA2-like-YFP-EcoRI_R | ataGAATTCTTACTTGTACAGCTCGTCCATGCC | pWEN N-ter fragment clone into pHD71 vector for agrobacterium |

| | | | |
|---|---|---|---|
| AS273 | JAZ1-PLA2-PYNLS_AgeI_F | ataACCGGTAGAGGGTTAAAAAGGCATGCT | pWEN N-ter YFP fusion |
| AS274 | JAZ1-PLA2-PYNLS_NotI_R | ataGCGGCCGCttTCAATAAGGTCTAAACCGATAAAC | pWEN N-ter YFP fusion |
| AS217 | FL_PHE1(MAD box)_NgoMIV_F | ataGCCGGCATGAGGGGGAAGATGAAGTTATCG | pWEN N-ter YFP fusion |
| AS218 | FL_PHE1(MAD box)_ApaI_R | ataGGGCCCCTATAAATCATTGATGATGTTAGGAGC | pWEN N-ter YFP fusion |
| AS358 | PHE1(MAD box)_megaprimer | CTGGATCGAGTTCGCCGCGCTACGGATGAC | pWEN N-ter YFP fusion |
| AS207 | JAZ1 short PY motif_F Ngo ApaI | ataGCCGGCatgAAGAGAAAGGACAGAGTTACGTCAAAGGCACCATACTAGGGGCCCata | pWEN N-ter YFP fusion |
| AS208 | JAZ1 short PY motif_R Ngo ApaI | tatGGGCCCCTAGTATGGTGCCTTTGACGTAACTCTGTCCTTTCTCTTcatGCCGGCtat | pWEN N-ter YFP fusion |
| AS396 | FL_IAA34_NgoMIV_F | ataGCCGGCatgtattgcagcgatcctccccatccc | pWEN N-ter YFP fusion |
| AS397_ | FL_IAA34_ApaI_R | tatGGGCCCttaaaagggaagtacagcatcgtttct | pWEN N-ter YFP fusion |
| AS398 | megaPrimer _IAA34_PYaa _R | accaaactccgtggtctgcgagtaggcggcatgatacttgggagaatttgg | pWEN N-ter YFP fusion |
| AS399 | megaprimer__BZR1_PYaa_R | agggctctggttctgtgatgaggcggcagttactcgagatgaagtccc | pWEN N-ter YFP fusion |
| AS400 | FL_BZR1_NgoMIV_F | ataGCCGGCatgacttcggatggagctacgtcg | pWEN N-ter YFP fusion |
| AS401 | FL_BZR1_ApaI_R | tatGGGCCCtcaaccacgagccttcccatttcc | pWEN N-ter YFP fusion |
| AS402 | FL_ZnGIS_NgoMIV_F | ataGCCGGCatggacgaggctaccggagaaaca | pWEN N-ter YFP fusion |
| AS403 | FL_ZnGIS_ApaI_R | tatGGGCCCttaaagatgaagatcgagactcac | pWEN N-ter YFP fusion |
| AS404 | ZnGIS megaprimer__PYaa_F | tacttttatcatcctgacaataacgctGCgagttaccgtcattacccgtcgtgg | pWEN N-ter YFP fusion |
| AS405 | FL_SKIP5_NgoMIV_F | ataGCCGGCatggagttacatgagatctcaaag | pWEN N-ter YFP fusion |
| AS406 | FL_SKIP5_ApaI_R | tatGGGCCCtcattcataatctacatcaaacca | pWEN N-ter YFP fusion |
| AS407 | megaprimer_SKIP5 _PYaa_R | gagattgttaagagacgtgagagatgcagctttcattctcttcttcatcttcga | pWEN N-ter YFP fusion |
| AS408 | FL_PABN2_NgoMIV_F | ataGCCGGCatggaggaagaggagcacgaggtt | pWEN N-ter YFP fusion |
| AS409 | FL_PABN2_ApaI_R | tatGGGCCCttattggtacggcatgtaacgcatagg | pWEN N-ter YFP fusion |
| AS410 | Δ Cter_PABN2_ApaI_R | tatGGGCCCttactgcaagacctttaattgacgacc | pWEN N-ter YFP fusion |
| AS368 | SALK_003127_LP | CATTGGCAAGAACTAGCCTTG | TRN1 genotyping |
| AS369 | SALK_003127_RP | TCTTAACGCAACCAATCCAAC | TRN1 genotyping |
| LBb1.3 | LBb1.3 | ATTTTGCCGATTTCGGAAC | T-DNA genotyping |
| AS367 | FL_TRN1_NgoMIV F | ataGCCGGCATGGCGGCGACGGCGGTGGTCTGGC | Pull-down assay cloning |

**Supplementary Table 2. Calibrated parameters of the pynls v1 scoring model.** All values are read at run time from pynls_v1_calibration.json (Supplementary Data 1). The weights of the five graded rule scores are applied as a weighted mean over the components with valid values and sum to 1.0.

| Parameter | Value | Role |
|---|---|---|
| **Weights of the graded rule scores** | | |
| w(S_sequence) | 0.120 | Linker-geometry term |
| w(S_disorder) | 0.259 | Consensus intrinsic-disorder term |
| w(S_structure) | 0.311 | DSSP β-strand compatibility term |
| w(S_exposure) | 0.104 | Relative solvent accessibility term |
| w(S_pLDDT) | 0.207 | AlphaFold local-confidence term |
| w_af3 | 0.100 | Weight of the AlphaFold3 interface term in the fused score |
| **Motif geometry** | | |
| linker1_target / linker1_sigma | 17 / 5 aa | Gaussian optimum and width for linker1 |
| linker2_target / linker2_sigma | 3 / 2 aa | Gaussian optimum and width for linker2 |
| linker1_max_strict | 20 aa | Upper bound on linker1 in the consensus regex |
| hydrophobic_downstream_warn | 3 residues | Hydrophobic residues within PY+1…PY+5 that trigger the 0.5 sequence penalty |
| **Structure and accessibility** | | |
| plddt_lo / plddt_hi | 60 / 85 | Bounds of the S_pLDDT ramp; pLDDT < 60 also masks DSSP calls to coil |
| rsa_mean_target | 25% | Reference mean motif RSA |
| rsa_min_target | 10% | Reference minimum motif RSA |
| iupred3_rescue_bonus | +0.15 | Disorder bonus when 60 ≤ pLDDT < 85 and IUPred3 ≥ 0.5 |
| **Gates** | | |
| topology_linker_threshold | 50 aa | Minimum permitted motif distance to the nearest transmembrane helix |
| topology_polytopic_min_loop | 50 aa | Minimum inter-helical loop length that may accommodate a motif |
| topology_soft_multiplier | 0.5 | S_topology of a topology-compliant membrane protein |
| cNLS Mapper cut-off | 5.0; region “Entire region” | Threshold and search mode for classical-NLS detection; S_cnls = 0.85 if reached |
| **Tier cut-offs (fused score)** | | |
| T1 / T2 / T3 | ≥ 0.65 / ≥ 0.45 / ≥ 0.25 | Confidence tiers; T4 otherwise, or on any hard veto |

### SUPPLEMENTARY METHODS

#### Genome-wide identification of PY-NLS candidates (Python script pynls v1)

##### *Implementation and availability*

The PY-NLS prediction pipeline is a single, self-contained command-line program written in Python 3.10 (pynls_v1.py, 2,448 lines), provided in Supplementary Data 1. Every calibrated constant — component weights, tier cut-offs, Gaussian targets and widths, and veto thresholds — is read at runtime from an external JSON file (pynls_v1_calibration.json). The complete parameter set is reproduced in Supplementary Table 2. Third-party Python dependencies are *numpy*, *pandas*, *scipy*, *requests*, *biopython*, *freesasa* and *metapredict*; the only external binary required is the DSSP implementation *mkdssp* (v4.6.1).

##### *Input modes and proteome retrieval*

The pipeline accepts three mutually exclusive inputs: a local FASTA file (--fasta), a list of UniProt accessions (--ids), or an organism name (--organism). In organism mode the species name is resolved against the UniProt Proteomes API restricted to proteome_type:REFERENCE, and the canonical proteome FASTA (Swiss-Prot plus TrEMBL) is streamed from the UniProt REST interface (UniProt Consortium, 2025). The six angiosperm reference proteomes analysed here were *Arabidopsis thaliana* UP000006548, *Solanum lycopersicum* UP000004994, *Vitis vinifera* UP000009183, *Hordeum vulgare* subsp. *vulgare* UP000011116, *Brachypodium distachyon* UP000008810 and *Amborella trichopoda* UP000017836. For every candidate protein the corresponding UniProt entry is additionally fetched in JSON format to recover gene names, protein names and the sequence features used by the veto terms (signal peptide, transit peptide, propeptide and transmembrane helix).

All computationally expensive steps — structure download, DSSP, solvent-accessibility and disorder calculation, and cNLS Mapper queries — are applied only to proteins containing at least one consensus motif. This reduces the structural workload by roughly 95%: in *A. thaliana*, 1,490 genes of the approximately 27,400 protein-coding genes carry a motif and are carried forward. Per-protein results and downloaded structures are cached on disk, so that interrupted runs resume without recomputation and repeated analyses of overlapping gene sets are inexpensive.

##### *Consensus motif definition*

Candidate PY-NLS motifs are located with the regular expression ([KR][KR].{3,20}[KRH].{1,5}PY))

The pattern is wrapped in a zero-width lookahead so that overlapping and nested matches within the same protein are all recovered, rather than only the leftmost non-overlapping match; a single protein therefore frequently yields several candidate motifs, which are subsequently ranked and collapsed.

#### *Protein feature extraction*

Each motif-containing protein is annotated with the following independent evidence layers.

Structural models: The predicted monomer model is downloaded from the AlphaFold Protein Structure Database (Jumper et al., 2021; Varadi et al., 2024), trying the current model release first and falling back to earlier versions. Accessions with no deposited model are substituted by a sequence-identical entry located through UniRef100 (Suzek et al., 2015) and, failing that, through UniParc; every substitution is recorded in the af_substitute_uid column of Supplementary Data 2, so that the provenance of each structure-derived feature remains traceable. Downloaded coordinate files are sanitised (DBREF records rewritten) because *mkdssp* rejects identifiers longer than six characters.

Intrinsic disorder: Per-residue disorder is predicted with AIUPred (Erdős and Dosztányi, 2024), metapredict (Emenecker et al., 2021) and IUPred3 (Erdős et al., 2021), the last accessed through its REST service. Predictions are made on the full-length sequence and summarised over the motif window as the mean, minimum, maximum and the fraction of motif residues scoring $\geq$ 0.5. AIUPred binding- and linker-propensity tracks are recorded alongside the disorder track but do not enter the score.

Secondary structure: DSSP assignments (Kabsch and Sander, 1983) are computed from the AlphaFold model with *mkdssp* v4.6.1 (Touw et al., 2015). Residues with pLDDT $< 60$ are masked to coil before the assignment is evaluated, because AlphaFold treats such regions as effectively disordered and secondary-structure calls within them are not informative.

Solvent accessibility**:** Absolute solvent-accessible surface areas are computed with FreeSASA (Mitternacht, 2016) and converted to relative solvent accessibility (RSA) using the maximum-ASA reference values of Tien et al. (2013). RSA is summarised over the whole motif and, separately, over the PY core dipeptide.

Localization annotation and membrane topology**:** Signal-peptide, transit-peptide, propeptide and transmembrane-helix spans are taken from the UniProt entry (UniProt Consortium, 2025) and converted into the annotation and topology gates defined below. No de novo topology prediction is performed, so proteins with incomplete UniProt feature annotation are treated as having no transmembrane segment.

Classical NLS co-occurrence: Each protein sequence is submitted to the cNLS Mapper web service (Kosugi et al., 2009b) with a prediction cut-off score of 5.0 and the search region set to *Entire region*, so that both monopartite and bipartite classical NLSs anywhere in the protein are reported. The highest monopartite and bipartite scores are retained separately (cNLS_mp_max_score, cNLS_bp_max_score) and a protein is called cNLS-positive if either reaches the cut-off.

Receptor-interface evidence (optional): When AlphaFold3 co-folding output for the protein together with TRN1 is supplied, the pipeline parses interface pSAE values (ipSAE; Dunbrack, 2025) alongside ipTM and pTM and converts them into an additional score component. Note, that this evidence was not used in the genome-wide runs reported in Supplementary Data 2, Supplementary Fig 3, 4.

#### *Component scores*

Nine component scores are computed for each motif, all bounded to [0, 1]. Five are graded rule scores that enter a weighted mean; four are multiplicative gates. A component that cannot be computed — for example because no structural model is available — is left empty and excluded from the mean rather than being imputed.

#### *Graded rule scores*

$S_{sequence}$ — linker geometry: Each linker length is scored by a Gaussian centred on the experimentally preferred length:

```
l1 = exp( −(len(linker1) − 17)² / (2 × 5²) )
l2 = exp( −(len(linker2) −  3)² / (2 × 2²) )
S_sequence = clip(0.6·l1 + 0.4·l2, 0, 1) × d
```

Linker1 receives the larger weight because it is the stronger determinant of import activity in our mutagenesis series. The downstream penalty $d$ equals 0.5 when at least three hydrophobic residues (A, V, I, L, M, F, W, Y or C) fall within the five positions immediately following the PY dipeptide, and 1.0 otherwise; it flags motifs embedded in hydrophobic, potentially membrane-associated or aggregation-prone context.

$S_{disorder}$ — intrinsic disorder: The maximum of the mean motif disorder scores available from AIUPred, metapredict and IUPred3, so that one confident predictor suffices to support flexibility. A rescue bonus of +0.15 is added when the motif has ambiguous structural confidence (60 ≤ mean pLDDT < 85) but IUPred3 nevertheless reports ≥ 0.5; the sum is re-clipped to [0, 1] and application of the bonus is recorded in iupred3_rescue_applied.

$S_{structure}$ — secondary-structure compatibility: Assigned from the masked DSSP string across the motif window: 1.00 when no β-strand residue is present (strict pass); 0.70 when at most one β-strand residue is present (relaxed pass); 0.45 for a single isolated β-strand call, treated as possible DSSP noise (borderline); and 0.15 when two or more consecutive β-strand residues are present, a rigid arrangement incompatible with the extended conformation adopted by PY-NLS peptides in the karyopherin-β2 groove (Lee et al., 2006). The 0.15 case is also recorded as a gate failure for tier assignment.

$S_{exposure}$ — surface accessibility: A weighted combination of motif RSA terms, with RSA in per cent:

```
S_exposure = clip( 0.4·(RSA_mean / 25) + 0.5·(RSA_PY_core / 25) + 0.1·m , 0, 1 )
```

where $m$ = 1.0 if the minimum motif RSA is ≥ 10% and 0.5 otherwise. The PY core carries the largest weight because it makes the most deeply buried contacts with the receptor.

$S_{pLDDT}$ — model confidence as a disorder proxy: Low local model confidence is favourable for an accessible linear motif:

S_pLDDT = 1.0 − 0.8 × clip( (mean pLDDT − 60) / (85 − 60), 0, 1 )

This gives 1.0 below pLDDT 60, 0.2 at or above pLDDT 85, and a linear ramp between. $S_{pLDDT}$ and $S_{disorder}$ are deliberately retained as separate terms because they draw on independent evidence (a structural model versus sequence-based predictors) and disagree in an informative minority of cases.

#### *Multiplicative gates*

$S_{annotation}$: Zero if the protein carries a signal-peptide, transit-peptide or propeptide annotation, indicating a secretory or organellar destination incompatible with nuclear function; otherwise 1.0. In *A. thaliana*, 242 of the 1,490 motif-containing genes are vetoed on this basis (128 signal peptide, 82 transit peptide, 5 propeptide, and 27 in combination with a topology veto).

$S_{topology}$: Evaluated from the annotated transmembrane spans relative to the motif position. Proteins with no annotated transmembrane helix score 1.0. A hard veto of zero is applied when the motif overlaps a transmembrane helix, when it lies in an inter-helical loop shorter than 50 residues, or when it lies within 50 residues of the nearest transmembrane helix. Membrane proteins whose motif satisfies all three distance criteria receive 0.5 — a deliberate downweight that retains them as plausible but lower-priority candidates instead of discarding them, since a membrane-tethered PY-NLS can be released by proteolysis. In *A. thaliana* this partitions the 1,490 genes into 1,249 with no transmembrane annotation (gate 1.0), 143 topology-compliant membrane proteins (gate 0.5) and 98 hard vetoes (gate 0.0: 66 motif too close to a helix, 20 motif inside a helix, 12 motif in a short inter-helical loop).

$S_{cNLS}$: 0.85 if cNLS Mapper reports a classical NLS at or above the cut-off, reflecting possible redundancy with the importin-α/β route; otherwise 1.0. This is a soft penalty only — cNLS-positive proteins are retained and ranked throughout, and are removed only at the final candidate-selection step (Supplementary Methods 1.7).

$S_{af3}$: clip(mean $ipSAE_{d0chn}$ / 0.25, 0, 1) when AlphaFold3 co-folding evidence is available; not applicable to the genome-wide runs reported here.

#### *Score fusion and tier assignment*

The five graded scores are combined as a weighted mean over the components that could be computed, and the result is then gated multiplicatively:

```
S_rules       = Σ(wᵢ·Sᵢ) / Σ(wᵢ)      over the five graded scores with valid values
S_rules_gated = S_rules × S_annotation × S_topology × S_cnls
S_af3_gated   = S_af3   × S_annotation × S_topology
fused         = 0.1·S_af3_gated + 0.9·S_rules_gated   (AlphaFold3 evidence present)
              = S_rules_gated                          (otherwise)
```

The ungated weighted mean is reported alongside it (ungated_fused_score) so that the contribution of the gates to any individual ranking can be inspected.

Candidates are then binned into four confidence tiers: T1, fused ≥ 0.65; T2, fused ≥ 0.45; T3, fused ≥ 0.25; and T4, fused < 0.25 or any hard veto ($S_{annotation} = 0$ or $S_{topology} = 0$) or failure of the secondary-structure gate.

Because the lookahead regex returns overlapping matches, each protein is first collapsed to its single best motif by fused score. Proteins are then collapsed to genes using the UniProt primary gene name where available, falling back to the first listed gene synonym and finally to the accession; within a gene, ties are broken in favour of the longer sequence and then by alphabetical accession. All rows of Supplementary Data 2 are consequently gene-level, one row per gene, with the number of collapsed protein entries and their accessions retained in n_proteins_per_gene and all_protein_ids.

***Calibration***

Component weights were set from the experimental measurements. The relative weights of the five graded components (Supplementary Table 2) place the greatest emphasis on secondary-structure compatibility and on model confidence, consistent with the requirement that a PY-NLS be presented in an extended, flexible conformation, and the smallest on linker geometry, which serves mainly to discriminate between competing motifs within the same protein. Tier cut-offs were chosen so that T1 corresponds to candidates in which all graded criteria are simultaneously favorable. Because the weights derive from the same experimental series used elsewhere in this study, the tier boundaries should be read as an internally consistent ranking device rather than as calibrated probabilities, and prospective performance on independent cargoes remains to be established.

***Outputs***

A run writes four files: pynls_v1_hits.tsv (every motif in every motif-containing protein, with all features and component scores), pynls_v1_summary.tsv (one row per protein, best motif only), pynls_v1_by_gene.tsv (one row per gene) and pynls_v1_run.log (a run record file including the resolved proteome identifier, per-stage counts and all failed retrievals).

**Structural modelling of TRN1–cargo complexes and interface scoring**

The complexes shown in Fig. 3 were predicted with AlphaFold2-Multimer (Evans et al., 2021; Jumper et al., 2021). The complexes used for the quantitative interface comparison in Fig. 4 and Supplementary Fig. 3 were predicted with AlphaFold3 (Abramson et al., 2024). In both cases the *A. thaliana* TRN1/IMB2 sequence (AT2G16950) was co-folded with the cargo sequence or with the isolated motif peptide, and five models were generated per complex.

Models were ranked by the composite confidence measure 0.8 × ipTM + 0.2 × pTM, which tracks the accuracy of predicted interactions involving intrinsically disordered regions better than either term alone (Omidi et al., 2024), and the best-scoring of the five models was retained for each complex. Because ipTM is computed over all inter-chain residue pairs, it is systematically diluted when a short peptide is docked onto a large receptor; interfaces were therefore additionally scored with ipSAE (Dunbrack, 2025), an interface-restricted reformulation of the same underlying predicted aligned error. ipSAE values reported here are the chain-normalised variant ($ipSAE_{d0chn}$) averaged across the five models, with per-model maxima retained for reference. Structures were inspected and rendered in UCSF ChimeraX (Meng et al., 2023).

**Functional enrichment of PY-NLS candidate datasets and species comparison**

Gene Ontology Biological Process enrichment of the high-confidence candidate sets was computed with ShinyGO v0.85 (Ge et al., 2020) using each species' own genome annotation as the statistical background, with significance expressed as a Benjamini–Hochberg false-discovery rate and a cut-off of FDR $\leq$ 0.05. All 183 enriched terms reported across the six species satisfy this threshold. The complete enriched-term tables for all six species, as exported from ShinyGO, are provided in the Suppl_Data_S2-GOBP_enrichment sheet of Supplementary Data 2.

Overlaps between candidate sets and previously reported TRN1-associated protein lists were computed and drawn with the Venn diagram web tool of the VIB–UGent Center for Plant Systems Biology (https://bioinformatics.psb.ugent.be/webtools/Venn/).

### SUPPLEMENTARY DATA

#### Supplementary Data 1

A compressed archive containing the complete prediction code and its documentation. Four files are included:

- pynls_v1.py — the entire pipeline as one executable Python 3 script.
- pynls_v1_calibration.json — the external parameter file holding all component weights; reproduced in full as Supplementary Table 1.
- pynls_v1_README.md — installation and dependency notes.
- pynls_v1_scoring_description.md — a narrative summary of the motif definition, the nine component scores, the fusion formula and the tier criteria.

#### Supplementary Data 2

Gene-level PY-NLS predictions for six angiosperm reference proteomes, together with the Gene Ontology Biological Process enrichment of the candidate set in each species.
