## Supplementary Data 1 for "Sequence and structural rules define PY-NLS–dependent nuclear import by TRANSPORTIN 1 in plants": pynls_v1.py

```
#!/usr/bin/env python3
"""
pynls_v1.py — basic PY-NLS motif prediction pipeline for plants.

Fresh implementation steered by the seven biology rules (a)-(g) and an
AF3-with-TRN1 reference dataset.

Three input modes:
  --fasta      FASTA file with protein sequences
  --ids        newline/comma-separated UniProt accession list
  --organism   organism name; downloads UniProt reference proteome (canonical)

Outputs (in --out-dir, default ./pynls_v1_out/):
  pynls_v1_hits.tsv       one row per (uniprot_id, motif) with all features + tier
  pynls_v1_summary.tsv    one row per protein (best motif) + counts
  pynls_v1_run.log        per-protein retrieval + processing log

Pipeline (per candidate protein):
  A. Regex scan            [KR][KR].{3,20}[KRH].{1,5}PY (overlapping, all hits)
  B. Sequence features     linker1/2 length, hydrophobic flanking, PY core
  C. Structure             AlphaFold monomer (EBI DB), mkdssp SSE, FreeSASA RSA
  D. Disorder              AIUPred (local CLI, primary) + metapredict + IUPred3 (REST fallback)
  E. UniProt annotation    TM, signal, transit, propeptide
  F. cNLS Mapper           monopartite + bipartite
  G. Topology gate         plant default: linker >=50 aa from TM; motif not in TM
  H. AF3 (opportunistic)   parse candidate-TRN1 ipSAE if --af3-dir supplied
  I. Component scores      7 orthogonal [0,1] scores
  J. Fused score + Tier    calibrated on AF-214, gated on annotation/topology

Usage
-----
  python pynls_v1.py --ids ids.txt --out-dir out/
  python pynls_v1.py --fasta seqs.fa --out-dir out/
  python pynls_v1.py --organism "Arabidopsis thaliana" --out-dir out/
  python pynls_v1.py --ids ids.txt --af3-dir /path/to/af3/ --out-dir out/

"""
from __future__ import annotations

import argparse
import concurrent.futures as cfut
import hashlib
import io
import json
import logging
import os
import re
import shutil
import subprocess
import sys
import tempfile
import time
from dataclasses import dataclass, field, asdict
from pathlib import Path

import numpy as np
import pandas as pd
import requests
from tqdm import tqdm

__version__ = "1.0"

# ==========================================================================
# CONSTANTS
# ==========================================================================

# Primary consensus regex (rule a): [KR][KR] {3-20 spacer} [KRH] {1-5 spacer} PY
# Lookahead form yields overlapping matches.
MOTIF_REGEX = re.compile(r'(?=([KR][KR].{3,20}[KRH].{1,5}PY))')

# Amino-acid classifications.
AA3_TO_1 = {
    'ALA': 'A', 'ARG': 'R', 'ASN': 'N', 'ASP': 'D', 'CYS': 'C', 'GLN': 'Q',
    'GLU': 'E', 'GLY': 'G', 'HIS': 'H', 'ILE': 'I', 'LEU': 'L', 'LYS': 'K',
    'MET': 'M', 'PHE': 'F', 'PRO': 'P', 'SER': 'S', 'THR': 'T', 'TRP': 'W',
    'TYR': 'Y', 'VAL': 'V',
}
HYDROPHOBIC = set('AVILMFWYC')  # standard plus C (borderline)
# Tien 2013 empirical Gly-X-Gly maximum ASA (Å²), used for RSA (%) calculation.
TIEN_MAX_ASA = {
    'A': 129, 'R': 274, 'N': 195, 'D': 193, 'C': 167, 'Q': 225, 'E': 223, 'G': 104,
    'H': 224, 'I': 197, 'L': 201, 'K': 236, 'M': 224, 'F': 240, 'P': 159, 'S': 155,
    'T': 172, 'W': 285, 'Y': 263, 'V': 174,
}

# External endpoints
AF_URL_TEMPLATE = "https://alphafold.ebi.ac.uk/files/AF-{uid}-F1-model_v{v}.pdb"
AF_VERSIONS = (6, 5, 4, 3, 2, 1)
UNIPROT_ENTRY_URL = "https://rest.uniprot.org/uniprotkb/search"
UNIPROT_PROTEOME_LOOKUP_URL = "https://rest.uniprot.org/proteomes/search"
UNIPROT_STREAM_URL = "https://rest.uniprot.org/uniprotkb/stream"
UNISAVE_URL = "https://rest.uniprot.org/unisave/{}?format=fasta&versions=1"
UNIPARC_SEARCH_URL = "https://rest.uniprot.org/uniparc/search"
IUPRED3_BASE = "https://iupred3.elte.hu"
IUPRED3_API_URL = f"{IUPRED3_BASE}/plot_api"
CNLS_MAPPER_URL = "http://nls-mapper.iab.keio.ac.jp/cgi-bin/NLS_Mapper_y.cgi"

_TRANSMEM_RE = re.compile(r'TRANSMEM\s+(\d+)\.\.(\d+)')
_SIGNAL_RE   = re.compile(r'SIGNAL\s+(\d+)\.\.(\d+)')

# Default scoring weights and thresholds. These conservative starting values
# can be overridden by a calibration JSON file.
DEFAULT_CALIBRATION = {
    "rule_weights": {
        "S_sequence":   0.15,
        "S_disorder":   0.30,
        "S_structure":  0.20,
        "S_exposure":   0.20,
        "S_pLDDT":      0.15,  # standalone pLDDT signal (low = disordered = good)
    },
    "w_af3": 0.40,
    "tier_cuts": {"T1": 0.65, "T2": 0.45, "T3": 0.25},
    "linker1_target": 17.0,
    "linker1_sigma": 6.0,
    "linker2_target": 3.0,
    "linker2_sigma": 2.0,
    "plddt_hi": 85.0,
    "plddt_lo": 60.0,
    "iupred3_rescue_bonus": 0.15,
    "rsa_mean_target": 25.0,   # rule e: mean >=25
    "rsa_min_target": 10.0,    # rule e: min >=10
    "hydrophobic_downstream_warn": 3,  # improvement #3
    "topology_linker_threshold": 50,   # plants (rule g)
    "topology_polytopic_min_loop": 50,
    "topology_soft_multiplier": 0.5,
    "linker1_max_strict": 17,          # for --strict-regex
    "used_linker1_term": True,
    "calibration_note": "default seed values"
}

# ==========================================================================
# LOGGING
# ==========================================================================

def setup_logging(logfile: Path, verbose: bool) -> logging.Logger:
    logger = logging.getLogger("pynls_v1")
    logger.setLevel(logging.DEBUG if verbose else logging.INFO)
    logger.handlers.clear()
    fmt = logging.Formatter("%(asctime)s %(levelname)s %(message)s", "%H:%M:%S")
    fh = logging.FileHandler(logfile, mode="w"); fh.setFormatter(fmt); fh.setLevel(logging.DEBUG)
    sh = logging.StreamHandler(sys.stderr); sh.setFormatter(fmt); sh.setLevel(logging.INFO if not verbose else logging.DEBUG)
    logger.addHandler(fh); logger.addHandler(sh)
    return logger


# ==========================================================================
# INPUT ADAPTERS  (rule preface)
# ==========================================================================

def read_fasta(path: str) -> dict[str, str]:
    """Parse a FASTA file, return {header_id: sequence}."""
    out = {}
    cur, seq = None, []
    with open(path) as f:
        for line in f:
            line = line.rstrip("\n").rstrip("\r")
            if not line:
                continue
            if line.startswith(">"):
                if cur is not None:
                    out[cur] = "".join(seq)
                # Take the first whitespace token; strip potential '|sp|ACC|NAME'
                hdr = line[1:].split()[0]
                # UniProt-style header like sp|P12345|NAME
                if "|" in hdr:
                    parts = hdr.split("|")
                    if len(parts) >= 3 and parts[0] in ("sp", "tr"):
                        cur = parts[1]
                    else:
                        cur = parts[0]
                else:
                    cur = hdr
                seq = []
            else:
                seq.append(re.sub(r'[^A-Za-z]', '', line).upper())
        if cur is not None:
            out[cur] = "".join(seq)
    return out


def parse_id_list(path: str) -> list[str]:
    """Parse a comma/newline-separated file of UniProt IDs."""
    with open(path) as f:
        raw = f.read()
    ids = re.split(r'[\s,;]+', raw)
    return [i.strip().upper() for i in ids if i.strip()]


def fetch_uniprot_sequences(uids: list[str], logger: logging.Logger,
                            batch_size: int = 100) -> dict[str, str]:
    """Fetch canonical sequences from UniProt for a batch of accessions."""
    out = {}
    uids = [u.upper() for u in uids]
    logger.info(f"Fetching {len(uids)} sequences from UniProt")
    for i in range(0, len(uids), batch_size):
        batch = uids[i:i+batch_size]
        query = " OR ".join(f"accession:{u}" for u in batch)
        params = {"query": query, "format": "fasta", "size": 500}
        try:
            r = requests.get(UNIPROT_ENTRY_URL, params=params, timeout=120)
            r.raise_for_status()
            # Parse the returned FASTA
            for header, seq in _parse_fasta_text(r.text).items():
                # UniProt returns headers like 'sp|ACCESSION|ENTRY_NAME ...'
                for token in header.split("|"):
                    tu = token.strip().upper()
                    if tu in batch:
                        out[tu] = seq
                        break
        except Exception as e:
            logger.warning(f"UniProt batch fetch failed: {e}")
        time.sleep(0.1)
    # Fallback: try individually via unisave for any missing (deleted entries).
    missing = [u for u in uids if u not in out]
    for u in missing:
        try:
            r = requests.get(UNISAVE_URL.format(u), timeout=60)
            if r.status_code == 200 and r.text.startswith(">"):
                seq = "".join(line for line in r.text.split("\n")[1:] if line and not line.startswith(">"))
                if seq:
                    out[u] = seq
                    logger.info(f"  {u}: fetched via UniSave (deleted entry)")
        except Exception:
            pass
    still_missing = [u for u in uids if u not in out]
    if still_missing:
        logger.warning(f"{len(still_missing)} sequences not retrieved: {still_missing[:5]}"
                       + ("..." if len(still_missing) > 5 else ""))
    return out


def _parse_fasta_text(text: str) -> dict[str, str]:
    """Parse FASTA text into {header: sequence}."""
    out = {}
    cur, seq = None, []
    for line in text.split("\n"):
        line = line.rstrip("\r")
        if not line:
            continue
        if line.startswith(">"):
            if cur is not None:
                out[cur] = "".join(seq)
            cur = line[1:]
            seq = []
        else:
            seq.append(re.sub(r'[^A-Za-z]', '', line).upper())
    if cur is not None:
        out[cur] = "".join(seq)
    return out


def resolve_organism_to_proteome(organism: str, logger: logging.Logger) -> str | None:
    """Query UniProt proteomes API to resolve organism name to proteome ID.

    Uses the proteomes/search endpoint with organism_name and proteome_type
    filters.  proteome_type values are REFERENCE / NON_REFERENCE / EXCLUDED
    (not numeric).  Falls back to organism_name-only if the REFERENCE filter
    returns nothing.
    """
    url = UNIPROT_PROTEOME_LOOKUP_URL
    # Try with reference proteome filter first
    for query in (
        f'organism_name:"{organism}" AND proteome_type:REFERENCE',
        f'organism_name:"{organism}"',
        organism,  # last resort: free-text search
    ):
        params = {"query": query, "format": "tsv", "size": 10}
        try:
            r = requests.get(url, params=params, timeout=60)
            if r.status_code != 200:
                continue
            lines = r.text.strip().split("\n")
            if len(lines) < 2:
                continue
            header = lines[0].split("\t")
            # Prefer the entry whose Organism field exactly matches (avoids
            # hybrid organisms like "Arabidopsis thaliana x Arabidopsis arenosa")
            best = None
            for line in lines[1:]:
                cols = dict(zip(header, line.split("\t")))
                org_field = cols.get("Organism", "")
                # Exact match on the species name (before any parenthetical)
                org_base = org_field.split("(")[0].strip()
                if org_base.lower() == organism.lower():
                    best = cols
                    break
            if best is None:
                best = dict(zip(header, lines[1].split("\t")))
            upid = best.get("Proteome Id") or best.get("upid")
            if upid:
                logger.info(f"Organism '{organism}' -> proteome {upid} "
                            f"({best.get('Organism','?')}, {best.get('Protein count','?')} proteins)")
                return upid
        except Exception:
            continue
    logger.error(f"Could not resolve organism '{organism}' to a UniProt proteome ID")
    return None


def fetch_proteome_canonical(upid: str, logger: logging.Logger,
                             max_seqs: int | None = None) -> dict[str, str]:
    """Stream the canonical FASTA of a UniProt reference proteome.

    Fetches the full proteome (Swiss-Prot reviewed + TrEMBL unreviewed) so
    that gene-deduplication can collapse isoforms and fragments to one
    representative per gene.  The expensive downstream steps only run on
    regex-positive candidates (~7 %), so downloading the full set adds
    only download + regex-scan time.
    """
    params = {
        "query": f"proteome:{upid}",
        "format": "fasta",
        "compressed": "false",
    }
    logger.info(f"Streaming full proteome {upid} (reviewed + TrEMBL) from UniProt")
    try:
        r = requests.get(UNIPROT_STREAM_URL, params=params, timeout=1200, stream=True)
        r.raise_for_status()
        text_chunks = []
        for chunk in r.iter_content(chunk_size=65536, decode_unicode=True):
            if chunk:
                text_chunks.append(chunk)
        text = "".join(text_chunks)
        seqs = _parse_fasta_text(text)
    except Exception as e:
        logger.error(f"Proteome stream failed: {e}")
        return {}
    # Extract UID from UniProt-style headers
    out = {}
    for hdr, seq in seqs.items():
        parts = hdr.split("|")
        if len(parts) >= 3 and parts[0] in ("sp", "tr"):
            uid = parts[1].upper()
        else:
            uid = hdr.split()[0].upper()
        out[uid] = seq
        if max_seqs and len(out) >= max_seqs:
            break
    logger.info(f"Retrieved {len(out)} sequences from proteome {upid}")
    return out


# ==========================================================================
# REGEX SCAN + SEQUENCE FEATURES  (rules a, c)
# ==========================================================================

@dataclass
class MotifHit:
    uniprot_id: str
    seq_len: int
    motif_start_idx: int    # 1-based inclusive
    motif_end_idx: int      # 1-based inclusive
    motif_seq: str
    motif_len: int
    linker1_len: int        # spacer1: aa between [KR][KR] and [KRH]
    linker2_len: int        # spacer2: aa between [KRH] and P
    # Downstream flank +1..+5 relative to Y (last motif position)
    n_hydrophobic_upstream: int   # -5..-1 window
    n_hydrophobic_downstream: int # +1..+5 window
    downstream_warn: bool
    # Filled by later stages:
    linker1_score: float | None = None
    linker2_score: float | None = None


def scan_motifs(uniprot_id: str, seq: str, strict_regex: bool = False,
                calibration: dict = DEFAULT_CALIBRATION) -> list[MotifHit]:
    """Find all regex-consistent PY-NLS candidate motifs in a sequence."""
    if not seq:
        return []
    seq = re.sub(r'[^A-Z]', 'X', seq.upper())  # non-standard → X (won't match)
    hits = []
    for m in MOTIF_REGEX.finditer(seq):
        motif = m.group(1)
        start = m.start() + 1
        end = start + len(motif) - 1
        # Break motif into components: [KR][KR] + linker1 + [KRH] + linker2 + PY
        # Parse: first 2 chars = [KR][KR], then find last [KRH] before PY at end.
        # Since regex is greedy with overlapping via lookahead, decompose:
        # motif = kk + linker1 + krh + linker2 + "PY"
        # motif ends in PY, then before PY: linker2 (1..5 chars), then [KRH].
        # Search from right: locate PY at end; linker2 is the substring between [KRH] and PY.
        # The [KRH] is the last KRH within the last 6 chars before PY.
        py_idx = len(motif) - 2  # index of 'P'
        # linker2 length varies 1-5: find KRH between positions 2 and py_idx-1
        krh_idx = None
        # We prefer the rightmost KRH such that (py_idx - krh_idx - 1) in [1,5].
        for k in range(py_idx - 2, 1, -1):  # k >= 2 to leave [KR][KR] at 0-1
            if motif[k] in "KRH" and 1 <= (py_idx - k - 1) <= 5:
                krh_idx = k
                break
        if krh_idx is None:
            continue  # should not happen given regex
        linker1_len = krh_idx - 2       # aa between position 1 (2nd of [KR][KR]) and krh
        linker2_len = py_idx - krh_idx - 1
        # Optional strict variant
        if strict_regex and linker1_len > calibration.get("linker1_max_strict", 17):
            continue
        # Flanking hydrophobic counts (positions -5..-1 and +1..+5 relative to Y)
        y_pos = end - 1  # 1-based position of Y in seq
        up_win = seq[max(0, start-1-5):start-1]  # 5 aa upstream of first [KR]
        dn_win = seq[end:end+5]                  # 5 aa downstream of Y
        n_up = sum(1 for a in up_win if a in HYDROPHOBIC)
        n_dn = sum(1 for a in dn_win if a in HYDROPHOBIC)
        warn = n_dn >= calibration.get("hydrophobic_downstream_warn", 3)
        hits.append(MotifHit(
            uniprot_id=uniprot_id, seq_len=len(seq),
            motif_start_idx=start, motif_end_idx=end,
            motif_seq=motif, motif_len=len(motif),
            linker1_len=linker1_len, linker2_len=linker2_len,
            n_hydrophobic_upstream=n_up, n_hydrophobic_downstream=n_dn,
            downstream_warn=warn,
        ))
    return hits


def linker_length_score(linker_len: int, target: float, sigma: float) -> float:
    """Gaussian-centered score, in [0,1]."""
    if linker_len is None or np.isnan(linker_len):
        return 0.0
    return float(np.exp(-0.5 * ((linker_len - target) / sigma) ** 2))


# ==========================================================================
# STRUCTURE  (rules b, c, e — DSSP + FreeSASA + pLDDT)
# ==========================================================================

def find_uniref100_substitute(uid: str, seq_len: int,
                              logger: logging.Logger) -> str | None:
    """Find a 100% sequence-identical UniProt entry that has an AF model.

    Queries the UniRef100 cluster via UniProt REST API to find cluster members
    with the same sequence length, then checks which ones have an AlphaFold
    model available on EBI.
    """
    params = {
        "query": f"uniref_cluster_100:UniRef100_{uid.upper()}",
        "format": "tsv",
        "fields": "accession,length",
        "size": 20,
    }
    try:
        r = requests.get(UNIPROT_ENTRY_URL, params=params, timeout=30)
        r.raise_for_status()
    except Exception:
        return None
    lines = r.text.strip().split("\n")
    if len(lines) < 2:
        return None
    for line in lines[1:]:
        cols = line.split("\t")
        if len(cols) < 2:
            continue
        candidate_uid = cols[0].strip()
        candidate_len_str = cols[1].strip()
        if not candidate_uid or candidate_uid.upper() == uid.upper():
            continue
        try:
            candidate_len = int(candidate_len_str)
        except ValueError:
            candidate_len = 0
        if seq_len > 0 and candidate_len != seq_len:
            continue
        # Check if this candidate has an AF model (HEAD request)
        for v in AF_VERSIONS:
            url = AF_URL_TEMPLATE.format(uid=candidate_uid, v=v)
            try:
                r2 = requests.head(url, timeout=10)
                if r2.status_code == 200:
                    return candidate_uid
            except Exception:
                continue
    return None


def find_uniparc_substitute(uid: str, seq_len: int,
                            logger: logging.Logger) -> str | None:
    """Find an alternative UniProt accession with an AF model via UniParc.

    Queries UniParc for all accessions sharing the same 100% identical
    sequence (same UniParc entry), then checks which alternative
    accessions have an AlphaFold model on EBI.  This catches cases where
    the primary accession was deleted from UniProtKB TrEMBL but
    alternative active accessions for the same sequence still have AF
    models — a scenario not covered by UniRef100 (which may return empty
    for deleted entries).
    """
    params = {
        "query": uid,
        "format": "tsv",
        "fields": "upi,accession,length",
        "size": 5,
    }
    try:
        r = requests.get(UNIPARC_SEARCH_URL, params=params, timeout=30)
        r.raise_for_status()
    except Exception:
        return None
    lines = r.text.strip().split("\n")
    if len(lines) < 2:
        return None
    for line in lines[1:]:
        cols = line.split("\t")
        if len(cols) < 2:
            continue
        # cols: [upi, accession_list, length]
        # accession_list is "; "-separated and may include isoform suffixes
        acc_field = cols[1].strip() if len(cols) >= 2 else ""
        if not acc_field:
            continue
        # Validate length if available (UniParc guarantees same sequence,
        # but this is a safety check)
        if len(cols) >= 3 and seq_len > 0:
            try:
                entry_len = int(cols[2].strip())
                if entry_len != seq_len:
                    continue
            except ValueError:
                pass
        # Parse accessions: split by "; ", strip isoform suffixes
        # Isoform suffixes use either ".N" or "-N".
        candidates = []
        for acc in acc_field.split("; "):
            base = acc.strip()
            # Strip isoform suffixes
            for sep in (".", "-"):
                if sep in base:
                    base = base.split(sep)[0]
            base = base.strip()
            if base and base.upper() != uid.upper() and base not in candidates:
                candidates.append(base)
        # Check each candidate for an AF model (HEAD request)
        for candidate_uid in candidates:
            for v in AF_VERSIONS:
                url = AF_URL_TEMPLATE.format(uid=candidate_uid, v=v)
                try:
                    r2 = requests.head(url, timeout=10)
                    if r2.status_code == 200:
                        return candidate_uid
                except Exception:
                    continue
    return None


def fetch_af_pdb(uid: str, cache_dir: Path, logger: logging.Logger,
                 seq_len: int = 0) -> tuple[Path | None, str]:
    """Fetch monomeric AlphaFold model from EBI DB (v6 → v1).

    Falls back to UniRef100 substitute lookup, then UniParc alternative
    accession lookup, if no model is found for the primary accession.
    Returns (pdb_path, substitute_uid) where substitute_uid is "" when
    the model belongs to *uid* itself, or the substitute accession when
    a UniRef100 or UniParc fallback was used.
    """
    cache_dir.mkdir(parents=True, exist_ok=True)
    # --- Try primary accession ---
    for v in AF_VERSIONS:
        cached = cache_dir / f"{uid}_v{v}.pdb"
        if cached.exists() and cached.stat().st_size > 500:
            return cached, ""
        url = AF_URL_TEMPLATE.format(uid=uid, v=v)
        try:
            r = requests.get(url, timeout=60)
            if r.status_code == 200 and r.content[:6] == b"HEADER":
                cached.write_bytes(r.content)
                return cached, ""
        except Exception:
            continue

    # --- UniRef100 fallback ---
    if seq_len > 0:
        substitute = find_uniref100_substitute(uid, seq_len, logger)
        if substitute:
            logger.info(f"  {uid}: using UniRef100 substitute {substitute}")
            for v in AF_VERSIONS:
                url = AF_URL_TEMPLATE.format(uid=substitute, v=v)
                try:
                    r = requests.get(url, timeout=60)
                    if r.status_code == 200 and r.content[:6] == b"HEADER":
                        # Save under original UID for cache consistency
                        cached = cache_dir / f"{uid}_v{v}.pdb"
                        cached.write_bytes(r.content)
                        return cached, substitute
                except Exception:
                    continue

    # --- UniParc fallback ---
    if seq_len > 0:
        substitute = find_uniparc_substitute(uid, seq_len, logger)
        if substitute:
            logger.info(f"  {uid}: using UniParc substitute {substitute}")
            for v in AF_VERSIONS:
                url = AF_URL_TEMPLATE.format(uid=substitute, v=v)
                try:
                    r = requests.get(url, timeout=60)
                    if r.status_code == 200 and r.content[:6] == b"HEADER":
                        # Save under original UID for cache consistency
                        cached = cache_dir / f"{uid}_v{v}.pdb"
                        cached.write_bytes(r.content)
                        return cached, substitute
                except Exception:
                    continue

    logger.debug(f"  {uid}: no AF model found on EBI DB")
    return None, ""


def parse_pdb_atoms(pdb_path: Path):
    """Return list of dicts, one per ATOM record: resnum, resname, atomname, coord, bfac."""
    atoms = []
    with open(pdb_path) as f:
        for line in f:
            if not line.startswith("ATOM"):
                continue
            try:
                atoms.append({
                    "atomname": line[12:16].strip(),
                    "resname": line[17:20].strip(),
                    "chain": line[21],
                    "resnum": int(line[22:26]),
                    "x": float(line[30:38]),
                    "y": float(line[38:46]),
                    "z": float(line[46:54]),
                    "bfac": float(line[60:66]) if line[60:66].strip() else np.nan,
                })
            except ValueError:
                continue
    return atoms


def run_mkdssp(pdb_path: Path, dssp_bin: str = "mkdssp") -> dict[int, str] | None:
    """Run mkdssp on a PDB file, return {resnum: SSE_char}.

    Sanitizes the PDB by dropping non-ATOM header records (DBREF, SEQRES,
    REMARK) that mkdssp's PDB parser sometimes chokes on when a UniProt
    accession is longer than 6 chars (e.g. A0A6..., which overflows the
    DBREF fixed columns).  Only ATOM/HETATM/TER/END records are needed
    for DSSP calculation.
    """
    if not shutil.which(dssp_bin):
        return None
    try:
        # Sanitize: keep only coordinate records
        with tempfile.NamedTemporaryFile(suffix=".pdb", delete=False, mode="w") as tmp_in:
            in_path = tmp_in.name
            with open(pdb_path) as fin:
                for line in fin:
                    rec = line[:6].strip()
                    if rec in ("ATOM", "HETATM", "TER", "END", "MODEL", "ENDMDL",
                                "CRYST1", "SCALE1", "SCALE2", "SCALE3",
                                "HEADER", "TITLE", "COMPND", "SOURCE",
                                "KEYWDS", "AUTHOR"):
                        tmp_in.write(line)
        with tempfile.NamedTemporaryFile(suffix=".dssp", delete=False) as tmp:
            out_path = tmp.name
        r = subprocess.run([dssp_bin, "--output-format", "dssp", in_path, out_path],
                           capture_output=True, timeout=60)
        try:
            os.unlink(in_path)
        except Exception:
            pass
        if r.returncode != 0:
            return None
        sse = {}
        with open(out_path) as f:
            data = False
            for line in f:
                if line.startswith("  #  RESIDUE"):
                    data = True
                    continue
                if not data:
                    continue
                if len(line) < 17:
                    continue
                try:
                    resnum = int(line[5:10])
                    ss = line[16]
                    if ss == " ":
                        ss = "-"
                    sse[resnum] = ss
                except ValueError:
                    continue
        os.unlink(out_path)
        return sse
    except Exception:
        return None


def dssp_to_hec(sse_dict: dict[int, str]) -> dict[int, str]:
    """Map DSSP 8-state alphabet to H/E/C."""
    hec = {}
    for k, v in sse_dict.items():
        if v in ('H', 'G', 'I'):
            hec[k] = 'H'
        elif v in ('E', 'B'):
            hec[k] = 'E'
        else:
            hec[k] = 'C'
    return hec


def compute_sse_features(sse_hec: dict[int, str], motif_start: int, motif_end: int,
                        seq_len: int, min_helix_run: int = 4,
                        min_beta_run: int = 3, beta_relax_cutoff: float = 0.10) -> dict:
    """
    Rule (b): PY residues must be unstructured; rest of motif preferentially
    unstructured or alpha helix; beta strand strongly unfavored.
    Tolerate SHORT (<3) stretches of E or H because they cannot form real SSE,
    but require they do NOT continue beyond the motif (i.e. the run entirely
    inside the motif is short but if extended past motif boundaries becomes long,
    reject).

    Emits both strict and relaxed pass booleans (side-by-side).
    """
    if not sse_hec:
        return {
            "sse_string": "", "sse_string_valid": "",
            "helix_fraction_raw": np.nan, "beta_fraction_raw": np.nan,
            "helix_run_max_motif": 0, "beta_run_max_motif": 0,
            "core_PY_unstructured": None,
            "passes_sse_strict": None, "passes_sse_relaxed": None,
            "passes_sse_borderline": None,
            "helix_run_extends_upstream": None, "beta_run_extends_downstream": None,
            "sse_source": "NA"
        }
    # Full-length HEC string
    L = seq_len
    full = ''.join(sse_hec.get(i, 'C') for i in range(1, L + 1))
    motif_seg = full[motif_start-1:motif_end]  # 0-based slice
    n_motif = len(motif_seg)
    helix_frac = motif_seg.count('H') / max(1, n_motif)
    beta_frac = motif_seg.count('E') / max(1, n_motif)
    # Max run inside motif
    def max_run(s, ch):
        m = 0; cur = 0
        for c in s:
            if c == ch: cur += 1; m = max(m, cur)
            else: cur = 0
        return m
    helix_run_max = max_run(motif_seg, 'H')
    beta_run_max  = max_run(motif_seg, 'E')

    # Check if short runs at motif boundaries EXTEND into flanking regions.
    # Extension = run continues past motif start or end.
    def run_extends_past_boundary(ch: str) -> bool:
        # Left boundary: does a run at motif_start extend upstream?
        i = motif_start
        left_run = 0
        while i >= 1 and sse_hec.get(i) == ch:
            left_run += 1
            i -= 1
        # If run at boundary is short (<3 for H, <3 for E) inside the motif but
        # extends beyond, that's an extension.
        left_run_inside = 0
        for j in range(motif_start, motif_end + 1):
            if sse_hec.get(j) == ch:
                left_run_inside += 1
            else:
                break
        left_extends = (left_run_inside > 0) and (i >= 1) and (sse_hec.get(motif_start - 1) == ch)

        # Right boundary
        right_run_inside = 0
        for j in range(motif_end, motif_start - 1, -1):
            if sse_hec.get(j) == ch:
                right_run_inside += 1
            else:
                break
        right_extends = (right_run_inside > 0) and (motif_end + 1 <= L) and (sse_hec.get(motif_end + 1) == ch)
        return left_extends or right_extends

    helix_extends = run_extends_past_boundary('H')
    beta_extends  = run_extends_past_boundary('E')

    # Run-length filter: short SSE runs (H<4 or E<3) inside motif -> demote to C
    # ONLY when they do NOT extend beyond the motif (the extension check above).
    valid = list(motif_seg)
    def demote_short(ch: str, min_run: int, extends: bool):
        if extends:
            return  # keep as-is (real SSE that leaks out)
        # Find runs of ch and demote if short
        i = 0
        while i < len(valid):
            if valid[i] == ch:
                j = i
                while j < len(valid) and valid[j] == ch:
                    j += 1
                run_len = j - i
                if run_len < min_run:
                    for k in range(i, j):
                        valid[k] = 'C'
                i = j
            else:
                i += 1
    demote_short('H', min_helix_run, helix_extends)
    demote_short('E', min_beta_run, beta_extends)
    valid_str = ''.join(valid)

    beta_count_valid = valid_str.count('E')
    helix_count_valid = valid_str.count('H')

    # Core PY: last 2 residues of the motif (positions motif_end-1 = P, motif_end = Y)
    py_hec = full[motif_end-2:motif_end]
    core_PY_unstructured = all(c == 'C' for c in py_hec)

    # Strict: no E after run-filter AND core PY is unstructured
    passes_strict = (beta_count_valid == 0) and core_PY_unstructured
    # Relaxed: raw beta fraction < 0.10 AND core PY unstructured (Edison-style)
    passes_relaxed = (beta_frac < beta_relax_cutoff) and core_PY_unstructured
    # Borderline: allow up to 1 residual E and up to 1-length beta run.
    # Alpha-helical structure is NOT penalised — PY-NLS literature explicitly
    # tolerates helical scaffolds (the motif prefers unstructured OR alpha-helical,
    # beta-strand is the only forbidden SSE).  The previous helix_run_max <= 6
    # constraint was removed in v6 because it vetoed purely helical motifs with
    # zero beta strand (e.g. At1g04770, helix_run=9, no beta → was tier 4, should
    # be borderline/acceptable).
    passes_borderline = (
        (beta_count_valid <= 1)
        and (beta_run_max <= 1)
    )

    return {
        "sse_string": motif_seg,
        "sse_string_valid": valid_str,
        "helix_fraction_raw": helix_frac,
        "beta_fraction_raw": beta_frac,
        "helix_run_max_motif": helix_run_max,
        "beta_run_max_motif": beta_run_max,
        "core_PY_unstructured": core_PY_unstructured,
        "passes_sse_strict": passes_strict,
        "passes_sse_relaxed": passes_relaxed,
        "passes_sse_borderline": passes_borderline,
        "helix_run_extends_upstream": helix_extends,
        "beta_run_extends_downstream": beta_extends,
        "sse_source": "mkdssp"
    }


def compute_plddt_features(atoms: list, motif_start: int, motif_end: int) -> dict:
    """Mean pLDDT (from B-factor column) over motif CA atoms."""
    ca = [a for a in atoms if a["atomname"] == "CA" and motif_start <= a["resnum"] <= motif_end]
    if not ca:
        return {"mean_pLDDT_motif": np.nan, "min_pLDDT_motif": np.nan,
                "pLDDT_source": "NA"}
    vals = [a["bfac"] for a in ca if not np.isnan(a["bfac"])]
    return {
        "mean_pLDDT_motif": float(np.mean(vals)) if vals else np.nan,
        "min_pLDDT_motif": float(np.min(vals)) if vals else np.nan,
        "pLDDT_source": "AF"
    }


def compute_sasa_features(pdb_path: Path, motif_start: int, motif_end: int) -> dict:
    """FreeSASA-based RSA in percent using Tien 2013 max-ASA."""
    try:
        import freesasa
        # Silence freesasa verbosity for cleaner logs
        try:
            freesasa.setVerbosity(freesasa.silent)
        except Exception:
            pass
        struct = freesasa.Structure(str(pdb_path))
        # API changed across versions: prefer freesasa.calc, fall back to Calc()
        if hasattr(freesasa, "calc"):
            result = freesasa.calc(struct)
        else:
            result = freesasa.Calc().calculate(struct)
        # residueAreas: dict chain -> resnum -> ResidueArea
        areas = result.residueAreas()
    except Exception as e:
        return {"mean_RSA_motif_pct": np.nan, "min_RSA_motif_pct": np.nan,
                "mean_RSA_PY_core_pct": np.nan, "sasa_source": f"NA:{type(e).__name__}"}
    # Assume chain 'A'; some AF files use other single-chain IDs.
    chain_key = next(iter(areas.keys()))
    per_res = areas[chain_key]

    def rsa(resnum: int) -> float:
        ra = per_res.get(str(resnum))
        if ra is None:
            return np.nan
        total = ra.total
        aa = ra.residueType.upper()
        if aa in AA3_TO_1:
            aa1 = AA3_TO_1[aa]
            max_asa = TIEN_MAX_ASA.get(aa1, np.nan)
        else:
            return np.nan
        if not max_asa or np.isnan(max_asa):
            return np.nan
        return (total / max_asa) * 100.0

    rsa_vals = []
    for r in range(motif_start, motif_end + 1):
        v = rsa(r)
        if not np.isnan(v):
            rsa_vals.append(v)
    if not rsa_vals:
        return {"mean_RSA_motif_pct": np.nan, "min_RSA_motif_pct": np.nan,
                "mean_RSA_PY_core_pct": np.nan, "sasa_source": "freesasa"}
    # PY core = last 2 residues; extended PY-adjacent = last 5 aa
    py_range = range(motif_end - 4, motif_end + 1)
    py_vals = [rsa(r) for r in py_range if not np.isnan(rsa(r))]
    return {
        "mean_RSA_motif_pct": float(np.mean(rsa_vals)),
        "min_RSA_motif_pct":  float(np.min(rsa_vals)),
        "mean_RSA_PY_core_pct": float(np.mean(py_vals)) if py_vals else np.nan,
        "sasa_source": "freesasa"
    }


# ==========================================================================
# DISORDER  (rule c — metapredict + IUPred3)
# ==========================================================================

def compute_metapredict(seq: str, motif_start: int, motif_end: int) -> dict:
    """metapredict disorder score across motif + flanks + full protein."""
    try:
        import metapredict as meta
        disorder = np.array(meta.predict_disorder(seq))
    except Exception:
        return {"meta_disorder_motif_mean": np.nan, "meta_disorder_motif_min": np.nan,
                "meta_disorder_motif_max": np.nan,
                "meta_disorder_frac_ge_0p5_motif": np.nan,
                "meta_disorder_full_mean": np.nan, "meta_source": "NA"}
    motif_slice = disorder[motif_start-1:motif_end]
    return {
        "meta_disorder_motif_mean": float(motif_slice.mean()) if len(motif_slice) else np.nan,
        "meta_disorder_motif_min": float(motif_slice.min()) if len(motif_slice) else np.nan,
        "meta_disorder_motif_max": float(motif_slice.max()) if len(motif_slice) else np.nan,
        "meta_disorder_frac_ge_0p5_motif": float((motif_slice >= 0.5).mean()) if len(motif_slice) else np.nan,
        "meta_disorder_full_mean": float(disorder.mean()) if len(disorder) else np.nan,
        "meta_source": "metapredict"
    }


class IUPred3Client:
    """IUPred3 client using the official REST API:
        GET https://iupred3.elte.hu/iupred3/{accession}[.json]
        GET https://iupred3.elte.hu/iupred3/{type}/{accession}[.json]

    Response fields (json): sequence, iupred2 (list), type, meta.
    Falls back to Bokeh scraping only for raw-sequence input (which the
    REST API does not support).

    Returns np.ndarray of per-residue IUPred3 scores, or None on failure.
    """

    IUPRED3_REST_TEMPLATE = "https://iupred3.elte.hu/iupred3/{itype}/{acc}.json"

    def __init__(self):
        self.sess = requests.Session()

    def predict_by_accession(self, uid: str, itype: str = "long",
                              retries: int = 3, backoff: float = 1.5) -> np.ndarray | None:
        url = self.IUPRED3_REST_TEMPLATE.format(itype=itype, acc=uid)
        for attempt in range(retries):
            try:
                r = self.sess.get(url, timeout=60)
                if r.status_code == 200:
                    payload = r.json()
                    arr = payload.get("iupred2")
                    if arr is not None and isinstance(arr, list):
                        return np.array(arr, dtype=float)
                elif r.status_code in (404, 500):
                    return None  # deleted / obsolete UniProt entry
            except Exception:
                pass
            if attempt < retries - 1:
                time.sleep(backoff ** (attempt + 1))
        return None

    def _get_csrf(self) -> str | None:
        try:
            r = self.sess.get(f"{IUPRED3_BASE}/", timeout=30)
            r.raise_for_status()
            m = re.search(r'name="csrfmiddlewaretoken"\s+value="([^"]+)"', r.text)
            return m.group(1) if m else None
        except Exception:
            return None

    def predict_by_sequence(self, seq: str, retries: int = 3, backoff: float = 1.5) -> np.ndarray | None:
        """Fallback for raw sequence input: submit via web form, then hit the
        session-scoped download endpoint. Less reliable than the REST API."""
        for attempt in range(retries):
            csrf = self._get_csrf()
            if not csrf:
                if attempt < retries - 1:
                    time.sleep(backoff ** (attempt + 1))
                    continue
                return None
            try:
                form = {'csrfmiddlewaretoken': csrf, 'inp_seq': seq,
                         'iupred_type': 'long', 'context': 'anchor', 'smoothing': 'medium'}
                r2 = self.sess.post(f"{IUPRED3_BASE}/plot", data=form,
                                     headers={'Referer': f"{IUPRED3_BASE}/"},
                                     timeout=60)
                r2.raise_for_status()
                # Try the raw-text download endpoint (result_download_txt)
                for url in (f"{IUPRED3_BASE}/result_download_txt", f"{IUPRED3_BASE}/download"):
                    try:
                        r3 = self.sess.get(url, timeout=60,
                                            headers={'Referer': f"{IUPRED3_BASE}/plot"})
                        if r3.status_code == 200 and r3.text.strip():
                            arr = _parse_iupred_text(r3.text)
                            if arr is not None:
                                return arr
                    except Exception:
                        continue
            except Exception:
                pass
            if attempt < retries - 1:
                time.sleep(backoff ** (attempt + 1))
        return None

    def predict(self, seq: str, uid: str | None = None, analysis: str = "long") -> np.ndarray | None:
        """Try REST-by-accession first; fall back to raw-sequence Web submission."""
        arr = None
        if uid:
            arr = self.predict_by_accession(uid, itype=analysis)
            if arr is not None and len(arr) == len(seq):
                return arr
            if arr is not None and len(arr) != len(seq):
                # Deleted/altered UniProt entry — we have a sequence mismatch.
                arr = None
        arr2 = self.predict_by_sequence(seq)
        if arr2 is not None and len(arr2) == len(seq):
            return arr2
        return arr if arr is not None else arr2


def _parse_iupred_text(text: str) -> np.ndarray | None:
    """Parse IUPred3 text-download format (# comments then 'pos aa score [anchor]')."""
    scores = []
    for line in text.split("\n"):
        line = line.strip()
        if not line or line.startswith("#"):
            continue
        parts = line.split()
        if len(parts) >= 3:
            try:
                scores.append(float(parts[2]))
            except ValueError:
                continue
    return np.array(scores, dtype=float) if scores else None


def compute_iupred3(seq: str, motif_start: int, motif_end: int,
                    client: IUPred3Client) -> dict:
    arr = client.predict(seq, analysis="long")
    if arr is None or len(arr) != len(seq):
        return {"iupred3_motif_mean": np.nan, "iupred3_motif_min": np.nan,
                "iupred3_motif_max": np.nan, "iupred3_frac_ge_0p5_motif": np.nan,
                "iupred3_source": "unavailable"}
    m = arr[motif_start-1:motif_end]
    return {
        "iupred3_motif_mean": float(m.mean()) if len(m) else np.nan,
        "iupred3_motif_min": float(m.min()) if len(m) else np.nan,
        "iupred3_motif_max": float(m.max()) if len(m) else np.nan,
        "iupred3_frac_ge_0p5_motif": float((m >= 0.5).mean()) if len(m) else np.nan,
        "iupred3_source": "iupred3_rest"
    }


# ==========================================================================
# AIUPred  (local CLI; default disorder predictor, replaces IUPred3 REST)
#   Erdos & Dosztanyi (2024) Nucleic Acids Res. 52(W1):W176-W181
#   Erdos, Deutsch & Dosztanyi (2025) J. Mol. Biol. 437:169071
# ==========================================================================

AIUPRED_BIN = shutil.which("aiupred") or "aiupred"


class AIUPredClient:
    """Local AIUPred CLI wrapper.

    Runs:  aiupred -i <fasta> -o <tsv> -b -l [--force-cpu]
    Parses the TSV to extract per-residue Disorder, Binding, and Linker
    probability arrays.  Returns a dict of np.ndarrays keyed by
    'disorder', 'binding', 'linker', or None on failure.

    AIUPred is the recommended default disorder predictor (replaces the
    IUPred3 REST endpoint).  It is run locally so there is no rate limit,
    and it additionally provides binding-region and flexible-linker
    predictions that IUPred3 does not.
    """

    def __init__(self, binary: str = AIUPRED_BIN, force_cpu: bool = True,
                 timeout: int = 600):
        self.binary = binary
        self.force_cpu = force_cpu
        self.timeout = timeout
        # Cache results by sequence hash to avoid re-running on identical seqs
        self._cache: dict[str, dict] = {}

    def predict(self, seqs: dict[str, str]) -> dict[str, dict]:
        """Run AIUPred on a batch of {uid: sequence} entries.

        Writes a single multi-FASTA, invokes aiupred once, and parses the
        multi-protein TSV output.  Returns {uid: {'disorder': arr,
        'binding': arr, 'linker': arr}} (entries that fail get empty dict).
        """
        # Filter to sequences we haven't cached and that are non-empty
        to_run = {u: s for u, s in seqs.items()
                  if s and u not in self._cache}
        if not to_run:
            return {u: self._cache.get(u, {}) for u in seqs}

        out: dict[str, dict] = {}
        # Write multi-FASTA to a temp file
        with tempfile.NamedTemporaryFile(suffix=".fasta", delete=False, mode="w") as fin:
            in_path = fin.name
            for uid, s in to_run.items():
                # Sanitise header: aiupred preserves it in the #> line
                fin.write(f">{uid}\n{s}\n")
        with tempfile.NamedTemporaryFile(suffix=".tsv", delete=False) as fout:
            out_path = fout.name
        try:
            cmd = [self.binary, "-i", in_path, "-o", out_path, "-b", "-l"]
            if self.force_cpu:
                cmd.append("--force-cpu")
            # Scale timeout with batch size: ~2s per 1000 residues on CPU
            total_residues = sum(len(s) for s in to_run.values())
            dynamic_timeout = max(self.timeout, int(total_residues / 1000 * 3) + 60)
            r = subprocess.run(cmd, capture_output=True, timeout=dynamic_timeout)
            if r.returncode != 0:
                # Failed — return empty dicts for all
                for u in to_run:
                    out[u] = {}
                return {u: out.get(u, self._cache.get(u, {})) for u in seqs}
            # Parse the TSV
            parsed = self._parse_tsv(out_path)
            for u in to_run:
                self._cache[u] = parsed.get(u, {})
                out[u] = parsed.get(u, {})
        except Exception:
            for u in to_run:
                out[u] = {}
        finally:
            for p in (in_path, out_path):
                try:
                    os.unlink(p)
                except Exception:
                    pass
        return {u: out.get(u, self._cache.get(u, {})) for u in seqs}

    @staticmethod
    def _parse_tsv(path: str) -> dict[str, dict]:
        """Parse aiupred TSV output (multi-protein).

        Format:
          # ... comment lines ...
          # Position\tResidue\tDisorder\tBinding\tLinker
          #>protein_header
          1\tM\t0.969\t0.982\t0.002
          ...
        Returns {uid: {'disorder': arr, 'binding': arr, 'linker': arr}}.
        """
        out: dict[str, dict] = {}
        cur_uid: str | None = None
        cur_dis: list[float] = []
        cur_bin: list[float] = []
        cur_lnk: list[float] = []

        def _flush():
            nonlocal cur_uid, cur_dis, cur_bin, cur_lnk
            if cur_uid is not None and cur_dis:
                out[cur_uid] = {
                    "disorder": np.array(cur_dis, dtype=float),
                    "binding":  np.array(cur_bin, dtype=float),
                    "linker":   np.array(cur_lnk, dtype=float),
                }
            cur_uid = None; cur_dis = []; cur_bin = []; cur_lnk = []

        with open(path) as f:
            for line in f:
                line = line.rstrip("\n")
                if not line:
                    continue
                if line.startswith("#>"):
                    _flush()
                    # Header is "#>header" — take first whitespace-delimited
                    # token after '>' as the UID
                    hdr = line[2:].strip()
                    cur_uid = hdr.split()[0].lstrip(">") if hdr else None
                    # Strip a leading '>' if present (some versions)
                    if cur_uid and cur_uid.startswith(">"):
                        cur_uid = cur_uid[1:]
                    # Strip UniProt-style prefixes: tr|UID|... or sp|UID|...
                    if cur_uid in ("tr", "sp") and len(hdr.split()) > 1:
                        parts = hdr.split()
                        if len(parts) >= 2 and "|" in parts[1]:
                            cur_uid = parts[1].split("|")[1]
                    elif "|" in cur_uid:
                        parts = cur_uid.split("|")
                        if len(parts) >= 2 and parts[0] in ("tr", "sp"):
                            cur_uid = parts[1]
                elif line.startswith("#"):
                    continue
                else:
                    parts = line.split("\t")
                    if len(parts) >= 5 and cur_uid is not None:
                        try:
                            cur_dis.append(float(parts[2]))
                            cur_bin.append(float(parts[3]))
                            cur_lnk.append(float(parts[4]))
                        except ValueError:
                            continue
        _flush()
        return out


def compute_aiupred(aiupred_result: dict, motif_start: int, motif_end: int) -> dict:
    """Slice AIUPred arrays to the motif window and compute summary stats.

    Returns dict with keys:
      aiupred_disorder_motif_mean / _min / _max / _frac_ge_0p5
      aiupred_binding_motif_mean / _min / _max / _frac_ge_0p5
      aiupred_linker_motif_mean / _min / _max
      aiupred_source
    """
    if not aiupred_result:
        return {
            "aiupred_disorder_motif_mean": np.nan, "aiupred_disorder_motif_min": np.nan,
            "aiupred_disorder_motif_max": np.nan, "aiupred_disorder_frac_ge_0p5_motif": np.nan,
            "aiupred_binding_motif_mean": np.nan, "aiupred_binding_motif_min": np.nan,
            "aiupred_binding_motif_max": np.nan, "aiupred_binding_frac_ge_0p5_motif": np.nan,
            "aiupred_linker_motif_mean": np.nan, "aiupred_linker_motif_min": np.nan,
            "aiupred_linker_motif_max": np.nan,
            "aiupred_source": "unavailable",
        }
    out = {"aiupred_source": "aiupred_local"}
    for key, prefix in (("disorder", "aiupred_disorder"), ("binding", "aiupred_binding")):
        arr = aiupred_result.get(key)
        if arr is None or len(arr) == 0:
            out[f"{prefix}_motif_mean"] = np.nan
            out[f"{prefix}_motif_min"] = np.nan
            out[f"{prefix}_motif_max"] = np.nan
            out[f"{prefix}_frac_ge_0p5_motif"] = np.nan
        else:
            m = arr[motif_start - 1:motif_end]
            if len(m):
                out[f"{prefix}_motif_mean"] = float(m.mean())
                out[f"{prefix}_motif_min"] = float(m.min())
                out[f"{prefix}_motif_max"] = float(m.max())
                out[f"{prefix}_frac_ge_0p5_motif"] = float((m >= 0.5).mean())
            else:
                out[f"{prefix}_motif_mean"] = np.nan
                out[f"{prefix}_motif_min"] = np.nan
                out[f"{prefix}_motif_max"] = np.nan
                out[f"{prefix}_frac_ge_0p5_motif"] = np.nan
    # Linker (no frac_ge_0p5 — linker is a different kind of signal)
    arr = aiupred_result.get("linker")
    if arr is None or len(arr) == 0:
        out["aiupred_linker_motif_mean"] = np.nan
        out["aiupred_linker_motif_min"] = np.nan
        out["aiupred_linker_motif_max"] = np.nan
    else:
        m = arr[motif_start - 1:motif_end]
        if len(m):
            out["aiupred_linker_motif_mean"] = float(m.mean())
            out["aiupred_linker_motif_min"] = float(m.min())
            out["aiupred_linker_motif_max"] = float(m.max())
        else:
            out["aiupred_linker_motif_mean"] = np.nan
            out["aiupred_linker_motif_min"] = np.nan
            out["aiupred_linker_motif_max"] = np.nan
    return out


# ==========================================================================
# UniProt annotation  (rule f)
# ==========================================================================

def fetch_uniprot_annotations(uids: list[str], logger: logging.Logger,
                              batch_size: int = 100) -> dict[str, dict]:
    """Batch-fetch TM/signal/transit/propeptide + gene names from UniProt REST.
    """
    out = {u.upper(): {"TM_helix": "", "Signal_peptide": "", "Transit_peptide": "",
                       "Propeptide": "", "gene_primary": "", "gene_names": "",
                       "gene_oln": "", "gene_orf": "", "protein_name": ""} for u in uids}
    fields = ("accession,ft_transmem,ft_signal,ft_propep,ft_transit,"
              "gene_primary,gene_names,gene_oln,gene_orf,protein_name")
    logger.info(f"Fetching UniProt annotations for {len(uids)} IDs")
    for i in range(0, len(uids), batch_size):
        batch = [u.upper() for u in uids[i:i+batch_size]]
        query = " OR ".join(f"accession:{u}" for u in batch)
        params = {"query": query, "format": "tsv", "fields": fields, "size": 500}
        try:
            r = requests.get(UNIPROT_ENTRY_URL, params=params, timeout=120)
            r.raise_for_status()
        except Exception as e:
            logger.warning(f"UniProt annot batch {i//batch_size+1} failed: {e}")
            continue
        lines = r.text.strip().split("\n")
        if len(lines) < 2:
            continue
        header = lines[0].split("\t")
        for line in lines[1:]:
            cols = line.split("\t")
            if len(cols) < len(header):
                cols += [""] * (len(header) - len(cols))
            row = dict(zip(header, cols))
            uid = row.get("Entry", "").upper()
            if uid in out:
                out[uid] = {
                    "TM_helix": row.get("Transmembrane", ""),
                    "Signal_peptide": row.get("Signal peptide", ""),
                    "Transit_peptide": row.get("Transit peptide", ""),
                    "Propeptide": row.get("Propeptide", ""),
                    "gene_primary": row.get("Gene Names (primary)", ""),
                    "gene_names": row.get("Gene Names", ""),
                    "gene_oln": row.get("Gene Names (ordered locus)", ""),
                    "gene_orf": row.get("Gene Names (ORF)", ""),
                    "protein_name": row.get("Protein names", ""),
                }
        time.sleep(0.1)
    return out


def has_annotation_value(x) -> bool:
    if x is None or (isinstance(x, float) and np.isnan(x)):
        return False
    s = str(x).strip()
    return s != '' and s.lower() not in {'nan', 'none', 'no', 'false', '0'}


def resolve_gene_key(annot: dict) -> str:
    """Cross-organism gene key extraction from UniProt annotation.

    Priority (v6 — ordered locus names first for better isoform merging):
      1. gene_oln  — ordered locus name (e.g. At1g19180 for Arabidopsis).
                     Most stable across isoforms; shared by all splice variants
                     of a gene.  This is the AGI locus code in Arabidopsis.
      2. gene_primary — UniProt curated primary gene symbol (e.g. TIFY10A, TP53).
      3. gene_names  — first token of the full gene names string (fallback).
      4. gene_orf    — ORF name (less stable but better than accession).
      5. Empty string (no gene annotation available).

    For Arabidopsis, using gene_oln (AGI codes) as the primary key ensures
    that all isoforms of a gene are merged into a single entry.  For organisms
    without ordered locus names (e.g. human), gene_primary is used instead.
    """
    oln = annot.get("gene_oln", "")
    if oln and oln.strip():
        return oln.strip()
    gp = annot.get("gene_primary", "")
    if gp and gp.strip():
        return gp.strip()
    gn = annot.get("gene_names", "")
    if gn and gn.strip():
        return gn.strip().split()[0]
    orf = annot.get("gene_orf", "")
    if orf and orf.strip():
        return orf.strip()
    return ""


def parse_tm_spans(s) -> list[tuple[int, int]]:
    if not isinstance(s, str) or 'TRANSMEM' not in s:
        return []
    return [(int(a), int(b)) for a, b in _TRANSMEM_RE.findall(s)]


# ==========================================================================
# cNLS-Mapper (rule f complement)
# ==========================================================================

def query_cnls_mapper(seq: str, cutoff: float = 5.0, timeout: int = 60,
                      linker_mode: str = "entire") -> dict:
    """Submit sequence to cNLS Mapper; return best mp + bp scores.

    Parameters
    ----------
    linker_mode : str
        Controls the cNLS Mapper form field ``linker``:
        - ``"entire"`` (default): search the *Entire region* for bipartite
          NLSs with long linkers (13-20 aa).  This is the v6 default —
          the previous ``long_bp=off`` was the wrong field name and was
          silently ignored, defaulting to terminal-60-aa-only search.
        - ``"terminal"``: search only within terminal 60-aa regions.
    """
    clean = re.sub(r'[^A-Za-z]', '', seq).upper()
    if len(clean) < 11 or len(clean) > 4999:
        return {"cNLS_present": False, "cNLS_mp_max_score": np.nan,
                "cNLS_bp_max_score": np.nan, "cNLS_source": "seq_length"}
    invalid = set(clean) - set("ACDEFGHIKLMNPQRSTVWY")
    if invalid:
        return {"cNLS_present": False, "cNLS_mp_max_score": np.nan,
                "cNLS_bp_max_score": np.nan, "cNLS_source": "invalid_aa"}
    linker_value = ("Entire region" if linker_mode == "entire"
                    else "Within terminal 60-amino-acid regions")
    data = {"typedseq": clean, "cut_off": str(cutoff), "linker": linker_value}
    try:
        r = requests.post(CNLS_MAPPER_URL, data=data, timeout=timeout)
        r.raise_for_status()
    except Exception:
        return {"cNLS_present": False, "cNLS_mp_max_score": np.nan,
                "cNLS_bp_max_score": np.nan, "cNLS_source": "unavailable"}
    html = r.text
    mp_scores = _extract_scores(html, "Predicted monopartite NLS")
    bp_scores = _extract_scores(html, "Predicted bipartite NLS")
    return {
        "cNLS_present": bool(mp_scores or bp_scores),
        "cNLS_mp_max_score": float(max(mp_scores)) if mp_scores else np.nan,
        "cNLS_bp_max_score": float(max(bp_scores)) if bp_scores else np.nan,
        "cNLS_source": "cnls_mapper"
    }


def _extract_scores(html: str, label: str) -> list[float]:
    """Extract cNLS Mapper scores from the HTML section under `label`.

    Scores can appear as either floats (e.g. 12.5) or integers (e.g. 4). The
    section layout has alternating <code> blocks with position (int), sequence,
    then score (int or float). We take EVERY <code> value in the section, then
    keep the numeric ones that could be scores by their position relative to
    <td align="center"> markers.
    """
    parts = html.split(label)
    if len(parts) < 2:
        return []
    section = parts[1].split("</TABLE>")[0] if "</TABLE>" in parts[1] else parts[1][:5000]
    # The score cells are marked as <td align="center">. Extract those cells
    # and their <code>...</code> contents.
    score_cells = re.findall(r'<td align="center">(.*?)</td>', section, re.DOTALL)
    scores = []
    for cell in score_cells:
        for val in re.findall(r'<code>([-+]?[0-9]+(?:\.[0-9]+)?)</code>', cell):
            try:
                scores.append(float(val))
            except ValueError:
                continue
    return scores


# ==========================================================================
# Topology gate (rule g)
# ==========================================================================

def compute_topology_penalty(motif_start: int, motif_end: int,
                             tm_spans: list[tuple[int, int]],
                             linker_threshold: int = 50,
                             polytopic_min_loop: int = 50,
                             soft_multiplier: float = 0.5) -> dict:
    """Plant default: linker >=50 aa from TM. Motif in TM = veto."""
    if not tm_spans:
        return dict(status='no_tm', multiplier=1.0, n_tm=0,
                    linker_dist=None, motif_in_tm=False, in_short_loop=False)
    motif_in_tm = any(not (motif_end < ts or motif_start > te) for ts, te in tm_spans)
    if motif_in_tm:
        return dict(status='hard_veto_motif_in_tm', multiplier=0.0,
                    n_tm=len(tm_spans), linker_dist=0,
                    motif_in_tm=True, in_short_loop=False)
    min_dist = min(min(abs(motif_start - te), abs(motif_end - ts)) for ts, te in tm_spans)
    in_short_loop = False
    for i in range(len(tm_spans) - 1):
        ts_i, te_i = tm_spans[i]; ts_j, te_j = tm_spans[i+1]
        if te_i < motif_start and motif_end < ts_j:
            if (ts_j - te_i - 1) < polytopic_min_loop:
                in_short_loop = True; break
    if in_short_loop:
        return dict(status='hard_veto_loop', multiplier=0.0,
                    n_tm=len(tm_spans), linker_dist=min_dist,
                    motif_in_tm=False, in_short_loop=True)
    if min_dist < linker_threshold:
        return dict(status='hard_short_linker', multiplier=0.0,
                    n_tm=len(tm_spans), linker_dist=min_dist,
                    motif_in_tm=False, in_short_loop=False)
    return dict(status='compliant', multiplier=float(soft_multiplier),
                n_tm=len(tm_spans), linker_dist=min_dist,
                motif_in_tm=False, in_short_loop=False)


# ==========================================================================
# AF3 opportunistic evidence
# ==========================================================================

def find_af3_files(af3_dir: Path, uid: str) -> list[tuple[Path, Path]] | None:
    """Find matching AF3 files for a UniProt ID in the given directory.
    Expected schema: fold_*_<uid>_*model_{0..4}*.txt and *_byres.txt.
    Case-insensitive UID match.
    """
    if not af3_dir or not af3_dir.exists():
        return None
    uid_lower = uid.lower()
    # Look for candidate folder names OR flat files
    matches = []
    for p in af3_dir.rglob(f"*{uid_lower}*model_*.txt"):
        if "_byres.txt" in p.name:
            continue
        byres = p.parent / p.name.replace(".txt", "_byres.txt")
        if byres.exists():
            matches.append((p, byres))
    if not matches:
        # Case-sensitive alt
        for p in af3_dir.rglob(f"*{uid}*model_*.txt"):
            if "_byres.txt" in p.name:
                continue
            byres = p.parent / p.name.replace(".txt", "_byres.txt")
            if byres.exists():
                matches.append((p, byres))
    return matches if matches else None


def parse_af3_metrics(uid: str, files: list[tuple[Path, Path]]) -> dict:
    """Extract minimal AF3 evidence per protein: mean/max of ipSAE-d0chn,
    ipSAE-d0dom, and ipTM across the 5 models, plus whole-chain iptm/ptm."""
    ipsae_d0chn = []; ipsae_d0dom = []; iptm_pae = []
    wc_iptm = []; wc_ptm = []
    for wc_path, byres_path in files:
        # Parse whole-chain file (blank line 1, header line 2)
        try:
            wc = pd.read_csv(wc_path, sep=r'\s+', skiprows=1)
        except Exception:
            continue
        # Prefer the asym B->A row (motif is on chain A of cargo)
        if 'Type' in wc.columns:
            asym = wc[wc['Type'] == 'asym']
            # Pick the row where Chn2 == 'A' (scored chain A)
            row = None
            for _, r in asym.iterrows():
                if r.get('Chn2') == 'A':
                    row = r; break
            if row is None and len(asym) > 0:
                row = asym.iloc[0]
        else:
            row = wc.iloc[0] if len(wc) else None
        if row is None:
            continue
        # These are whole-model ipTM/PTM from the file (columns vary)
        for col in ('ipTM_af', 'ipTM'):
            if col in row.index and pd.notna(row[col]):
                wc_iptm.append(float(row[col])); break
        for col in ('pTM_af', 'pTM', 'PTM'):
            if col in row.index and pd.notna(row[col]):
                wc_ptm.append(float(row[col])); break

        # Parse byres and compute mean of ipSAE_d0chn per protein
        try:
            byres = pd.read_csv(byres_path, sep=r'\s+')
        except Exception:
            continue
        chn_a = byres[byres['ScoredChain'] == 'A'] if 'ScoredChain' in byres.columns else byres
        for col_dst, col_src in [('ipsae_d0chn', 'ipSAE_d0chn'),
                                  ('ipsae_d0dom', 'ipSAE_d0dom'),
                                  ('iptm_pae', 'ipTM_pae')]:
            if col_src in chn_a.columns:
                vals = pd.to_numeric(chn_a[col_src], errors='coerce').dropna()
                if len(vals):
                    if col_dst == 'ipsae_d0chn':
                        ipsae_d0chn.append(float(vals.mean()))
                    elif col_dst == 'ipsae_d0dom':
                        ipsae_d0dom.append(float(vals.mean()))
                    else:
                        iptm_pae.append(float(vals.mean()))
    def _stat(xs):
        return (float(np.mean(xs)) if xs else np.nan, float(np.max(xs)) if xs else np.nan)

    ipsae_d0chn_mean, ipsae_d0chn_max = _stat(ipsae_d0chn)
    ipsae_d0dom_mean, ipsae_d0dom_max = _stat(ipsae_d0dom)
    iptm_pae_mean, iptm_pae_max = _stat(iptm_pae)
    return {
        "af3_ipSAE_d0chn_mean_across_models": ipsae_d0chn_mean,
        "af3_ipSAE_d0chn_max_across_models": ipsae_d0chn_max,
        "af3_ipSAE_d0dom_mean_across_models": ipsae_d0dom_mean,
        "af3_ipSAE_d0dom_max_across_models": ipsae_d0dom_max,
        "af3_ipTM_pae_mean_across_models": iptm_pae_mean,
        "af3_ipTM_pae_max_across_models": iptm_pae_max,
        "af3_wc_iptm_mean": float(np.mean(wc_iptm)) if wc_iptm else np.nan,
        "af3_wc_ptm_mean": float(np.mean(wc_ptm)) if wc_ptm else np.nan,
        "af3_source": "af3_dir",
    }


def load_af3_from_table(af3_table_path: Path) -> dict[str, dict]:
    """Parse the pre-aggregated 214-protein table format into per-UID summaries.
    Used when the AF3 evidence has already been aggregated by an upstream script
    (analyze_segments_byres_exact_v2.py schema).
    """
    df = pd.read_csv(af3_table_path, sep="\t")
    df["uniprot_id_uc"] = df["uniprot_id"].str.upper()
    out = {}
    # For each UID, aggregate the 5 models
    for uid, g in df.groupby("uniprot_id_uc"):
        rec = {
            "af3_ipSAE_d0chn_mean_across_models": float(g["ipSAE_d0chn__mean"].mean()),
            "af3_ipSAE_d0chn_max_across_models": float(g["ipSAE_d0chn__mean"].max()),
            "af3_ipSAE_d0dom_mean_across_models": float(g["ipSAE_d0dom__mean"].mean()),
            "af3_ipSAE_d0dom_max_across_models": float(g["ipSAE_d0dom__mean"].max()),
            "af3_ipTM_pae_mean_across_models": float(g["ipTM_pae__mean"].mean()),
            "af3_ipTM_pae_max_across_models": float(g["ipTM_pae__mean"].max()),
            "af3_wc_iptm_mean": float(g["FL_wc_iptm"].mean()) if "FL_wc_iptm" in g.columns else np.nan,
            "af3_wc_ptm_mean": float(g["FL_wc_ptm"].mean()) if "FL_wc_ptm" in g.columns else np.nan,
            "af3_PY_wc_iptm_mean": float(g["PY_wc_iptm"].mean()) if "PY_wc_iptm" in g.columns else np.nan,
            "af3_PY_wc_ptm_mean": float(g["PY_wc_ptm"].mean()) if "PY_wc_ptm" in g.columns else np.nan,
            "af3_source": "af3_table",
        }
        out[uid] = rec
    return out


# ==========================================================================
# COMPONENT SCORES + FUSED SCORE + TIER
# ==========================================================================

def clip01(x: float) -> float:
    if x is None or np.isnan(x):
        return np.nan
    return float(min(1.0, max(0.0, x)))


def compute_component_scores(row: dict, cal: dict) -> dict:
    """Compute the 7 orthogonal component scores in [0,1] from a feature row."""
    # S_sequence: linker geometry (pure, no arbitrary baseline) + downstream
    # hydrophobic penalty.  
    # Linker1 is weighted 0.6 (vs 0.5
    # previously) because it is the experimentally validated determinant of
    # NLS activity (linker1 length study: L1=17 → 4–7x activity, L1=5 → 1.9x).
    l1_score = linker_length_score(row.get("linker1_len"), cal["linker1_target"], cal["linker1_sigma"])
    l2_score = linker_length_score(row.get("linker2_len"), cal["linker2_target"], cal["linker2_sigma"])
    down_penalty = 1.0 if not row.get("downstream_warn") else 0.5
    S_sequence = clip01(0.6 * l1_score + 0.4 * l2_score) * down_penalty

    # S_disorder: consensus disorder from AIUPred (primary) + metapredict,
    # with IUPred3 as fallback when AIUPred is unavailable.
    # AIUPred (Erdos & Dosztanyi 2024) is the recommended default; IUPred3
    # is retained as a REST-based fallback for environments without the
    # local CLI.
    meta_m = row.get("meta_disorder_motif_mean", np.nan)
    aiupr_m = row.get("aiupred_disorder_motif_mean", np.nan)
    iupr_m = row.get("iupred3_motif_mean", np.nan)
    disorder_candidates = [
        aiupr_m if pd.notna(aiupr_m) else 0.0,   # AIUPred (primary)
        meta_m  if pd.notna(meta_m)  else 0.0,   # metapredict
        iupr_m  if pd.notna(iupr_m)  else 0.0,   # IUPred3 (fallback)
    ]
    disorder_consensus = np.nanmax(disorder_candidates)
    S_disorder = clip01(disorder_consensus)
    # Disorder rescue: pLDDT in 60-85 AND any disorder predictor >= 0.5
    # (originally IUPred3-only; now applies to AIUPred too since it is the
    # primary predictor)
    plddt = row.get("mean_pLDDT_motif", np.nan)
    rescue_pred = aiupr_m if pd.notna(aiupr_m) else iupr_m
    if (pd.notna(plddt) and cal["plddt_lo"] <= plddt < cal["plddt_hi"] and
        pd.notna(rescue_pred) and rescue_pred >= 0.5):
        S_disorder = clip01(S_disorder + cal["iupred3_rescue_bonus"])
        row["iupred3_rescue_applied"] = True
    else:
        row["iupred3_rescue_applied"] = False

    # S_structure: prefer strict OR relaxed SSE pass. Per rule (b), α-helix is
    # tolerated (PY-NLS "prefers unstructured or α-helical" scaffold, β-strand
    # forbidden). So we do NOT penalize helical runs by themselves; strict/
    # relaxed pass status (beta count + core PY unstructured) is the gate.
    passes_strict = row.get("passes_sse_strict")
    passes_relaxed = row.get("passes_sse_relaxed")
    passes_borderline = row.get("passes_sse_borderline")
    if passes_strict is None and passes_relaxed is None and passes_borderline is None:
        S_structure = np.nan
    else:
        # Strict > relaxed > borderline > fail.
        if bool(passes_strict):
            S_structure = 1.0
        elif bool(passes_relaxed):
            S_structure = 0.7
        elif bool(passes_borderline):
            S_structure = 0.45
        else:
            S_structure = 0.15

    # S_exposure: mean RSA vs 25% target + PY-core RSA weighted extra
    rsa_mean = row.get("mean_RSA_motif_pct", np.nan)
    rsa_py = row.get("mean_RSA_PY_core_pct", np.nan)
    rsa_min = row.get("min_RSA_motif_pct", np.nan)
    if all(pd.isna(x) for x in (rsa_mean, rsa_py, rsa_min)):
        S_exposure = np.nan
    else:
        m1 = (rsa_mean / cal["rsa_mean_target"]) if pd.notna(rsa_mean) else 0.0
        m2 = (rsa_py / cal["rsa_mean_target"]) if pd.notna(rsa_py) else 0.0
        m3 = 1.0 if (pd.notna(rsa_min) and rsa_min >= cal["rsa_min_target"]) else 0.5
        S_exposure = clip01(0.4 * m1 + 0.5 * m2 + 0.1 * m3)

    # S_pLDDT: low pLDDT (<60) is good (=1); high (>85) is bad (=0.2); linear between
    if pd.isna(plddt):
        S_pLDDT = np.nan
    elif plddt < cal["plddt_lo"]:
        S_pLDDT = 1.0
    elif plddt >= cal["plddt_hi"]:
        S_pLDDT = 0.2
    else:
        # linear from 1.0 at plddt_lo to 0.2 at plddt_hi
        frac = (plddt - cal["plddt_lo"]) / (cal["plddt_hi"] - cal["plddt_lo"])
        S_pLDDT = 1.0 - 0.8 * frac

    # S_annotation: hard-veto if signal/transit/propeptide present.
    # In v6, the veto still sets S_annotation=0 (gating fused_score to 0),
    # but we also record the reason in exclusion_flag so the protein remains
    # visible in the output with its ungated_fused_score.
    S_annotation = 1.0
    exclusion_flag = ""
    _annot_map = {"Signal_peptide": "signal_peptide",
                  "Transit_peptide": "transit_peptide",
                  "Propeptide":      "propeptide"}
    for k, label in _annot_map.items():
        val = row.get(k, "")
        if has_annotation_value(val):
            S_annotation = 0.0
            exclusion_flag = label
            break

    # S_topology: computed elsewhere; carry it through if present
    S_topology = row.get("topo_multiplier", 1.0)
    if pd.isna(S_topology):
        S_topology = 1.0
    if S_topology == 0.0:
        exclusion_flag = (exclusion_flag + ";tm_topology").lstrip(";")

    # S_cnls: strong cNLS present -> soft downweight (protein may already have cNLS)
    cnls_p = row.get("cNLS_present")
    S_cnls = 0.85 if cnls_p else 1.0

    # S_af3 (opportunistic)
    ipsae_d0chn = row.get("af3_ipSAE_d0chn_mean_across_models", np.nan)
    if pd.isna(ipsae_d0chn):
        S_af3 = np.nan
    else:
        # 0.25 is roughly the top of the AF-214 distribution for positives
        S_af3 = clip01(ipsae_d0chn / 0.25)

    return {
        "S_sequence": S_sequence,
        "S_disorder": S_disorder,
        "S_structure": S_structure,
        "S_exposure": S_exposure,
        "S_pLDDT": S_pLDDT,
        "S_annotation": S_annotation,
        "S_topology": S_topology,
        "S_cnls": S_cnls,
        "S_af3": S_af3,
        "exclusion_flag": exclusion_flag,
        "iupred3_rescue_applied": row.get("iupred3_rescue_applied", False),
    }


def fuse_score(comp: dict, cal: dict) -> dict:
    """Fuse component scores into a single [0,1] fused score.

    Returns a dict with two keys:
      - ``fused``: the gated score (annotation/topology/cNLS multiplicative
        gates applied; 0 if any gate is 0).
      - ``ungated_fused``: the score *without* annotation/topology gates,
        so excluded proteins remain visible with their structural/disorder
        quality score.  cNLS soft multiplier is still applied.

    disorder/structure/exposure/pLDDT/sequence are weighted-averaged;
    cNLS is a soft multiplier; AF3 is a weighted rescue when available.
    """
    weights = cal["rule_weights"]
    parts = []
    weight_sum = 0.0
    for key, w in weights.items():
        v = comp.get(key, np.nan)
        if pd.notna(v):
            parts.append(w * v); weight_sum += w
    if weight_sum == 0:
        S_rules_core = np.nan
    else:
        S_rules_core = sum(parts) / weight_sum

    # --- Gated score (annotation + topology + cNLS) ---
    S_rules = S_rules_core
    if pd.notna(S_rules):
        S_rules = S_rules * comp.get("S_annotation", 1.0) * comp.get("S_topology", 1.0) * comp.get("S_cnls", 1.0)
    s_af3 = comp.get("S_af3", np.nan)
    if pd.notna(s_af3):
        w_af3 = cal["w_af3"]
        s_af3_gated = s_af3 * comp.get("S_annotation", 1.0) * comp.get("S_topology", 1.0)
        if pd.isna(S_rules):
            fused = s_af3_gated
        else:
            fused = w_af3 * s_af3_gated + (1.0 - w_af3) * S_rules
    else:
        fused = S_rules

    # --- Ungated score (cNLS only, no annotation/topology gates) ---
    S_rules_ungated = S_rules_core
    if pd.notna(S_rules_ungated):
        S_rules_ungated = S_rules_ungated * comp.get("S_cnls", 1.0)
    if pd.notna(s_af3):
        if pd.isna(S_rules_ungated):
            ungated_fused = s_af3
        else:
            ungated_fused = w_af3 * s_af3 + (1.0 - w_af3) * S_rules_ungated
    else:
        ungated_fused = S_rules_ungated

    return {
        "fused": float(fused) if pd.notna(fused) else np.nan,
        "ungated_fused": float(ungated_fused) if pd.notna(ungated_fused) else np.nan,
    }


def assign_tier(fused: float, comp: dict, cal: dict, extras: dict) -> int:
    """1 = high, 2 = moderate, 3 = weak, 4 = hard-vetoed / fail."""
    if pd.isna(fused):
        return 4
    if comp.get("S_annotation", 1.0) == 0.0:
        return 4
    if comp.get("S_topology", 1.0) == 0.0:
        return 4
    # Veto if strict, relaxed, AND borderline all fail (i.e. strong β content).
    ps = extras.get("passes_sse_strict")
    pr = extras.get("passes_sse_relaxed")
    pb = extras.get("passes_sse_borderline")
    if (ps is False) and (pr is False) and (pb is False):
        return 4
    cuts = cal["tier_cuts"]
    if fused >= cuts["T1"]: return 1
    if fused >= cuts["T2"]: return 2
    if fused >= cuts["T3"]: return 3
    return 4


# ==========================================================================
# MAIN PIPELINE
# ==========================================================================

def load_calibration(path: Path | None) -> dict:
    if path is None or not path.exists():
        return dict(DEFAULT_CALIBRATION)
    with open(path) as f:
        return json.load(f)


# ==========================================================================
# Per-protein data cache (v6)
# ==========================================================================

class PerProteinCache:
    """Disk-based per-protein JSON cache for network-fetched data.

    Caches UniProt annotations, cNLS Mapper results, AIUPred, IUPred3,
    metapredict, and AF model paths so that repeated runs (e.g. after code
    changes that don't affect network data) skip re-fetching.

    Cache validity is keyed on a SHA-256 hash of the protein sequence —
    if UniProt updates the sequence, the cache entry is automatically
    invalidated.
    """

    def __init__(self, cache_dir: Path | None, enabled: bool = True):
        self.enabled = enabled
        self.cache_dir = cache_dir
        if enabled and cache_dir is not None:
            cache_dir.mkdir(parents=True, exist_ok=True)

    @staticmethod
    def _seq_hash(seq: str) -> str:
        return hashlib.sha256(seq.encode()).hexdigest()[:16]

    def get(self, uid: str, seq: str) -> dict | None:
        """Return cached data for *uid* if the sequence hash matches, else None."""
        if not self.enabled or self.cache_dir is None:
            return None
        path = self.cache_dir / f"{uid}.json"
        if not path.exists():
            return None
        try:
            data = json.loads(path.read_text())
        except (json.JSONDecodeError, OSError):
            return None
        if data.get("seq_hash") != self._seq_hash(seq):
            return None  # sequence changed → cache invalid
        return data

    def save(self, uid: str, seq: str, data: dict):
        """Persist *data* for *uid*, adding the seq_hash for validation."""
        if not self.enabled or self.cache_dir is None:
            return
        data["seq_hash"] = self._seq_hash(seq)
        data["uniprot_id"] = uid
        path = self.cache_dir / f"{uid}.json"
        try:
            path.write_text(json.dumps(data, default=str, indent=2))
        except OSError:
            pass  # non-fatal — cache is an optimization


def run_pipeline(seqs: dict[str, str], args, logger: logging.Logger,
                 calibration: dict, af3_table_data: dict | None = None) -> pd.DataFrame:
    """Process each sequence: regex → structure → disorder → annot → topology
    → components → fuse → tier."""

    logger.info(f"Regex scan on {len(seqs)} sequences")
    all_hits: list[dict] = []
    per_protein_hits: dict[str, list[MotifHit]] = {}
    for uid, s in seqs.items():
        hits = scan_motifs(uid, s, strict_regex=args.strict_regex, calibration=calibration)
        per_protein_hits[uid] = hits
    total_hits = sum(len(v) for v in per_protein_hits.values())
    n_regex_pos = sum(1 for v in per_protein_hits.values() if v)
    logger.info(f"Regex hits: {total_hits} motifs across {n_regex_pos}/{len(seqs)} proteins")

    # Fetch UniProt annotations for candidates only (regex-positive)
    candidate_uids = [u for u, hs in per_protein_hits.items() if hs]

    # --- Per-protein cache setup (v6) ---
    cache_enabled = not getattr(args, 'no_cache', False)
    cache_dir = Path(args.cache_dir) if getattr(args, 'cache_dir', None) else (Path(args.out_dir) / "per_protein_cache")
    pp_cache = PerProteinCache(cache_dir if cache_enabled else None, enabled=cache_enabled)

    # Check cache for each candidate; split into cached vs fresh
    cached_data: dict[str, dict] = {}
    uids_to_fetch: list[str] = []
    for u in candidate_uids:
        c = pp_cache.get(u, seqs[u])
        if c is not None:
            cached_data[u] = c
        else:
            uids_to_fetch.append(u)
    if cached_data:
        logger.info(f"Per-protein cache: {len(cached_data)} hit(s), {len(uids_to_fetch)} to fetch")

    # --- UniProt annotations ---
    _empty_annot = {"TM_helix": "", "Signal_peptide": "",
                    "Transit_peptide": "", "Propeptide": "",
                    "gene_primary": "", "gene_names": "",
                    "gene_oln": "", "gene_orf": "", "protein_name": ""}
    uniprot_annot = {}
    for u in candidate_uids:
        if u in cached_data:
            uniprot_annot[u] = cached_data[u].get("uniprot_annot", dict(_empty_annot))
        else:
            uniprot_annot[u] = dict(_empty_annot)
    if uids_to_fetch and not args.no_fetch_uniprot_annot:
        fresh_annot = fetch_uniprot_annotations(uids_to_fetch, logger)
        uniprot_annot.update(fresh_annot)

    uids_need_pname = [
        u for u in candidate_uids
        if u in cached_data and not uniprot_annot.get(u, {}).get("protein_name", "")
    ]
    if uids_need_pname and not args.no_fetch_uniprot_annot:
        logger.info(f"Backfilling protein_name for {len(uids_need_pname)} cached entries")
        pname_annot = fetch_uniprot_annotations(uids_need_pname, logger)
        for u, a in pname_annot.items():
            if a.get("protein_name"):
                uniprot_annot[u]["protein_name"] = a["protein_name"]

    # --- AF PDB fetch ---
    af_cache = Path(args.out_dir) / "af_cache"
    af_paths = {}
    af_substitutes = {}  # uid -> substitute_uid ("" if none)
    stale_pdb_uids: list[str] = []  # cached proteins whose referenced PDB file is gone
    # Load cached AF paths
    for u in candidate_uids:
        if u in cached_data:
            af_info = cached_data[u].get("af_pdb", {})
            cached_pdb = af_info.get("path", "")
            if cached_pdb and Path(cached_pdb).exists():
                af_paths[u] = Path(cached_pdb)
            else:
                af_paths[u] = None
                if cached_pdb:
                    # The per-protein cache says this protein was already
                    # processed, but the AF structure file it points to no
                    # longer exists on disk (e.g. af_cache/ was cleared or
                    # moved independently of per_protein_cache/). Treat this
                    # as needing a re-fetch rather than silently scoring the
                    # protein with structure-derived features left blank.
                    stale_pdb_uids.append(u)
            af_substitutes[u] = af_info.get("substitute_uid", "")
        else:
            af_paths[u] = None
            af_substitutes[u] = ""
    if stale_pdb_uids:
        logger.warning(
            f"{len(stale_pdb_uids)} cached protein(s) have a missing AF PDB file on disk "
            f"(af_cache/ likely out of sync with per_protein_cache/); re-fetching structures "
            f"for them instead of silently skipping SSE/pLDDT/RSA"
        )
    # Fetch AF PDBs for uncached proteins and cached proteins with missing files.
    uids_need_af = [u for u in candidate_uids if af_paths.get(u) is None]
    if uids_need_af and not args.no_fetch_af:
        logger.info(f"Fetching AF PDBs for {len(uids_need_af)} candidates")
        with cfut.ThreadPoolExecutor(max_workers=args.threads) as ex:
            futs = {ex.submit(fetch_af_pdb, u, af_cache, logger, len(seqs[u])): u for u in uids_need_af}
            for f in tqdm(cfut.as_completed(futs), total=len(futs), desc="AF"):
                u = futs[f]
                try:
                    pdb_path, sub_uid = f.result()
                    af_paths[u] = pdb_path
                    af_substitutes[u] = sub_uid
                except Exception as e:
                    logger.warning(f"  {u}: AF fetch error {e}")
                    af_paths[u] = None
                    af_substitutes[u] = ""
    else:
        # Preserve an explicit missing value when structure fetching is disabled.
        for u in uids_need_af:
            af_paths[u] = None
            af_substitutes[u] = ""

    # --- cNLS Mapper (rate-limited; skip cached proteins) ---
    cnls_linker_mode = getattr(args, 'cnls_linker', 'entire')
    _empty_cnls = {"cNLS_present": None, "cNLS_mp_max_score": np.nan,
                   "cNLS_bp_max_score": np.nan, "cNLS_source": "skipped"}
    cnls_by_uid: dict[str, dict] = {}
    uids_need_cnls: list[str] = []
    for u in candidate_uids:
        if u in cached_data:
            cnls_by_uid[u] = cached_data[u].get("cnls", dict(_empty_cnls))
        else:
            cnls_by_uid[u] = dict(_empty_cnls)
            uids_need_cnls.append(u)
    if uids_need_cnls and not args.no_fetch_cnls:
        logger.info(f"cNLS Mapper on {len(uids_need_cnls)} candidates (~0.5 s/req, linker={cnls_linker_mode})")
        for u in tqdm(uids_need_cnls, desc="cNLS"):
            cnls_by_uid[u] = query_cnls_mapper(seqs[u], linker_mode=cnls_linker_mode)
            time.sleep(0.5)

    # --- AIUPred (local CLI; skip cached proteins) ---
    aiupred_results: dict[str, dict] = {}
    uids_need_aiupred: list[str] = []
    for u in candidate_uids:
        if u in cached_data and cached_data[u].get("aiupred"):
            # Convert cached lists back to numpy arrays
            cached_aiupred = cached_data[u]["aiupred"]
            aiupred_results[u] = {
                k: (np.array(v) if isinstance(v, list) else v)
                for k, v in cached_aiupred.items()
            }
        else:
            uids_need_aiupred.append(u)
    if uids_need_aiupred and not args.no_fetch_aiupred:
        aiupred_client = AIUPredClient()
        seqs_for_aiupred = {u: seqs[u] for u in uids_need_aiupred}
        logger.info(f"Running AIUPred locally on {len(uids_need_aiupred)} sequences")
        try:
            fresh_aiupred = aiupred_client.predict(seqs_for_aiupred)
            aiupred_results.update(fresh_aiupred)
            n_ok = sum(1 for v in fresh_aiupred.values() if v)
            logger.info(f"AIUPred completed: {n_ok}/{len(uids_need_aiupred)} succeeded")
        except Exception as e:
            logger.warning(f"AIUPred batch failed: {e}")

    # IUPred3 (fallback only — used when AIUPred is skipped or fails for a protein)
    iupred_client = IUPred3Client() if not args.no_fetch_iupred3 else None
    iupred_cache: dict[str, dict] = {}

    # Per-protein processing
    rows: list[dict] = []
    for uid in tqdm(candidate_uids, desc="Score"):
        seq = seqs[uid]
        hits = per_protein_hits[uid]
        is_cached = uid in cached_data

        # metapredict — use cache if available, else compute
        meta_full = None
        if is_cached and cached_data[uid].get("metapredict") is not None:
            # Reconstruct array from cached list
            meta_list = cached_data[uid]["metapredict"]
            if meta_list:
                meta_full = np.array(meta_list)
        elif not args.no_fetch_metapredict:
            try:
                import metapredict as meta
                meta_full = np.array(meta.predict_disorder(seq))
            except Exception:
                meta_full = None

        # AIUPred result for this protein (already computed in batch or from cache)
        aiupred_res = aiupred_results.get(uid, {})

        # IUPred3 fallback: only run if AIUPred was skipped or returned empty
        iupred_arr = None
        if is_cached and cached_data[uid].get("iupred3") is not None:
            iupred_cached = cached_data[uid]["iupred3"]
            if iupred_cached:
                iupred_arr = np.array(iupred_cached)
        elif iupred_client is not None and not aiupred_res:
            uid_ok_for_api = bool(re.match(
                r'^[OPQ][0-9][A-Z0-9]{3}[0-9]$|^[A-NR-Z][0-9]([A-Z][A-Z0-9]{2}[0-9]){1,2}$',
                uid))
            iupred_arr = iupred_client.predict(seq, uid=uid if uid_ok_for_api else None)
            time.sleep(1.5)  # gentle rate limit
            if iupred_arr is None:
                logger.debug(f"  {uid}: IUPred3 returned None")
            elif len(iupred_arr) != len(seq):
                logger.warning(f"  {uid}: IUPred3 length mismatch ({len(iupred_arr)} vs {len(seq)})")

        # Structure (per-protein)
        atoms = []
        sse_hec = None
        pdb_path = af_paths.get(uid)
        _af_sub_uid = af_substitutes.get(uid, "")
        if pdb_path is not None:
            atoms = parse_pdb_atoms(pdb_path)
            sse_raw = run_mkdssp(pdb_path)
            sse_hec = dssp_to_hec(sse_raw) if sse_raw else None
            # Mask low-pLDDT residues to coil ('C'): SSE assignments on
            # disordered regions (pLDDT < 60) are unreliable and should not
            # veto a motif via the strict/relaxed SSE gate.
            if sse_hec and atoms:
                plddt_by_res = {a["resnum"]: a["bfac"] for a in atoms if a["atomname"] == "CA"}
                for r, ss in list(sse_hec.items()):
                    p = plddt_by_res.get(r)
                    if p is not None and not np.isnan(p) and p < 60.0:
                        sse_hec[r] = 'C'

        # UniProt annot
        annot = uniprot_annot.get(uid, {"TM_helix": "", "Signal_peptide": "",
                                          "Transit_peptide": "", "Propeptide": "",
                                          "gene_primary": "", "gene_names": "",
                                          "gene_oln": "", "gene_orf": "",
                                          "protein_name": ""})
        tm_spans = parse_tm_spans(annot.get("TM_helix", ""))

        # cNLS
        cnls = cnls_by_uid.get(uid, {})

        # AF3 (opportunistic)
        af3_rec = {}
        if af3_table_data and uid.upper() in af3_table_data:
            af3_rec = af3_table_data[uid.upper()]
        elif args.af3_dir:
            files = find_af3_files(Path(args.af3_dir), uid)
            if files:
                af3_rec = parse_af3_metrics(uid, files)

        for h in hits:
            row = asdict(h)
            row["af_substitute_uid"] = _af_sub_uid
            # Add sequence features
            row["linker1_score"] = linker_length_score(h.linker1_len, calibration["linker1_target"], calibration["linker1_sigma"])
            row["linker2_score"] = linker_length_score(h.linker2_len, calibration["linker2_target"], calibration["linker2_sigma"])
            # metapredict
            if meta_full is not None:
                ms = meta_full[h.motif_start_idx-1:h.motif_end_idx]
                row["meta_disorder_motif_mean"] = float(ms.mean()) if len(ms) else np.nan
                row["meta_disorder_motif_min"]  = float(ms.min()) if len(ms) else np.nan
                row["meta_disorder_motif_max"]  = float(ms.max()) if len(ms) else np.nan
                row["meta_disorder_frac_ge_0p5_motif"] = float((ms >= 0.5).mean()) if len(ms) else np.nan
                row["meta_disorder_full_mean"]  = float(meta_full.mean())
                row["meta_source"] = "metapredict"
            else:
                row.update({"meta_disorder_motif_mean": np.nan, "meta_disorder_motif_min": np.nan,
                            "meta_disorder_motif_max": np.nan, "meta_disorder_frac_ge_0p5_motif": np.nan,
                            "meta_disorder_full_mean": np.nan, "meta_source": "skipped"})
            # AIUPred (default disorder predictor — local CLI)
            row.update(compute_aiupred(aiupred_res, h.motif_start_idx, h.motif_end_idx))
            # IUPred3 (fallback — only populated when AIUPred was unavailable)
            if iupred_arr is not None and len(iupred_arr) == len(seq):
                ms = iupred_arr[h.motif_start_idx-1:h.motif_end_idx]
                row["iupred3_motif_mean"] = float(ms.mean()) if len(ms) else np.nan
                row["iupred3_motif_min"]  = float(ms.min()) if len(ms) else np.nan
                row["iupred3_motif_max"]  = float(ms.max()) if len(ms) else np.nan
                row["iupred3_frac_ge_0p5_motif"] = float((ms >= 0.5).mean()) if len(ms) else np.nan
                row["iupred3_source"] = "iupred3_rest"
            else:
                row.update({"iupred3_motif_mean": np.nan, "iupred3_motif_min": np.nan,
                            "iupred3_motif_max": np.nan, "iupred3_frac_ge_0p5_motif": np.nan,
                            "iupred3_source": "skipped_or_unavailable"})
            # Structure
            sse_feat = compute_sse_features(sse_hec, h.motif_start_idx, h.motif_end_idx, h.seq_len) if sse_hec else compute_sse_features({}, h.motif_start_idx, h.motif_end_idx, h.seq_len)
            row.update(sse_feat)
            plddt_feat = compute_plddt_features(atoms, h.motif_start_idx, h.motif_end_idx) if atoms else {"mean_pLDDT_motif": np.nan, "min_pLDDT_motif": np.nan, "pLDDT_source": "NA"}
            row.update(plddt_feat)
            sasa_feat = compute_sasa_features(pdb_path, h.motif_start_idx, h.motif_end_idx) if pdb_path else {"mean_RSA_motif_pct": np.nan, "min_RSA_motif_pct": np.nan, "mean_RSA_PY_core_pct": np.nan, "sasa_source": "NA"}
            row.update(sasa_feat)
            # Annotation
            row.update(annot)
            # Topology
            topo = compute_topology_penalty(h.motif_start_idx, h.motif_end_idx, tm_spans,
                                            linker_threshold=calibration["topology_linker_threshold"],
                                            polytopic_min_loop=calibration["topology_polytopic_min_loop"],
                                            soft_multiplier=calibration["topology_soft_multiplier"])
            row["topo_status"] = topo["status"]
            row["topo_multiplier"] = topo["multiplier"]
            row["topo_n_tm"] = topo["n_tm"]
            row["topo_linker_dist"] = topo["linker_dist"]
            # cNLS
            row.update(cnls)
            # AF3
            for k in ("af3_ipSAE_d0chn_mean_across_models","af3_ipSAE_d0chn_max_across_models",
                       "af3_ipSAE_d0dom_mean_across_models","af3_ipSAE_d0dom_max_across_models",
                       "af3_ipTM_pae_mean_across_models","af3_ipTM_pae_max_across_models",
                       "af3_wc_iptm_mean","af3_wc_ptm_mean","af3_PY_wc_iptm_mean","af3_PY_wc_ptm_mean"):
                row[k] = af3_rec.get(k, np.nan)
            row["af3_source"] = af3_rec.get("af3_source", "absent")

            # Component scores
            comp = compute_component_scores(row, calibration)
            row.update(comp)
            # Fuse (returns both gated and ungated scores)
            fused_result = fuse_score(comp, calibration)
            row["fused_score"] = fused_result["fused"]
            row["ungated_fused_score"] = fused_result["ungated_fused"]
            row["exclusion_flag"] = comp.get("exclusion_flag", "")
            row["tier"] = assign_tier(row["fused_score"], comp, calibration, row)
            rows.append(row)

        # Save per-protein cache (network-fetched data only, not motif-dependent)
        if not is_cached:
            pdb_path_str = str(af_paths.get(uid)) if af_paths.get(uid) else ""
            cache_payload = {
                "uniprot_annot": uniprot_annot.get(uid, dict(_empty_annot)),
                "cnls": cnls_by_uid.get(uid, dict(_empty_cnls)),
                "aiupred": (
                    {k: (v.tolist() if isinstance(v, np.ndarray) else v)
                     for k, v in aiupred_res.items()}
                    if aiupred_res else {}
                ),
                "iupred3": iupred_arr.tolist() if iupred_arr is not None else None,
                "metapredict": meta_full.tolist() if meta_full is not None else None,
                "af_pdb": {"path": pdb_path_str,
                           "substitute_uid": af_substitutes.get(uid, "")},
            }
            pp_cache.save(uid, seq, cache_payload)

    df = pd.DataFrame(rows)
    return df


# ==========================================================================
# GENE-DEDUPLICATED OUTPUT
# ==========================================================================

def _build_by_gene_output(summary: pd.DataFrame) -> pd.DataFrame:
    """Build one-row-per-gene output from the per-protein summary.

    Gene key resolution (cross-organism):
      1. gene_primary (UniProt curated primary symbol)
      2. First token of gene_names (fallback for entries without gene_primary)
      3. UniProt accession (singleton — no gene annotation to merge on)

    Within each gene group, the protein with the highest fused_score is
    selected as the representative.  Ties are broken by sequence length
    (longer wins), then by accession (alphabetical) for determinism.

    Adds columns:
      gene_id              — the gene key used for grouping
      gene_symbol          — same as gene_id (separate column for clarity)
      n_proteins_per_gene  — count of UniProt proteins merged into this gene
      all_protein_ids      — semicolon-separated list of merged accessions
    """
    rows = []
    for _, r in summary.iterrows():
        annot = {"gene_primary": r.get("gene_primary", ""),
                 "gene_names": r.get("gene_names", ""),
                 "gene_oln": r.get("gene_oln", ""),
                 "gene_orf": r.get("gene_orf", "")}
        gkey = resolve_gene_key(annot)
        if not gkey:
            gkey = str(r.get("uniprot_id", ""))
        rows.append({
            "gene_key": gkey,
            "uniprot_id": r.get("uniprot_id", ""),
            "fused_score": r.get("fused_score", np.nan),
            "seq_len": r.get("seq_len", 0),
            "_row": r,
        })
    gene_df = pd.DataFrame(rows)
    # Sort by fused_score desc, then seq_len desc, then uniprot_id asc
    gene_df = gene_df.sort_values(
        ["fused_score", "seq_len", "uniprot_id"],
        ascending=[False, False, True],
    )
    # Group by gene_key, take first (highest-scored) as representative
    groups = gene_df.groupby("gene_key", sort=False)

    out_rows = []
    for gkey, grp in groups:
        rep = grp.iloc[0]
        r = rep["_row"]
        n_proteins = len(grp)
        all_ids = ";".join(sorted(grp["uniprot_id"].tolist()))
        out_row = r.to_dict()
        out_row["gene_id"] = gkey
        out_row["gene_symbol"] = gkey
        out_row["n_proteins_per_gene"] = n_proteins
        out_row["all_protein_ids"] = all_ids
        out_rows.append(out_row)

    by_gene = pd.DataFrame(out_rows)
    # Sort final output by fused_score descending
    by_gene = by_gene.sort_values("fused_score", ascending=False).reset_index(drop=True)
    return by_gene


# ==========================================================================
# ARGPARSE + ENTRY
# ==========================================================================

def parse_args(argv=None):
    p = argparse.ArgumentParser(description="pynls_v1 — plant PY-NLS predictor")
    # Input modes
    grp = p.add_mutually_exclusive_group(required=False)
    grp.add_argument("--fasta", help="Input FASTA file")
    grp.add_argument("--ids",   help="Text file with UniProt IDs (comma/newline)")
    grp.add_argument("--organism", help='Organism name, e.g. "Arabidopsis thaliana"')

    p.add_argument("--out-dir", default="./pynls_v1_out", help="Output directory")
    p.add_argument("--af3-dir", default=None, help="Directory with AF3 candidate-TRN1 outputs (opportunistic)")
    p.add_argument("--af3-table", default=None, help="Pre-aggregated AF3 metrics TSV (analyze_segments_byres_exact_v2.py schema)")

    p.add_argument("--strict-regex", action="store_true", help="Require linker1 <= 17 aa")
    p.add_argument("--topology-mode", default="plant", choices=["plant","strict","none"])

    p.add_argument("--calibration-json", default=None, help="Path to calibration JSON (overrides default)")

    p.add_argument("--emit-all-motifs", action="store_true",
                   help="Emit every regex hit (default: best-scoring per protein)")
    p.add_argument("--min-fused", type=float, default=0.0, help="Minimum fused score to emit")

    p.add_argument("--no-fetch-af", action="store_true", help="Skip AlphaFold monomer download")
    p.add_argument("--no-fetch-uniprot-annot", action="store_true", help="Skip UniProt annotation fetch")
    p.add_argument("--no-fetch-cnls", action="store_true", help="Skip cNLS Mapper")
    p.add_argument("--cnls-linker", default="entire", choices=["entire", "terminal"],
                   help="cNLS Mapper bipartite NLS search region: 'entire' (Entire region, default) "
                        "or 'terminal' (Within terminal 60-aa regions)")
    p.add_argument("--no-fetch-iupred3", action="store_true", help="Skip IUPred3 REST (fallback disorder predictor)")
    p.add_argument("--no-fetch-aiupred", action="store_true", help="Skip AIUPred local CLI (default disorder predictor)")
    p.add_argument("--no-fetch-metapredict", action="store_true", help="Skip metapredict")

    p.add_argument("--cache-dir", default=None,
                   help="Directory for per-protein JSON cache (default: {out_dir}/per_protein_cache). "
                        "Caches UniProt annotations, cNLS results, AIUPred, IUPred3, metapredict, "
                        "and AF model paths to speed up repeated runs.")
    p.add_argument("--no-cache", action="store_true",
                   help="Disable per-protein caching (always re-fetch everything)")

    p.add_argument("--threads", type=int, default=8, help="Parallel workers for network I/O")
    p.add_argument("--max-proteome", type=int, default=None,
                   help="Optional cap on number of sequences downloaded in organism mode (testing). "
                        "The pipeline auto-filters to regex-positive proteins before running "
                        "AIUPred/AF/DSSP/SASA/cNLS, so this is not needed for production runs.")
    p.add_argument("-v","--verbose", action="store_true")

    args = p.parse_args(argv)
    return args


def main(argv=None) -> int:
    args = parse_args(argv)

    if not (args.fasta or args.ids or args.organism):
        print("ERROR: one of --fasta / --ids / --organism is required")
        return 2

    out_dir = Path(args.out_dir); out_dir.mkdir(parents=True, exist_ok=True)
    logger = setup_logging(out_dir / "pynls_v1_run.log", args.verbose)
    logger.info(f"pynls_v1 v{__version__} starting")

    # Load calibration
    cal_path = Path(args.calibration_json) if args.calibration_json else (Path(__file__).parent / "pynls_v1_calibration.json")
    calibration = load_calibration(cal_path)
    if args.topology_mode == "strict":
        calibration["topology_linker_threshold"] = 100
    elif args.topology_mode == "none":
        calibration["topology_linker_threshold"] = 0  # sentinel; compute_topology_penalty won't gate
    logger.info(f"Calibration loaded: {list(calibration.keys())}")

    # Load AF3 table if provided
    af3_table_data = None
    if args.af3_table:
        af3_table_data = load_af3_from_table(Path(args.af3_table))
        logger.info(f"Loaded AF3 table for {len(af3_table_data)} proteins")

    # Resolve input
    seqs: dict[str, str] = {}
    if args.fasta:
        seqs = read_fasta(args.fasta)
    elif args.ids:
        ids = parse_id_list(args.ids)
        seqs = fetch_uniprot_sequences(ids, logger)
    elif args.organism:
        upid = resolve_organism_to_proteome(args.organism, logger)
        if upid:
            seqs = fetch_proteome_canonical(upid, logger, max_seqs=args.max_proteome)
    logger.info(f"Loaded {len(seqs)} sequences")
    if not seqs:
        logger.error("No sequences to process")
        return 1

    df = run_pipeline(seqs, args, logger, calibration, af3_table_data=af3_table_data)

    # Emit results
    if df.empty:
        logger.warning("No regex-consistent motifs found")
        return 0

    # Filter by min_fused
    if args.min_fused > 0:
        df = df[df["fused_score"] >= args.min_fused].copy()

    # --- Column reordering (v1): front-load key columns ---
    def _reorder_columns(frame: pd.DataFrame, gene_col: str = "gene_oln") -> pd.DataFrame:
        """Move key columns to the front for readability.
        
        For hits/summary: gene_oln (or fallback) as 2nd column.
        For by_gene: gene_id as 2nd column (passed via gene_col).
        """
        # Determine the gene column to use (fallback chain)
        gc = gene_col
        if gc not in frame.columns:
            for alt in ("gene_primary", "gene_names", "uniprot_id"):
                if alt in frame.columns:
                    gc = alt
                    break
        front = ["uniprot_id", gc, "protein_name", "fused_score", "tier", "exclusion_flag"]
        front = [c for c in front if c in frame.columns]
        remaining = [c for c in frame.columns if c not in front]
        return frame[front + remaining]

    # Emit all vs best-per-protein
    df = _reorder_columns(df)
    hits_path = out_dir / "pynls_v1_hits.tsv"
    df.to_csv(hits_path, sep="\t", index=False, float_format="%.6g")
    logger.info(f"Wrote {len(df)} rows to {hits_path.name}")

    # Best per protein
    if "fused_score" in df.columns and "uniprot_id" in df.columns:
        df["_rank"] = df.groupby("uniprot_id")["fused_score"].rank(method="first", ascending=False)
        best = df[df["_rank"] == 1].drop(columns=["_rank"]).copy()
        counts = df.groupby("uniprot_id").size().rename("n_motifs_found")
        tier_counts = df.groupby("uniprot_id")["tier"].apply(lambda s: dict(s.value_counts()))
        summary = best.merge(counts, left_on="uniprot_id", right_index=True)
        summary["tier_counts"] = summary["uniprot_id"].map(tier_counts).astype(str)
        summary = _reorder_columns(summary)
        summary_path = out_dir / "pynls_v1_summary.tsv"
        summary.to_csv(summary_path, sep="\t", index=False, float_format="%.6g")
        logger.info(f"Wrote {len(summary)} rows to {summary_path.name}")

        # Best per gene (one row per gene, highest fused_score representative)
        if "gene_primary" in summary.columns:
            by_gene = _build_by_gene_output(summary)
            by_gene = _reorder_columns(by_gene, gene_col="gene_id")
            by_gene_path = out_dir / "pynls_v1_by_gene.tsv"
            by_gene.to_csv(by_gene_path, sep="\t", index=False, float_format="%.6g")
            logger.info(f"Wrote {len(by_gene)} rows to {by_gene_path.name}")

    logger.info("pynls_v1 done")
    return 0


if __name__ == "__main__":
    sys.exit(main())
```
