## Supplementary Data 1 for "Sequence and structural rules define PY-NLS–dependent nuclear import by TRANSPORTIN 1 in plants": pynls_v1_README.md

# pynls_v1.py — Plant PY-NLS motif identification pipeline

**Version**: 1.0
**Purpose**: Identify plant proteins carrying novel basic PY-NLS (PY nuclear localization signal) motifs from FASTA, UniProt IDs, or organism-level proteomes.

## Motif definition

The PY-NLS is a bipartite/monopartite nuclear localization signal recognized
by Karyopherin-β2 (Transportin-1). This pipeline searches for the consensus:

```
[KR][KR] {3–20 aa linker1} [KRH] {1–5 aa linker2} P Y
```
implemented as:
```python
MOTIF_REGEX = re.compile(r'(?=([KR][KR].{3,20}[KRH].{1,5}PY))')
```

An overlapping (lookahead) match is used so nested candidates are found.

## Biology rules encoded (a–g)

| Rule | Encoded as |
|------|-----------|
| (a) K/R rich N-terminal + [KRH] before PY  | `MOTIF_REGEX` |
| (b) PY residues must be unstructured, motif preferentially unstructured or α-helical, β-strand forbidden | `S_structure` (strict / relaxed / borderline / fail) |
| (c) Solvent-accessible motif (RSA mean ≥25%, min ≥10%) | `S_exposure` |
| (d) Disordered / low-pLDDT context | `S_disorder` + `S_pLDDT` + IUPred3 rescue |
| (e) Linker geometry: linker1 near 17 aa (σ=5), linker2 near 3 aa (σ=2); no downstream hydrophobic accumulation (<3 in +1..+5) | `S_sequence` (0.5× penalty when triggered) |
| (f) No TM span crossing the motif; polytopic → loop ≥50 aa | `S_topology` (0 = hard veto; 0.5 = soft downweight) |
| (g) No secretory/organellar destination (SP, TP, propeptide) | `S_annotation` (0 = hard veto) |

## Score architecture

Each candidate motif receives 9 component scores in [0, 1]:

- `S_sequence`: linker geometry — Gaussian around linker1=17 (σ=5) and linker2=3 (σ=2), combined as 0.6·l1 + 0.4·l2, with 0.5× penalty when downstream hydrophobic accumulation ≥3
- `S_disorder`: max(metapredict, IUPred3) + rescue bonus (+0.15) when pLDDT is 60–85 and IUPred3 ≥0.5
- `S_structure`: 1.0 (strict pass), 0.7 (relaxed), 0.45 (borderline), 0.15 (fail)
- `S_exposure`: RSA mean weighted with RSA_PY_core (Tien 2013 max-ASA)
- `S_pLDDT`: 1.0 (pLDDT <60), linear down to 0.2 (pLDDT ≥85)
- `S_annotation`: 0 if signal-peptide / transit-peptide / propeptide present
- `S_topology`: 0 hard-veto in TM; 0.5 soft when loop too short
- `S_cnls`: 0.85 soft downweight if a strong classical NLS is already present
- `S_af3`: AlphaFold3 co-fold ipSAE_d0chn_mean / 0.25, opportunistic

**Fused score**:
```
S_rules       = Σ(w_i * S_i) / Σ(w_i)  over rule_weights keys, only valid components
S_rules_gated = S_rules × S_annotation × S_topology × S_cnls
S_af3_gated   = S_af3 × S_annotation × S_topology
fused         = w_af3 × S_af3_gated + (1 - w_af3) × S_rules_gated   [if AF3 present]
              = S_rules_gated                                        [otherwise]
```

Tier assignment:
- **T1** (high confidence): `fused ≥ 0.65`
- **T2** (moderate): `fused ≥ 0.45`
- **T3** (weak): `fused ≥ 0.25`
- **T4** (fail / vetoed): `fused < 0.25`, or `S_annotation == 0` (SP/TP/propeptide veto), or `S_topology == 0` (in TM), or SSE gate rejects (β-strand run in motif with no borderline pass)

## Input modes
```bash
# Mode 1: FASTA (protein IDs from headers)
python pynls_v1.py --fasta proteins.fa --out-dir results/

# Mode 2: UniProt accession list
python pynls_v1.py --ids ids.txt --out-dir results/

# Mode 3: Organism proteome (from UniProt Proteomes API)
python pynls_v1.py --organism "Arabidopsis thaliana" --out-dir results/
```

In organism mode the full proteome is downloaded from UniProt, but the expensive downstream steps (AlphaFold fetch, DSSP, FreeSASA, cNLS Mapper, AIUPred) run **only on proteins with a regex-positive PY-NLS motif** — typically ~1,500 of ~27,000 in Arabidopsis — so no manual cap is needed.
`--max-proteome N` is available as an optional download cap for testing.

## Optional data sources

- `--af3-table PATH`: AF3-co-fold-with-TRN1 per-model metrics (analyze_segments_byres_exact_v2.py schema)
- `--af3-dir PATH`: Directory of AF3 raw outputs
- `--calibration-json PATH`: Custom calibration weights (default: `pynls_v1_calibration.json`)

## Feature toggles

- `--no-fetch-af`: skip AlphaFold PDB fetch (no structure-based features)
- `--no-fetch-uniprot-annot`: skip TM/signal peptide extraction (no S_annotation)
- `--no-fetch-cnls`: skip cNLS Mapper (no S_cnls)
- `--no-fetch-aiupred`: skip AIUPred local CLI (default disorder predictor; falls back to IUPred3 REST)
- `--no-fetch-iupred3`: skip IUPred3 REST (fallback disorder predictor)
- `--no-fetch-metapredict`: skip metapredict (secondary disorder predictor)
- `--topology-mode {plant, strict, none}`: gate stringency
- `--cnls-linker {entire, terminal}`: cNLS Mapper linker search mode (default: entire)
- `--emit-all-motifs`: retain all motifs per protein in `_hits.tsv` (default: retain all)
- `--min-fused`: only emit motifs above this fused-score threshold
- `--cache-dir PATH`: per-protein cache directory (default: `{out_dir}/per_protein_cache`)
- `--no-cache`: disable per-protein caching

## Output files

### `pynls_v1_hits.tsv`
One row per motif. 
`uniprot_id`, `gene_oln`, `protein_name`, `fused_score`, `tier`, `exclusion_flag`

Followed by all feature columns:
- Motif geometry: `motif_seq`, `motif_start_idx`, `motif_end_idx`, `motif_len`, `linker1_len`, `linker2_len`, `n_hydrophobic_downstream`
- Structure: `sse_string_valid`, `helix_fraction_raw`, `beta_fraction_raw`, `passes_sse_{strict,relaxed,borderline}`, `mean_pLDDT_motif`, `mean_RSA_motif_pct`, `mean_RSA_PY_core_pct`
- Disorder: `aiupred_disorder_motif_mean`, `aiupred_binding_motif_mean`, `aiupred_linker_motif_mean`, `meta_disorder_motif_mean`, `iupred3_motif_mean`
- Annotation: `Signal_peptide`, `Transit_peptide`, `Propeptide`, `TM_helix`, `gene_primary`, `gene_names`
- Topology: `topo_status`, `topo_n_tm`, `topo_linker_dist`
- cNLS: `cNLS_present`, `cNLS_mp_max_score`, `cNLS_bp_max_score`
- AF3: `af3_ipSAE_d0chn_mean_across_models`, `af3_ipTM_pae_mean_across_models`, `af3_wc_iptm_mean`, `af3_PY_wc_iptm_mean`
- Component scores: `S_sequence`, `S_disorder`, `S_structure`, `S_exposure`, `S_pLDDT`, `S_annotation`, `S_topology`, `S_cnls`, `S_af3`

### `pynls_v1_summary.tsv`
One row per protein: best motif (highest fused_score) selected.
Same column order as hits.tsv (first 6: `uniprot_id`, `gene_oln`, `protein_name`, `fused_score`, `tier`, `exclusion_flag`).

### `pynls_v1_by_gene.tsv`
One row per **gene** (deduplicated from the per-protein summary).
**First 6 columns**: `uniprot_id`, `gene_id`, `protein_name`, `fused_score`, `tier`, `exclusion_flag`

Gene key resolution (cross-organism, not Arabidopsis-specific):
1. `gene_primary` — UniProt's curated primary gene symbol
2. First token of `gene_names` — fallback when `gene_primary` is empty
3. UniProt accession — singleton row when no gene annotation is available

Within each gene group, the protein with the highest `fused_score` is
selected as the representative. Ties are broken by sequence length (longer
wins), then by accession (alphabetical) for determinism.

Additional columns: `gene_id`, `gene_symbol`, `n_proteins_per_gene`, `all_protein_ids`

### `pynls_v1_run.log`
Full run log

### `per_protein_cache/`
Per-protein cached results (feature vectors, annotations, component scores).
Reused across runs to avoid redundant EBI/DSSP/SASA/cNLS/AIUPred calls.
Can be disabled with `--no-cache`.

### `af_cache/`
Cached AlphaFold PDBs (`{UID}_v6.pdb`); reused across runs.

## AlphaFold model retrieval (three-tier fallback)

1. **Primary**: direct fetch from AlphaFold DB using the UniProt accession
2. **UniRef100 substitute**: if the primary accession has no AF model, query
   UniRef100 for cluster members and try each for an AF model (same sequence,
   different accession that may have an AF structure)
3. **UniParc substitute**: if UniRef100 yields nothing (e.g., deleted TrEMBL
   entries), query UniParc for cross-referenced accessions and check each for
   an AF model

This recovers AF models for proteins whose original accession was removed from
UniProtKB but whose structure is still hosted at EBI under a different
accession.

## External dependencies

- **Python 3.10+**: numpy, pandas, requests, biopython, tqdm, metapredict
- **AIUPred** (local CLI): `pip install git+https://github.com/doszilab/AIUPred.git` — default disorder/binding/linker predictor (Erdos & Dosztanyi 2024)
- **mkdssp** binary: `/opt/conda/bin/mkdssp` v4.6.1
- **freesasa** Python bindings
- **REST APIs** (fallbacks only):
  - UniProt (`rest.uniprot.org`, `rest.uniprot.org/unisave`, `rest.uniprot.org/uniparc`)
  - AlphaFold DB (`alphafold.ebi.ac.uk`, v6 → v1 fallback)
  - IUPred3 (`iupred3.elte.hu/iupred3/{acc}.json`)
  - cNLS Mapper (`nls-mapper.iab.keio.ac.jp/cgi-bin/NLS_Mapper_form.cgi`)

## Calibration

| Linker1 length | Mean activity (×neg control) |
|----------------|------------------------------|
| 17             | 5.45                         |
| 16             | 4.11                         |
| 13             | 3.07                         |
| 12             | 4.11                         |
| 11             | 2.69                         |
| 6              | 1.88                         |
| 5              | 1.88                         |
| 4              | 2.42                         |

Pearson correlation between Gaussian linker1 score and experimental activity:
r = 0.89 (target=17, σ=5).

**v1 calibration** (`pynls_v1_calibration.json`):

| Parameter | v1 |
|-----------|------|
| linker1_sigma | 5 |
| S_sequence weight | 0.12 |
| S_disorder weight | 0.259 |
| S_structure weight | 0.311 |
| S_exposure weight | 0.104 |
| S_pLDDT weight | 0.207 |
| w_af3 | 0.100 |
| S_sequence formula | 0.6·l1 + 0.4·l2 |


## Design notes

- **S_sequence reform rationale**: The 0.2 baseline constant in the original
  formula had no biological justification — it was an ad-hoc floor that
  compressed the dynamic range between optimal and suboptimal linker lengths.
  Removing it and increasing the S_sequence weight allows linker geometry to
  properly influence motif selection without overriding structure-based
  signals (S_structure, S_pLDDT remain the dominant weights).
- **UniParc fallback**: Some TrEMBL entries are deleted ("Not part of a
  reference proteome") but their AlphaFold models remain hosted at EBI. UniParc
  indexes all historical cross-references, enabling recovery of AF models via
  alternative accessions.
- **DBREF sanitization**: mkdssp fails on PDB DBREF lines when a UniProt
  accession is >6 chars. Fixed by writing a sanitized PDB containing only
  coordinate records before invoking mkdssp.
- **pLDDT masking**: SSE assignments on residues with pLDDT <60 are unreliable;
  masked to coil before the SSE gate.
- **SSE borderline tier**: a single-residue β-strand call in an otherwise
  coily motif downgrades but does not veto; only extended β runs (≥2 residues)
  fail outright.
- **AIUPred as default disorder predictor**: uses the local AIUPred CLI
  (Erdos & Dosztanyi 2024) as primary, with IUPred3 REST as fallback.
  `S_disorder` uses `max(AIUPred_disorder, metapredict, IUPred3)`.
- **Per-protein caching**: all expensive per-protein computations (AF fetch,
  DSSP, SASA, cNLS, AIUPred, annotations) are cached to
  `{out_dir}/per_protein_cache/` and reused across runs. Cache hits return
  identical scores to fresh fetches.
