## Supplementary Data 1 for "Sequence and structural rules define PY-NLS–dependent nuclear import by TRANSPORTIN 1 in plants": pynls_v1_scoring_description.md

# PY-NLS identification pipeline — scoring description (v1)

## Motif definition

The PY-NLS consensus motif is defined by the regular expression:
```
[KR][KR].{3,20}[KRH].{1,5}PY
```
A lookahead-based regex (`(?=(...))`) is used so that overlapping/nested candidate motifs within the same protein are all identified. The motif consists of: (i) two basic residues (K or R) at the N-terminal end, (ii) a variable linker1 region of 3–20 amino acids, (iii) a single basic residue (K, R, or H), (iv) a short linker2 of 1–5 amino acids, and (v) a terminal proline–tyrosine (PY) dipeptide.

## Component scores

Each candidate motif is evaluated by seven component scores in the range [0, 1], each capturing an independent biological property:

### S_sequence — linker geometry

Scores how close the linker lengths are to experimentally determined optima. Linker1 (the region between the basic cluster and the [KRH] residue) is the strong determinant of nuclear import activity, as shown by a linker1-length mutagenesis study. Linker2 (between [KRH] and PY) is a secondary contributor.

Each linker length is scored by a Gaussian function:
```
l1_score = exp(−(linker1_len − 17)² / (2 × 5²))
l2_score = exp(−(linker2_len − 3)²  / (2 × 2²))
```
where linker1 target = 17 aa (σ = 5) and linker2 target = 3 aa (σ = 2), derived from experimental data. The combined score is:
```
S_sequence = clip(0.6 × l1_score + 0.4 × l2_score, 0, 1) × down_penalty
```
where `down_penalty = 0.5` if ≥3 hydrophobic residues occur in positions +1 to +5 downstream of the PY dipeptide (indicating a potential membrane-associated or aggregation-prone region incompatible with NLS function), and 1.0 otherwise. Linker1 is weighted 0.6 (vs linker2 at 0.4) because the experimental linker1 study demonstrated it is a strong determinant of import activity.

### S_disorder — intrinsic disorder

Consensus disorder propensity of the motif region, computed as the maximum of available predictors: AIUPred (Erdos & Dosztanyi 2024), metapredict, and IUPred3 (REST fallback). An IUPred3 rescue bonus of +0.15 is applied when the motif pLDDT is 60–85 (ambiguous structure) and IUPred3 ≥ 0.5, capturing cases where AlphaFold confidence is moderate but sequence-based disorder prediction supports flexibility.

### S_structure — secondary structure

Evaluates whether the motif's secondary structure is compatible with NLS function. DSSP is run on the AlphaFold model; residues with pLDDT < 60 are masked to coil (AlphaFold considers them effectively disordered). The motif is classified as:

- **Strict pass** (1.0): no β-strand residues in the motif
- **Relaxed pass** (0.7): ≤1 β-strand residue, remainder coil/helix
- **Borderline** (0.45): single isolated β-strand call (possible DSSP noise)
- **Fail** (0.15): ≥2 consecutive β-strand residues (rigid structure incompatible with NLS recognition)

### S_exposure — solvent accessibility

Relative solvent-accessible area (RSA) of the motif, computed by FreeSASA using Tien et al. (2013) maximum ASA reference values. The score rewards motifs with RSA mean ≥ 25% and RSA minimum ≥ 10%, indicating the motif is surface-exposed and accessible to the import receptor.

### S_pLDDT — AlphaFold confidence

Scores the AlphaFold pLDDT (predicted local distance difference test) confidence of the motif region. Low pLDDT (< 60) indicates intrinsic disorder and scores 1.0 (favorable for NLS). High pLDDT (≥ 85) indicates rigid structure and scores 0.2 (unfavorable). A linear interpolation connects these extremes:

```
S_pLDDT = 1.0 − 0.8 × clip((mean_pLDDT − 60) / (85 − 60), 0, 1)
```

### S_annotation — subcellular annotation veto

Hard veto (score = 0) if the protein has a signal peptide, transit peptide, or propeptide annotation (UniProt), indicating  a secretory or organellar destination incompatible with nuclear function. Otherwise 1.0.

### S_topology — transmembrane topology

Hard veto (score = 0) if the motif overlaps a transmembrane helix. Soft downweight (0.5) if the protein is polytopic and the motif is in a loop shorter than 50 aa (potential membrane-proximal constraint). Otherwise 1.0.

### S_cnls — classical NLS co-occurrence

Soft downweight (0.85) if cNLS Mapper detects a strong classical NLS (bipartite or monopartite) in the protein, indicating the PY-NLS may be redundant. Otherwise 1.0.

### S_af3 — AlphaFold3 co-fold interface (opportunistic)

When available, scores the interface pSAE (ipSAE_d0chn_mean) from AlphaFold3 co-folding the protein with Transportin-1 (TRN1). Normalized as ipSAE / 0.25, clipped to [0, 1]. Used as a light rescue signal, not a primary determinant.

## Fused score

The fused score combines all component scores into a single ranking metric:
```
S_rules       = Σ(w_i × S_i) / Σ(w_i)    over {S_sequence, S_disorder, S_structure, S_exposure, S_pLDDT}
S_rules_gated = S_rules × S_annotation × S_topology × S_cnls
S_af3_gated   = S_af3 × S_annotation × S_topology
fused         = w_af3 × S_af3_gated + (1 − w_af3) × S_rules_gated    [if AF3 present]
              = S_rules_gated                                           [otherwise]
```
The multiplicative gating by S_annotation, S_topology, and S_cnls ensures that hard vetoes (signal peptide, transmembrane overlap) zero out the score regardless of other favorable features.

## Tier assignment

| Tier | Criterion | Interpretation |
|------|-----------|----------------|
| T1 | fused ≥ 0.65 | High confidence PY-NLS |
| T2 | fused ≥ 0.45 | Moderate confidence |
| T3 | fused ≥ 0.25 | Weak / borderline |
| T4 | fused < 0.25, or S_annotation = 0, or S_topology = 0, or SSE gate fail | Rejected |

## Calibration weights

| Component | Weight |
|-----------|--------|
| S_sequence | 0.120 |
| S_disorder | 0.259 |
| S_structure | 0.311 |
| S_exposure | 0.104 |
| S_pLDDT | 0.207 |
| w_af3 | 0.100 |

S_structure (DSSP-based β-strand veto) is the strongest individual predictor, reflecting that PY-NLS motifs must reside in flexible, non-β-strand contexts. S_pLDDT is the second-highest weight, capturing the preference for intrinsically disordered regions. S_sequence, though lower in absolute weight, plays a critical role in motif selection within multi-motif proteins by favoring experimentally validated linker lengths.
